# Integrated single cell and unsupervised spatial transcriptomic analysis defines molecular anatomy of the human dorsolateral prefrontal cortex

**DOI:** 10.1101/2023.02.15.528722

**Authors:** Louise Huuki-Myers, Abby Spangler, Nick Eagles, Kelsey D. Montgomery, Sang Ho Kwon, Boyi Guo, Melissa Grant-Peters, Heena R. Divecha, Madhavi Tippani, Chaichontat Sriworarat, Annie B. Nguyen, Prashanthi Ravichandran, Matthew N. Tran, Arta Seyedian, PsychENCODE consortium, Thomas M. Hyde, Joel E. Kleinman, Alexis Battle, Stephanie C. Page, Mina Ryten, Stephanie C. Hicks, Keri Martinowich, Leonardo Collado-Torres, Kristen R. Maynard

## Abstract

The molecular organization of the human neocortex has been historically studied in the context of its histological layers. However, emerging spatial transcriptomic technologies have enabled unbiased identification of transcriptionally-defined spatial domains that move beyond classic cytoarchitecture. Here we used the Visium spatial gene expression platform to generate a data-driven molecular neuroanatomical atlas across the anterior-posterior axis of the human dorsolateral prefrontal cortex (DLPFC). Integration with paired single nucleus RNA-sequencing data revealed distinct cell type compositions and cell-cell interactions across spatial domains. Using PsychENCODE and publicly available data, we map the enrichment of cell types and genes associated with neuropsychiatric disorders to discrete spatial domains. Finally, we provide resources for the scientific community to explore these integrated spatial and single cell datasets at research.libd.org/spatialDLPFC/.

**Summary:** Generation of a molecular neuroanatomical map of the human prefrontal cortex reveals novel spatial domains and cell-cell interactions relevant for psychiatric disease.

## Introduction

The emergence of single cell and spatially-resolved transcriptomics has facilitated the generation of integrated molecular anatomical maps across a variety of tissues in both rodents and humans (1–4). Data-driven unsupervised approaches to identify spatial domains in these large datasets, particularly in the rodent brain, have refined our understanding of the spatial organization of tissues beyond classical cytoarchitectural boundaries (2), and computational models have helped reveal new insights into cellular diversification during development (5). However, efforts to generate spatially-resolved transcriptomics data in the human brain at the scale and size necessary to employ these approaches with sufficient statistical power, have lagged behind.

We previously characterized the spatial topography of gene expression in the human dorsolateral prefrontal cortex (DLPFC) by manually annotating the six histological layers and white matter of the neocortex in a small cohort of 3 neurotypical adult donors (6). While we identified robust layer-enriched gene expression with this approach, new computational tools for unsupervised clustering (7–11) and cell to cell communication (12) combined with the ability to expand the generation of molecular neuroanatomical maps across a larger donor pool has enabled data-driven identification of higher resolution spatial domains. Applying these data-driven approaches to larger-scale studies facilitates the ability to demarcate fine cortical sublayers, which currently lack molecular annotations. They also enable identification of novel, non-laminar spatial domains associated with anatomical or topographical features in the human brain, including vasculature.

Recent single nucleus RNA-sequencing (snRNA-seq) analyses, including those in companion PsychENCODE studies, are defining transcriptionally distinct DLPFC cell types and revealing cell type-specific changes associated with neurodevelopmental and neuropsychiatric disorders, such as schizophrenia (SCZ), autism spectrum disorder (ASD), major depressive disorder (MDD) and post-traumatic stress disorder (PTSD) (13–16). However, snRNA-seq data lack spatial context, which when retained during molecular profiling, can provide important insights into cell-cell communication and disease pathogenesis. To facilitate this type of analysis, we generated large-scale, unsupervised spatial transcriptomic molecular maps of the human DLPFC from neurotypical brain donors, which we integrated with snRNA-seq data across a variety of brain disorders. This spatially-resolved, molecular atlas of gene expression architecture in the human brain is provided as an interactive data resource for the scientific community to help reveal molecular mechanisms associated with psychiatric illness.

## Results

### Study Overview

Here we created the first large-scale, data-driven spatial map of gene expression at single cell resolution in the adult human DLPFC by applying integrated single cell and spatial transcriptomic approaches to identify novel spatial domains, define cell-cell communication (CCC) patterns, and perform spatial registration of cell types across brain disorders (**Fig. 1A-B**). Using the Visium spatial transcriptomics platform (10x Genomics), we measured spatial gene expression in fresh frozen postmortem human tissue blocks from 10 neurotypical adult donors (**Table S1**) in three positions spanning the rostral-caudal axis of the DLPFC (anterior [Ant], middle [Mid] and posterior [Post]) for a total of 30 tissue sections [n=10 per position] (**Fig. 1A**). In parallel, we performed snRNA-seq (10x Genomics 3’ gene expression) on a subset of the same DLPFC samples (n=1-2 blocks per donor) to generate matched snRNA-seq and spatial transcriptomic data for 19 tissue blocks (**Fig. 1A-C**). To preserve Layer (L)1, blocks were microdissected across sulci in the plane perpendicular to the pia that extended to the gray-white matter junction. The morphology of each tissue block was assessed with RNAscope multiplex single molecule fluorescent *in situ* hybridization (smFISH) using regional and laminar marker genes to ensure dissection consistency (**Fig. 1D**). Sample orientation was confirmed by expression of genes enriched in the gray matter (*SNAP25*), white matter (WM; [*MBP*]), and L5 (*PCP4*; **Fig. 1C, Fig S1, Fig S2, Fig S3**). For Visium, 4,866 (4.1%) spots with low library size were excluded (**Fig S4**), resulting in a total of 113,927 spots across 30 tissue blocks and 10 donors. Downstream analyses at the gene-level (**Fig S5**) and spot-level (**Fig S6**) were not impacted by tissue artifacts, including wrinkles, shears, and folds (Supplemental Methods: Evaluating the impact of histology artifacts on Visium H&E data). For snRNA-seq, 54,394 nuclei across 19 tissue blocks from 10 donors passed quality control and were included in the study (**Fig. 1C)**. Using these integrated datasets, we performed several analyses including unsupervised clustering, spot deconvolution, CCC analyses, and spatial registration of PsychENCODE single cell datasets (**Fig. 1B**).

**Figure 1.**
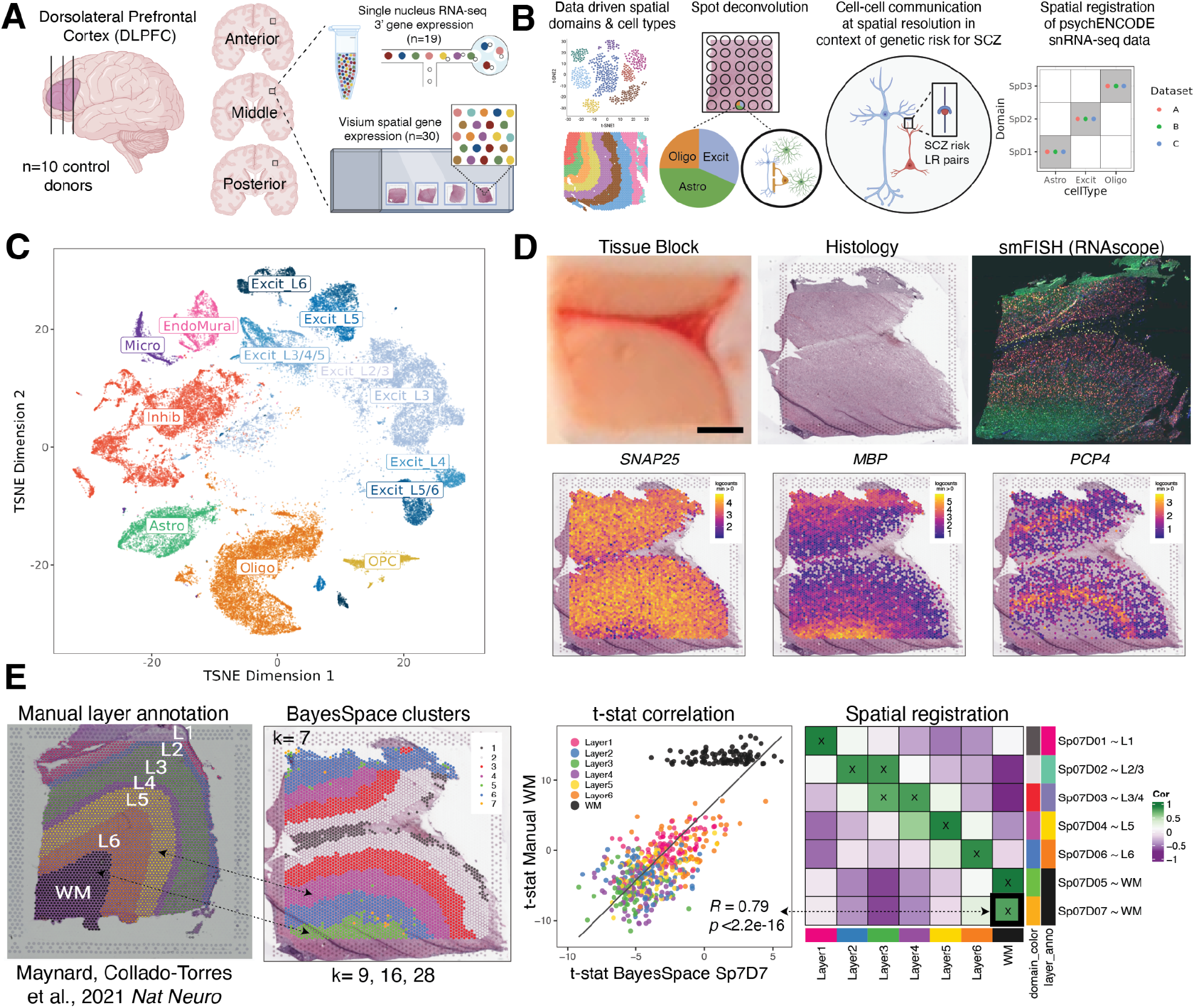
Study design to generate paired single nucleus RNA-sequencing (snRNA-seq) and spatially-resolved transcriptomic data across DLPFC. (**A**) DLPFC tissue blocks were dissected across the rostral-caudal axis from 10 adult neurotypical control postmortem human brains, including anterior (Ant), middle (Mid), and posterior (Post) positions (n=3 blocks per donor, n=30 blocks total). The same tissue blocks were used for snRNA-seq (10x Genomics 3’ gene expression assay, n=1-2 blocks per donor, n=19 samples) and spatial transcriptomics (10x Genomics Visium spatial gene expression assay, n=3 blocks per donor, n=30 samples). (**B**) Paired snRNA-seq and Visium data were used to identify data-driven spatial domains (SpDs) and cell types, perform spot deconvolution, conduct cell-cell communication analyses, and spatially register companion PsychENCODE snRNA-seq DLPFC data. (**C**) *t*-distributed stochastic neighbor embedding (t-SNE) summarizing layer resolution cell types identified by snRNA-seq. (**D**) Tissue block orientation and morphology was confirmed by hematoxylin and eosin (H&E) staining and single molecule fluorescent in situ hybridization (smFISH) with RNAscope (*SLC17A7* marking excitatory neurons in pink, *MBP* marking white matter (WM) in green, *RELN* marking layer (L)1 in yellow, and *NR4A2* marking L6 in orange). Scale bar is 2mm. Spotplots depicting log transformed normalized expression (logcounts) of *SNAP25, MBP*, and *PCP4* in the Visium data confirm the presence of gray matter, WM, and cortical layers, respectively (see also **Fig S1-Fig S3**). (**E**) Schematic of unsupervised SpD identification and registration using *BayesSpace* SpDs at *k*=7. Enrichment *t*-statistics computed on *BayesSpace* SpDs were correlated with manual histological layer annotations from (6) to map SpDs to known histological layers. The heatmap of correlation values summarizes the relationship between *BayesSpace* SpDs and classic histological layers. Higher confidence annotations (⍴ > 0.25, merge ratio = 0.1) are marked with an “X”.

### Identification of data-driven spatial domains at different resolutions across DLPFC

To select an unsupervised clustering method for robust identification of laminar spatial domains (SpDs) in Visium data, we benchmarked three algorithms, graph-based clustering, *SpaGGN* and *BayesSpace* (7,9,17,18), using our previously published DLPFC Visium data (12 sections from 3 donors; (6)) by comparing data-driven clustering accuracy against manual layer annotations (**Fig S8, Fig S9**). Among the algorithms tested, *BayesSpace* most accurately identified spatial domains (SpDs) consistent with the histological cortical layers. Therefore, we used this clustering algorithm to identify 7 unsupervised SpDs approximating the 6 cortical layers and WM. To relate unsupervised SpDs to the classic histological layers, we “pseudo-bulked” spots within each SpD across individual tissue sections to generate SpD-specific expression profiles and performed differential expression (DE) analysis to identify genes enriched in each SpD. Next, we performed “spatial registration” by correlating the enrichment statistics computed on *BayesSpace* SpDs with those from manually annotated cortical layers (6) to approximate the most strongly associated histological layer for a given *BayesSpace* SpD (**Fig. 1E**). We denote the association of a specific spatial domain (SpD) at cluster resolution *k* to a classic histological layer using the term Sp_k_D_d_~*L*, where *L* refers to the histological layer most strongly associated to domain *d* following cluster registration at resolution *k*. For example, spatial domain 7 at cluster resolution *k*=7 (Sp_7_D_7_) was mostly strongly associated with white matter (**Fig. 1E**) and is annotated as Sp_7_D_7_~WM. We found that *k*=7 was not sufficient to fully separate histological layers, especially superficial L2-L4, suggesting the presence of higher resolution data-driven SpDs and highlighting the challenges of manually annotating spatial domains (6).

We next evaluated how increasing cluster resolution (*k*) influenced the identification and cluster registration of unsupervised SpDs (**Fig. 2**). As expected, clustering at *k*=2 reliably separated white and gray matter (**Fig S10, Fig S11**). Next, we evaluated three clustering resolutions: a “broad” resolution *k*=9, a data-driven “fine” resolution *k*=16 (**Fig S12**), and a “super-fine” resolution *k*=28. Hereafter, we refer to these SpDs as Sp_9_D, Sp_16_D, and Sp_28_D, respectively. Broad clustering most accurately recapitulated the classic histological layers with clear separation of Sp_9_Ds enriched in genes expressed in L1-6 and WM (**Fig. 2A-B, Fig S13**). At fine clustering resolution, SpDs were largely laminar with two or more Sp_16_Ds registering to a given histological layer, suggesting the presence of molecularly-defined sublayers (**Fig. 2A-B, Fig S14**). At “super-fine” resolution, many SpDs lacked a laminar structure, but spots belonging to Sp_28_Ds frequently mapped back to a single broad or fine resolution spatial domain (**Fig S15, Fig S16**). To more deeply evaluate the unsupervised architecture of cortical layers in the human DLPFC, we focused on broad and fine SpDs based on the robust laminar features of these domains.

**Figure 2.**
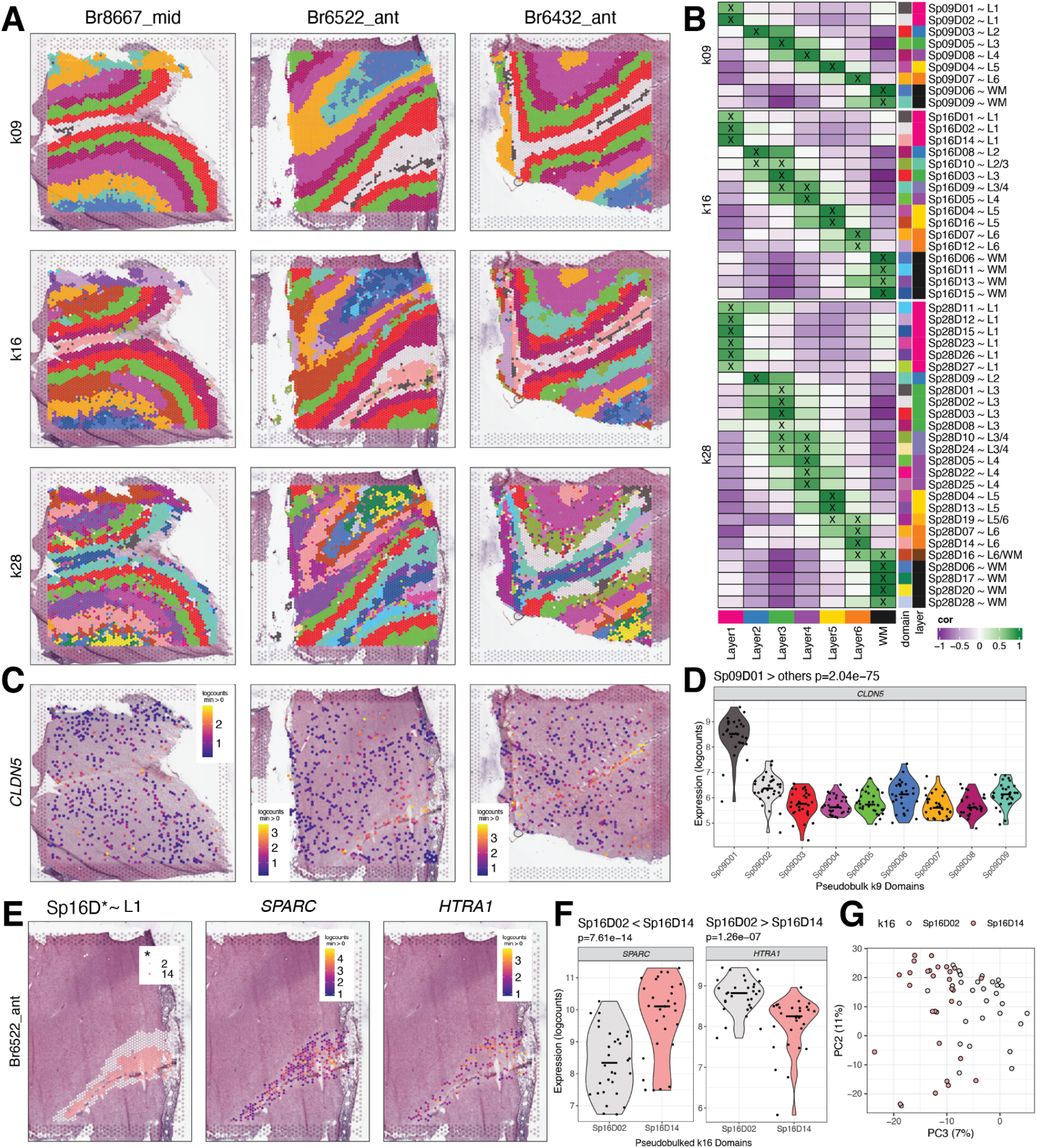
Unsupervised clustering at different resolutions identifies novel spatial domains (SpDs) and defines molecular anatomy of DLPFC. (**A**) *BayesSpace* clustering at *k*=9, 16, and 28 (broad, fine, and super-fine resolution, respectively, which we refer to as Sp_k_D_d_ for domain *d* from SpDs at *k* resolution) for three representative DLPFC tissue sections (Br8667_mid, Br6522_ant, Br6432_ant). (**B**) Heatmap of spatial registration with manually annotated histological layers from (6). *BayesSpace* identifies laminar SpDs at increasing *k* with the majority of Sp_k_Ds correlating with one or more histological layer(s). SpDs were assigned layer-level annotations following spatial registration to histological layers. Annotations with high confidence (⍴ > 0.25, merge ratio = 0.1) are marked with an “X”, and this histological layer association is denoted for a given Sp_k_D_d_ by adding “~*L*,” where *L* is the most strongly correlated histological layer (or WM). See also **Fig S10-Fig S16**. (**C**) Spotplots depicting expression of *CLDN5* in vasculature domain 1 at *k*=9 resolution (Sp_9_D_1_). (**D**) Boxplot confirming enrichment of *CLDN5* in Sp_9_D_1_ compared to other Sp_9_Ds across 30 tissue sections. (**E**) Spotplots of representative section Br6522_ant showing identification of molecularly-defined sublayers for histological L1 at k=16 (Sp_16_D_2_ and Sp_16_D_14_) and enrichment of *HTRA1* and *SPARC*, respectively. (**F**) Boxplots quantifying enrichment of *SPARC* and *HTRA1* in Sp_16_D_14_ and Sp_16_D_2_, respectively, across 30 tissue sections. (**G**) PCA plot showing separation of Sp_16_D_2_ and Sp_16_D_14_ supporting identification of molecularly distinct SpDs.

### Enrichment of differentially expressed genes in unsupervised spatial domains

To identify the molecular signatures of SpDs within each clustering resolution, we next performed differential expression (DE) analyzes with linear mixed-effects modeling using the Sp_9_D-or Sp_16_D-specific expression profiles. As previously described, we employed three different statistical models: ANOVA model, enrichment model, and pairwise model (6). As expected, all three models confirmed that unsupervised clustering at broad or fine resolution identified biologically meaningful SpDs with significant DE of genes across the laminar architecture of the cortex (5,931 FDR<5% unique enriched genes in at least one Sp_9_D, Supplemental Methods: Layer-level data processing and differential expression modeling). While we did identify 512 unique genes that were differentially enriched across the anterior-posterior axis of the DLPFC, SpD had a much stronger effect on gene expression compared to anatomical position (anterior, middle, posterior, **Fig S17**).

Analysis of DEGs identified using the enrichment model allowed for characterization of novel data-driven SpDs, such as Sp_9_D_1_ and Sp_16_D_1_ (**Fig. 2B**). At both *k*=9 and *k*=16, domain 1 is adjacent to histological L1 and enriched for genes associated with blood vessels and brain vasculature, such as *CLDN5* (p=2.04e-75; **Fig. 2C-D**). Due to its thinness, this vascular-rich meninges layer was not manually annotated in our previous study (6), demonstrating the utility of unsupervised approaches to robustly identify biologically meaningful SpDs. Data-driven clustering at fine resolution also revealed molecularly-defined sublayers, including two adjacent laminar spatial subdomains (Sp_16_D_14_~L1 and Sp_16_D_2_~L1; **Fig. 2E**) enriched in L1 marker genes, including *RELN* (p=6.98e-17, p=3.19e-19 respectively) and *AQP4* (p=9.37e-21, p=9.43e-12 respectively). Pairwise tests across these L1-related subdomains highlighted differential expression of *SPARC* (enriched in Sp_16_D_14_~L1, p=7.61e-14) and *HTRA1* (enriched in Sp_16_D_2_~L1, p=1.26e-07), and principal component analysis (PCA) further confirmed the unique nature of these molecularly-defined sublayers (**Fig. 2E-G**). We also identified multiple SpDs associated with histological L4 (Sp_16_D_5_-L4 and Sp_16_D_9_-L4), L5 (Sp_16_D_4_-L5 and Sp_16_D_16_-L5), and L6 (Sp_16_D_7_-L6 and Sp_16_D_12_-L6; **Fig. 2B**). Taken together, these analyses validate the biological relevance of data-driven SpDs and reveal novel molecular neuroanatomy across the laminar architecture of the adult human DLPFC.

### Identification of molecularly and spatially distinct neuronal populations across cortical layers

To add single cell resolution to our molecular maps, we performed snRNA-seq on a subset of the same tissue blocks used for Visium (**Fig. 1**). Following assessment of quality control metrics (**Fig S18, Fig S19**), we performed batch correction (**Fig S20, Fig S21, Fig S22**) and data-driven clustering to generate 29 fine-resolution clusters across 7 broad cell types represented throughout the anterior-posterior DLPFC axis (**Fig. 1A, Fig S23**). To add anatomical context to snRNA-seq clusters, we spatially registered all clusters to the histological layers using manually annotated Visium data in (6) (**Fig. 3B**). Given that molecularly-defined excitatory neuron populations in the cortex have distinct laminar identities (19), we systematically assigned a histological layer to excitatory neuron clusters (**Table S2**), resulting in the identification of 13 “layer-resolution” clusters with distinct marker genes (**Fig. 3B-D, Fig S24**). At both fine-resolution and layer-resolution, our clusters strongly correlated with those derived from the reference-based mapping tool, *Azimuth* (https://azimuth.hubmapconsortium.org) (20,21) (**Fig S25**).

**Figure 3.**
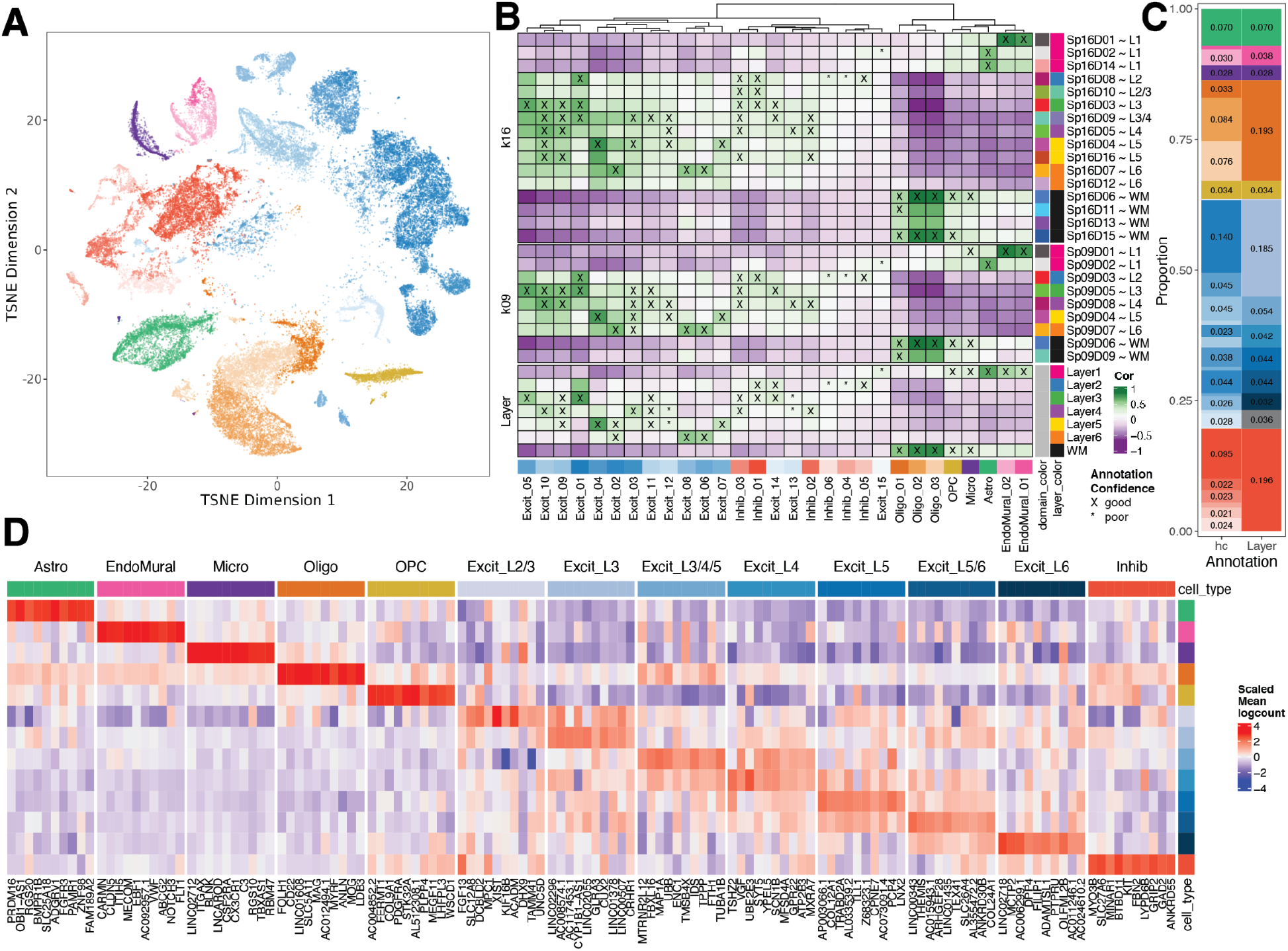
Spatial registration of fine resolution snRNA-seq clusters defines laminar cell types. (**A**) t-distributed stochastic neighbor embedding (t-SNE) plot of 56,447 nuclei across 29 cell type-annotated fine resolution hierarchical clusters (hc; related to **Fig S23A)**. (**B**) Spatial registration heatmap showing correlation between snRNA-seq hierarchical clusters (hc) and manually annotated histological layers from (6) as well as unsupervised *BayesSpace* clusters at *k*=9 and 16 (Sp_9_Ds and Sp_16_Ds). Hierarchical clusters for excitatory neurons (Excit) were assigned layer-level annotations following spatial registration to histological layers (⍴ > 0.25, merge ratio = 0.25). For Sp_9_Ds and Sp_16_Ds, annotations with good confidence (⍴ > 0.25, merge ratio = 0.1) are marked with “X” and poor confidence are marked with “*”. (**C**) Summary barplot of cell type composition for hc and layer level resolutions (related to **Fig S23B** & **Fig S24**) (**D**) Heatmap of the scaled mean pseudo-bulked logcounts for the top 10 marker genes identified for each cell type at layer-level resolution.

To gain further insight into the relationship between our snRNA-seq clusters and Visium unsupervised SpDs, we performed spatial registration of fine resolution snRNA-seq clusters with Visium SpDs at *k*=9 and *k*=16 (**Fig. 3B**). Using this approach, we refined the spatial positioning of our 29 fine resolution snRNA-seq clusters and validated the laminar associations of broad and fine unsupervised SpDs. For example, we showed that snRNA-seq excitatory neuron clusters Excit_06 and Excit_08, which spatially registered to histological L6, were also highly correlated with Visium Sp_9_D_7_, a domain enriched for L6-associated genes (**Fig. 2B**). These snRNA-seq clusters were also highly correlated with a single spatial domain (Sp_16_D_7_), thereby further refining their anatomical position to upper L6. Inhibitory GABAergic populations were also assigned to specific spatial locations. For example, Inhib_05 uniquely registered to histological L2, which was confirmed with strong correlations to Sp_9_D_3_ and Sp_16_D_8_ showing enrichment for L2-associated genes (**Fig. 2B**). We also showed registration of snRNA-seq endothelial cell populations to vascular spatial domains (Sp_9_D_1_ and Sp_16_D_1_) enriched in *HBA1* and *CLDN5*.

### Defining cell type composition of unsupervised spatial domains using spot deconvolution

Given that individual Visium spots in the human DLPFC contain an average of 3 cells per spot (6), we used our paired snRNA-seq data to perform cellular deconvolution of Visium spots to better understand the cell type composition of unsupervised SpDs. First, we benchmarked 3 spot-level deconvolution algorithms, *SPOTlight, Tangram*, and *Cell2location* (8,22,23), using a gold standard reference dataset acquired with the Visium Spatial Proteogenomics (Visium-SPG) assay. Visium-SPG replaces H&E histology with immunofluorescence staining, enabling us to label and quantify 4 broad cell types across the DLPFC, including NeuN (neurons), OLIG2 (oligodendrocytes), GFAP (astrocytes), and TMEM119 (microglia) (**Fig. 4A, Fig S26**). After verifying marker genes for each snRNA-seq cluster (**Fig S27**) and confirming the utility of these genes for spot deconvolution (**Fig. 4B, Fig S28, Fig S29, Fig S30**), we applied *SPOTlight, Tangram*, and *Cell2location* to our Visium-SPG data and calculated the predicted cell type counts per spot at broad and fine resolution (**Fig. 4B, Fig S31**).

**Figure 4.**
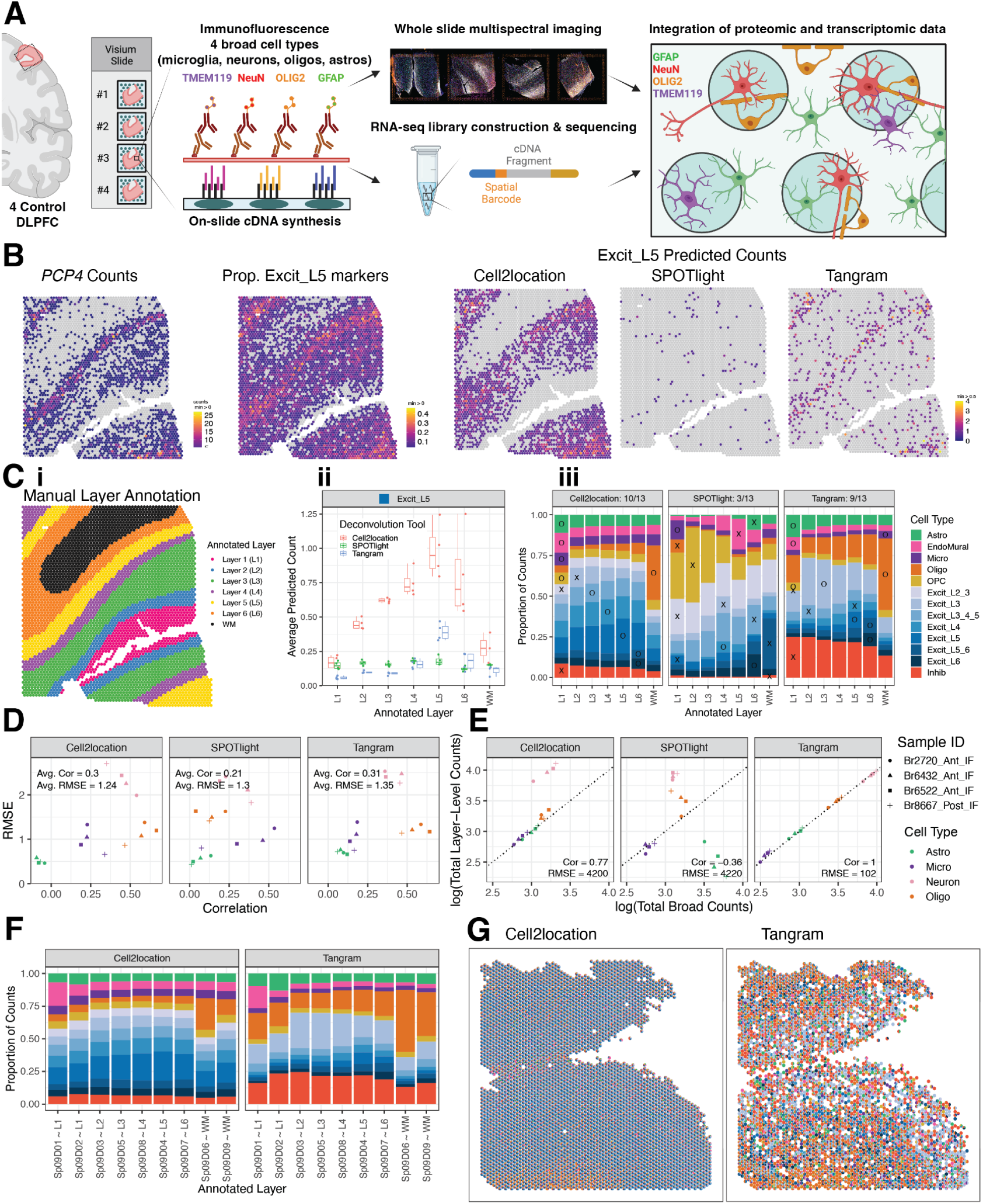
Integration of snRNA-seq and Visium data to benchmark spot deconvolution algorithms and define cellular composition across spatial domains. (**A**) Schematic of the Visium-SPG protocol. (**B**) For Br6522_Ant_IF, counts for L5 marker gene *PCP4* are compared to the proportion of Excit_L5 marker genes with nonzero expression as well as the counts of Excit_L5 cells as predicted by the 3 evaluated deconvolution algorithms. (**C**) Example of manually annotated layer assignments for Br6522_Ant_IF (i), which are used to benchmark predicted cell type composition across layers. Using Excit_L5 as an example, predicted Excit_L5 counts for each method are averaged across all spots within each annotated layer for each tissue section (ii). These data are summarized across layers and tissue sections for the 13 cell types using a bar plot (iii). An “X” or “O” is placed on the layer with maximal proportion; an “O” is placed for a “correct” match for the given cell type, and an “X” is placed otherwise. For example, *Tangram* correctly predicts the maximal proportion of Excit_L5 cells in L5 annotated spots, leading to the placement of an “O” for Excit_L5. The “O”s are tallied for each method to generate a summary score in each facet’s title (i.e. 9 of 13 cell types were correctly predicted to the expected layer using *Tangram*). (**D**) Predicted counts for a given method, section, and cell type are compared against the corresponding CART predictions by computing the Pearson correlation and RMSE, forming a single point in the scatterplot. Each of these values is then averaged to generate a single correlation and RMSE value for each method, indicated in the top left inside each plot facet. (**E**) Section-wide counts for each cell type are compared between broad and layer-level resolutions, collapsed onto the cell-type resolution used by the CART, where values theoretically should precisely match. (**F**) The predicted proportion of cells in each Sp_9_D, deconvoluted by *Cell2location* and *Tangram*, are averaged across all Visium samples (n=30). (**G**) Cell composition of each Visium spot for Br8667_mid, deconvoluted by *Cell2location* and *Tangram*, revealing differences in cell composition prediction.

To quantify algorithm performance, we took 2 complementary approaches: 1) evaluating the localization of laminar cell types to their expected cortical layer (**Fig. 4C, Fig S32, Fig S33**) and 2) comparing predicted cell type counts to those obtained from immunofluorescent images using a classification and regression tree (CART) strategy to categorize nuclei into the 4 immunolabeled cell types (**Fig. 4D, Fig S34, Fig S35, Fig S36, Fig S37**). Using the first approach, we found that *Tangram* and *Cell2location* performed best across all cell types, but *SPOTlight* failed to accurately map excitatory neuron subtypes to the correct layer (**Fig. 4B**).

Using the second approach, we found that the predicted counts from *Tangram* and *Cell2location* also had the highest correlation to CART-calculated counts (**Fig. 4D**) and *Tangram* showed the most consistent performance at both broad and layer level resolution across all cell types and samples (**Fig. 4E**). Finally, we applied *Tangram* and *Cell2location* to our H&E Visium dataset to predict the cellular composition of SpDs across the anterior-posterior axis of the DLPFC (**Fig. 4F-G, Fig S38, Fig S39**) and found that while *Tangram* and *Cell2location* showed differences in the predicted cell counts per spot (**Fig. 4F**), for both tools, the predicted cellular composition of SpDs was consistent across samples at both broad (*k*=9) and fine (*k*=16) resolution regardless of DLPFC position (i.e, anterior, middle, posterior).

### Spatial mapping of ligand-receptor (LR) interactions associated with schizophrenia (SCZ)

To add clinical relevance to this integrated DLPFC dataset, we next sought to identify interacting cell types and spatially map ligand receptor (LR) interactions associated with neuropsychiatric disorders. We focused on genetic risk for schizophrenia (SCZ) because receptors occur more frequently in SCZ risk genes than would be expected of a brain-expressed gene list of this size (p<0.0001, **Fig S38A**). First, we identified interacting cell types using cell-cell communication (CCC) analysis (12), which uses a data-driven approach to predict crosstalk between sender and receiver cells based on known LR interactions (**Fig. 5A, Table S3**). In parallel, using the OpenTargets and Omnipath databases (24,25), we identified 834 LR pairs (**Table S4**) associated with genetic risk for SCZ and prioritized 18 inter- and intra-cellular interactions where both counterparts showed disease association, including 9 interactions involving the protein tyrosine kinase, *FYN* (**Fig. 5A, Fig S40**). A consensus LR pair was identified between these complementary data-driven and clinical risk-driven approaches: the membrane-bound ligand ephrin A5 (*EFNA5*) and its receptor ephrin type-A receptor 5 (*EPHA5*). As part of this signaling cascade, we also evaluated the intracellular interaction between *EFNA5* and *FYN*, which was one of the 18 SCZ-associated LR pairs (**Fig. 5A**).

**Figure 5.**
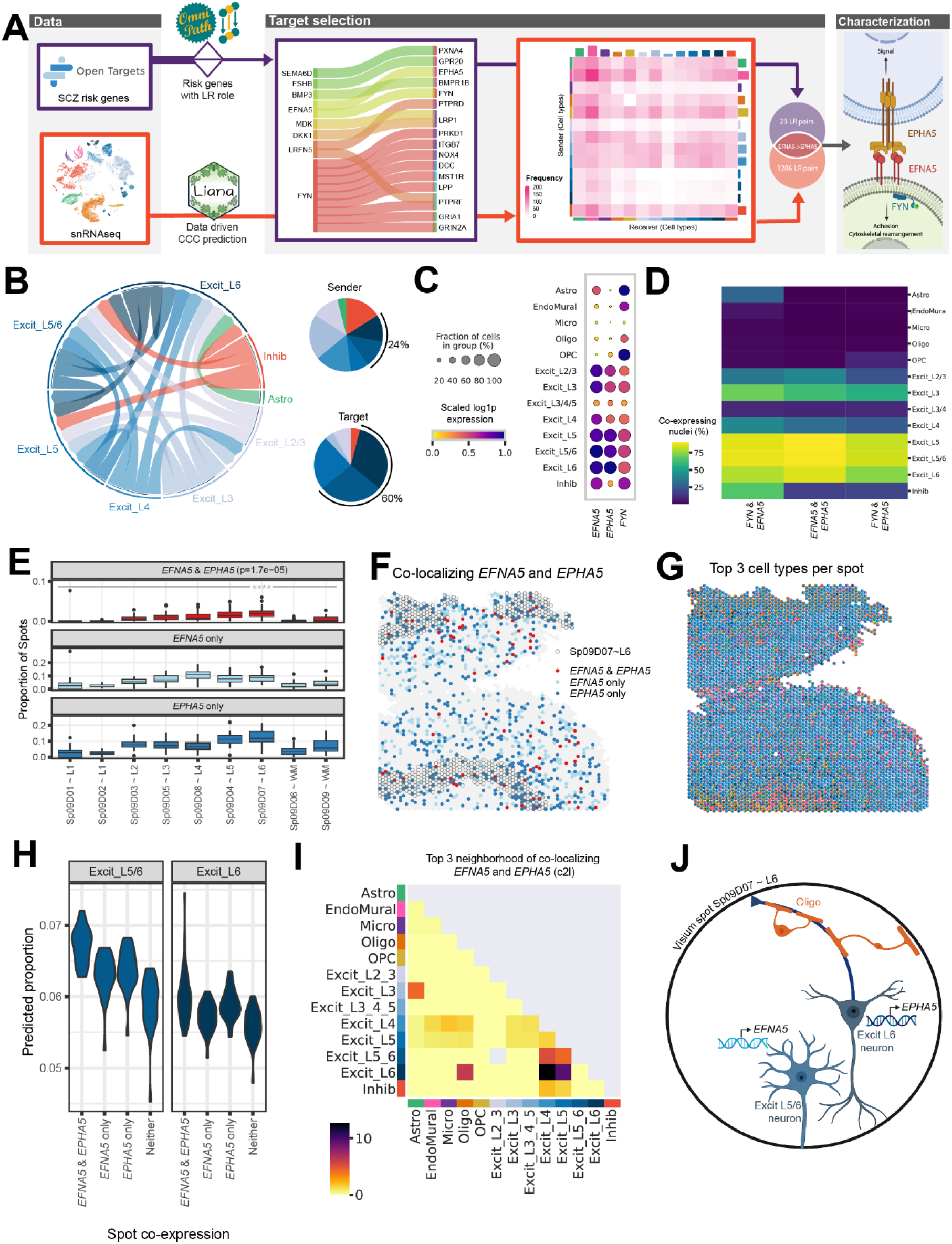
Integrative analysis of snRNA-seq and Visium data identifies ligand-receptor (LR) interactions associated with schizophrenia (SCZ). (**A**) The LR interaction between membrane-bound ligand ephrin A5 (*EFNA5*) and ephrin type-A receptor 5 (*EPHA5*) is a consensus target identified in both data-driven (**Table S3**) and clinical risk-driven LR (**Table S4**) analyses. Notably, this interaction also requires an intracellular interaction between *EFNA5* and protein tyrosine kinase (*FYN*), which was also identified among clinical risk targets. (**B**) Cell-cell communication analysis predicts the sender/receiver cross-talk pattern of *EFNA5*-*EPHA5* between layer-level cell types visualized in a circular plot. (**C-D**) Downstream analysis of snRNA-seq data characterizes *FYN-EFNA5*-*EPHA5* signaling pathway, showing these genes are highly enriched (**C**) and co-expressed (**D**) in excitatory neuron populations. (**E**) Across all 30 tissue sections, *EFNA5* and *EPHA5* are co-expressed in a higher proportion of spots in Sp_9_D_7_ compared to other Sp_9_Ds. (**F**) Spotplot of *EFNA5* and *EPHA5* co-expression in Br8667_mid. (**G**) Spotplot with spot-level pie charts for Br8667_mid showing the top 3 dominant cell types in each Visium spot predicted by *Cell2location (c2l)*. (**H**) Visium spots co-expressing *EFNA5* and *EPHA5* have higher proportions of predicted Excit_L5/6 neurons (p=1.8e-12) and Excit_L6 (p=3.9e-4) compared to non-coexpressing spots, consistent with snRNA-seq specificity analyses (**Fig S40**). Few other cell types show this relationship (**Fig S41**). Complementary analyses of *EFNA5* and *FYN* co-expression are shown in **Fig S40**. (**I**) Spatial network analysis of all 30 tissue sections, using top 3 dominant *c2l* cell types in each spot (exemplified in **G** with Br8667_mid), confirms *EFNA5* and *EPHA5* co-expression occurs frequently in spots containing Excit_L6 neurons. Complementary analyses using top 6 dominant *c2l* cell types as well as *Tangram* predictions are reported in **Fig S40**. (**J**) Schematic of a Visium spot depicting *EFNA5*-*EPHA5* interactions between Excit_L5/6 neurons and Excit_L6. The high colocalization score in the spatial network analysis in (**I**) suggests oligodendrocytes also likely co-exist with Excit_L6 neurons.

The Ephrin/Eph signaling system is critical for neuronal wiring during brain development and neural plasticity and synaptic homeostasis in adulthood (26). To better understand the role of this SCZ-associated signaling pathway in the adult DLPFC, we next characterized the cell types mediating this interaction using our snRNA-seq data. We identified enrichment of *EFNA5, EPHA5*, and *FYN* in excitatory neuron populations (**Fig. 5C**), which are the dominant sender and receiver cells for this LR interaction (**Fig. 5B**). Particularly, Excit_L5/6 neurons most specifically expressed the ligand *EFNA5 (Specificity Measure, SPM = 0*.*6847)* and its intracellular partner *FYN (SPM = 0*.*5345)*, while Excit_L6 neurons most specifically expressed the receptor *EPHA5* (*SPM = 0*.*6508*, **Fig S40B**). Furthermore, Excit_L5 and L6 neurons showed the highest co-expression of *FYN* and *EFNA5*, co-expressed in 87.25% of this population (**Fig. 5D**).

Since *EFNA5*-*EPHA5* is a contact-dependent interaction (26), we used the Visium data to spatially map sites of likely *EFNA5* and *EPHA5* crosstalk. Across data-driven SpDs, the highest proportion of spots co-expressing *EFNA5* and *EPHA5* localized to Sp_9_D_7_~L6 (p=1.7e-05, **Fig. 5E-F**). Consistent with snRNA-seq specificity analyses (**Fig S38B**,**C**), spots co-expressing *EFNA5* and *EPHA5* showed a higher predicted proportion of Excit_L5/6 neurons and Excit_L6 neurons compared to spots lacking co-expression (**Fig. 5H, Fig S41**). Spatial network analyses further supported that co-localization of *EFNA5* and *EPHA5* occurs frequently in spots containing Excit_L6 neurons - with strongest co-localization relationships between Excit_L6/Excit_L5 neurons, Excit_L6/Excit_L4 neurons and Excit_L6/oligodendrocytes (**Fig. 5I, Fig S38D**). Spatial mapping of *EFNA5* and *FYN* interactions also showed significant co-expression of these genes in Sp_9_D_7_~L6 (p=0.0046, **Fig S38F-G**) with frequent co-localization between Excit_L5_L6 and Excit_4 neurons (**Fig S40H-I**). In summary, we demonstrate the utility of this integrated single cell and spatial transcriptomic data for identifying and mapping disease-associated interactions in spatially localized cell types across the human DLPFC.

### Spatial registration of cell populations across neuropsychiatric disorders

To leverage the large amount of snRNA-seq data collected across the PsychENCODE consortium (PEC) (27), we spatially registered 6 DLPFC snRNA-seq datasets generated in the context of several brain disorders (Autism Spectrum Disorder [ASD], SCZ, bipolar disorder, and Williams Syndrome) to both the histological layers and unsupervised SpDs annotated in Visium data (**Fig. 6A, Fig S42**). Across the consortium, in neurotypical controls, we found that excitatory neuron subtypes with a laminar annotation spatially register to the relevant histological layers and converge on the same unsupervised SpDs. As expected, most inhibitory populations registered to multiple histological layers and unsupervised SpDs, with the exception of Pvalb and VLMC subtypes, which mapped to Sp_9_D_8_~L4 and Sp_9_D_1_~L1, respectively. Finally, glial populations also showed expected spatial registration with astrocytes strongly mapping to L1-associated SpDs, oligodendrocytes and OPCs strongly mapping to the WM, and endothelial, pericyte (PC), and smooth muscle cells (SMC) mapping to the newly characterized vascular domain Sp_9_D_1_.

**Figure 6.**
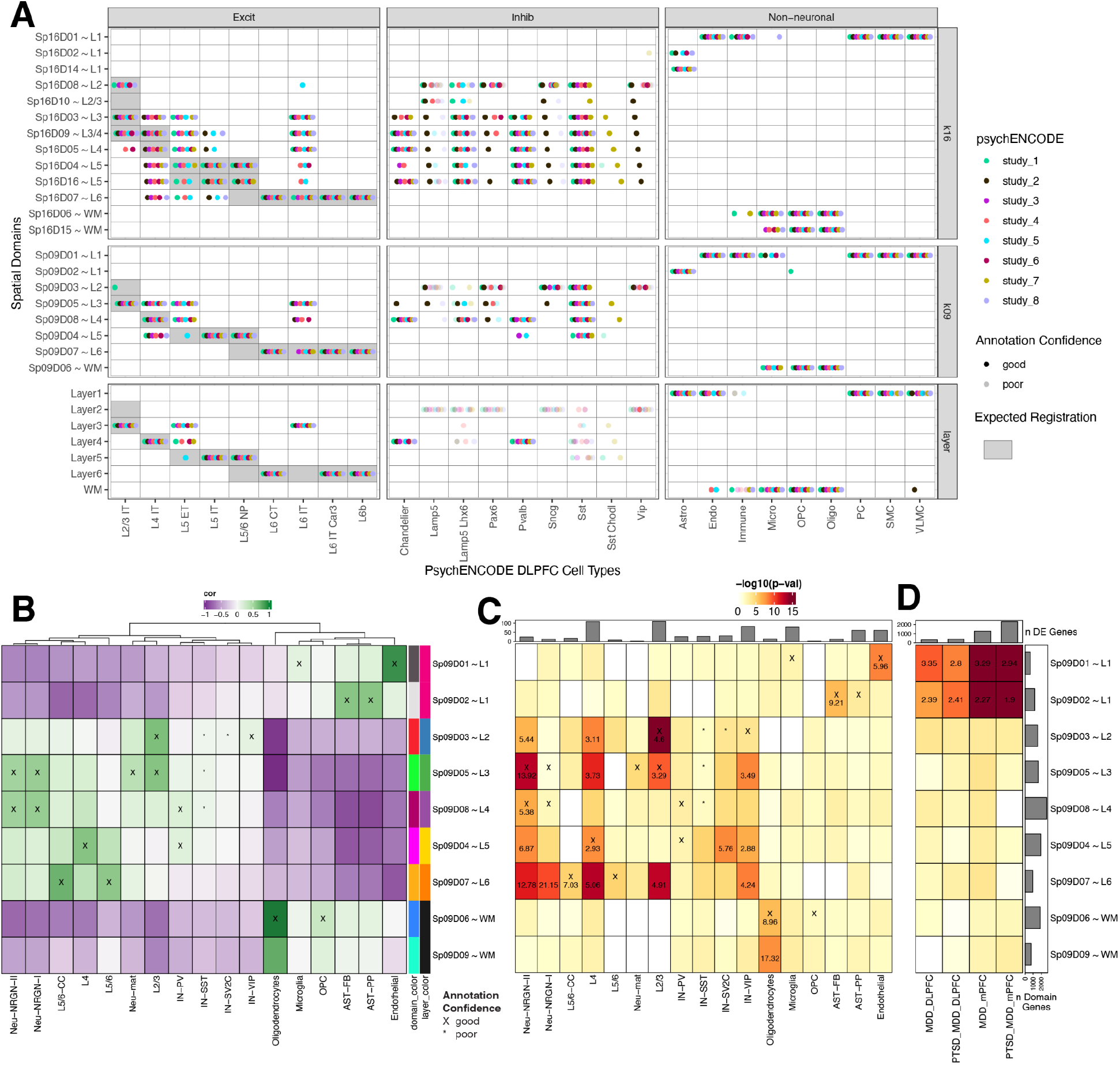
Spatial enrichment of cell types and genes associated with neurodevelopmental and neuropsychiatric disorders. (**A**) Dot plot summarizing spatial registration results for eight PsychENCODE (PEC) snRNA-seq datasets from human DLPFC. snRNA-seq data was uniformly processed through the same pipeline and annotated with common nomenclature based on work from Allen Brain Institute (27,47). Registration was performed for control donors only (see **Fig S42** for full dataset) across manually annotated histological layers from (6) as well as unsupervised *BayesSpace* clusters at *k*=9 and *k*=16 (Sp_9_Ds and Sp_16_Ds, respectively). Each dot shows the histological layer(s) or SpD(s) where a dataset’s cell type was annotated during spatial registration. Solid dots show good confidence in the spatial annotation, translucent dots show poor confidence in the annotation. IT, intratelencephalon-projecting; ET, extratelencephalon-projecting; CT, corticothalamic-projecting; NP, near-projecting; VLMC, vascular lepotomeningeal cell; OPC, oligodendrocyte precursor cell; PC, pericyte; SMC, smooth muscle cell. (**B**) Spatial registration of cell type populations from control samples from (14) against unsupervised *BayesSpace* clusters at *k*=9 (Sp_9_Ds). Higher confidence annotations (⍴ > 0.25, merge ratio = 0.1) are marked with an “X”. (**C**) Enrichment analysis using Fisher’s exact test for Sp_9_D-enriched statistics versus differentially expressed genes (DEGs, FDR < 0.05) in Autism spectrum disorder (ASD) for each cell type population. The values are the odds ratios (ORs) for the enrichment in significant cells, and the color scale indicates −log10(*p*-value) for the enrichment test. The top bar plot shows the number of DEGs for each cell type. (**C**) Enrichment analysis using Fisher’s exact test for Sp_9_D-enriched statistics versus differentially expressed genes (DEGs, FDR < 0.05) in Post Traumatic Stress Disorder (PTSD) and/or Major Depressive Disorder (MDD) in bulk RNA-seq of DLPFC and medial prefrontal cortex (mPFC). Top bar plot shows the number of DEGs for each DE test. Left bar plot shows the number of significantly enriched genes for each Sp_9_D in both enrichment analyses.

Next, we spatially registered snRNA-seq data from ~100,000 nuclei derived from the human prefrontal cortex and anterior cingulate cortex in a study of ASD that included 41 samples obtained from 31 donors (**Fig. 6B**) **(14)**. As expected, glial populations and laminar excitatory cell types mapped to the relevant SpDs (i.e. L2/3 neurons mapped to Sp_9_D_3_~L2 and Sp_9_D_5_~L3). We were also able to provide novel laminar assignments to some cell populations, such as mapping NRGN neuronal subtypes to Sp_9_D_5_~L3 and Sp_9_D_8_~L4. For each cell type, we next used clinical gene set enrichment analyses to assess the SpD enrichment of cell type-specific differentially expressed genes (DEGs) between individuals with ASD compared to neurotypical controls (**Fig. 6C**). Across many cell types, we observed multiple Sp_9_Ds enriched for ASD DEGs. Not surprisingly, Sp_9_D_3_~L2 showed significant enrichment of genes differentially expressed in L2/3 nuclei between individuals with ASD and neurotypical controls (p=2.60e-11), highlighting that these L2/3 DEGs are core Sp_9_D_3_~L2 marker genes. We also identified spatial enrichments for DEGs expressed in inhibitory neuron and neurogranin populations, including Sp_9_D_4_~L5 for SV2C inhibitory neurons and Sp_9_D_7_~L6 for Neu_NRGN_I neurons. Finally, to demonstrate how this large-scale dataset can be used to provide spatial information about genes associated with neuropsychiatric disease, we performed gene set enrichment analysis of bulk RNA-seq DEGs identified in a companion PEC study of PTSD and MDD (PEC study 6). For both DLPFC and ventral medial prefrontal cortex (mPFC), we demonstrate that vasculature domain Sp_9_D_1_~L1 and L1-associated domain Sp_9_D_2_~L1 are enriched in DEGs that are associated with both PTSD and MDD. This is consistent with previous studies implicating neuroimmune signaling in PTSD (28), and current PEC single cell analyses that implicate glial and vascular-cells in both MDD and PTSD (PEC study 6, personal communication). Together, these spatial registration and clinical gene set enrichment analyses add anatomical context to cell type identities and provide new biological insights into molecular changes associated with brain disorders, including ASD, MDD, and PTSD.

### Interactive single cell and spatial transcriptomic data resources

We provide several interactive web applications to explore this highly integrated DLPFC dataset as listed at http://research.libd.org/spatialDLPFC/#interactive-websites. First, we developed 2 *spatialLIBD* apps (29) that allow users to analyze Visium gene expression, cell segmentation, spot deconvolution, and clinical gene set enrichment results at *k*=9 and *k*=16. We added a third *spatialLIBD* app for the position differential expression results and a fourth for the Visium-SPG data. Second, both our Visium H&E (n=30 tissue sections) and Visium-SPG (n=4 tissue sections) data is available through a newly created performant web-based interactive visualization tool, called *Samui Browser (30)*, which allows rapid loading, visualization, and custom annotation of high resolution images and corresponding gene expression. Finally, snRNA-seq data at both fine and laminar resolution as well as pseudo-bulked Visium data (*k*=9, 16, and 28) are available through *iSEE* apps (31) that allows users to visualize expression levels for genes of interest through violin and heatmap plots. These large scale datasets and accompanying tools represent a landmark resource that can be used by both biologists to perform detailed molecular exploration of the DLPFC as well as method developers to benchmark novel computational tools for integrative analysis of single cell and spatial transcriptomic data.

## Discussion

Here we generated a large-scale, transcriptome-wide, data-driven molecular map across the anterior-posterior axis of adult human DLPFC from ten neurotypical control donors. This highly integrated single cell and spatial gene expression reference dataset enabled identification of novel unsupervised SpDs, which were characterized in terms of both cellular composition and domain-enriched genes, at different resolutions across the DLPFC. We provide a landmark molecular neuroanatomical atlas that complements our understanding of classic cortical cytoarchitecture through identification and characterization of discrete molecularly-defined layers and sublayers. In particular, we annotate a vasculature-rich meninges layer and several molecularly distinct subdomains in histological L1, 4, 5 and 6. An advantage of Visium over snRNA-seq approaches is its ability to capture transcripts in the cell cytoplasm and neuropil, which we speculate may influence identification of higher resolution spatial domains, particularly for demarcating laminar transitions and at the gray/white matter junction (32–34).

In this study, we provide a roadmap for the implementation and biological validation of unsupervised spatial clustering approaches in human brain tissue. While manual annotation of spatial domains is feasible for a limited number of samples in brain regions where neuroanatomical boundaries are well-characterized (6,16), the application of data-driven clustering methods is critical for future studies that aim to analyze spatial gene expression changes across diagnostic cohorts groups to identify changes in spatially-resolved cell types. Furthermore, unsupervised approaches will be essential for spatial profiling in brain regions that lack clear molecular or histological boundaries since they allow for identification of unknown or unexpected SpDs as well as SpDs that may be technically difficult to manually annotate (e.g. meninges). While here we evaluated several of the first-available spatial clustering algorithms, including *SpaGCN* and *BayesSpace* (7,9), there are a plethora of new tools coming online for both spatially variable gene detection, including *nnSVG (35)*, and also spatial domain identification, including *GraphST* and *PRECAST* (10,11). Large-scale, integrated datasets, including the present study, continue to offer developers of computational tools opportunities to develop novel methods scalable to atlas-level data, while also extracting meaningful biological information.

While Visium offers transcriptome-wide information at spatial resolution, a limitation of the platform is that spots often contain multiple cells and cell types. However, this can be theoretically overcome using spot-level deconvolution tools, as discussed here. We rigorously benchmarked the utility of these tools against Visium-SPG data in the DLPFC, where we manually assigned spots to the histological layers while also immunolabeling 4 broad cell populations in the same tissue sections. From these data we predicted the proportion of cell types in each spot, allowing us to achieve cellular resolution for our spatio-molecular map. While other imaging-based, spatially-resolved transcriptomics platforms, such as Xenium and MERFISH (36,37), directly measure transcripts in individual cells, only a limited number of genes can currently be probed. In contrast, the discovery-based approach afforded by Visium, as well as its scalability to a large number of samples, allowed for robust identification of novel spatial marker genes across many donors. These Visium-identified genes can be followed up at single cell resolution in smaller cohorts using probe-based approaches, such as MERFISH, which was recently applied to the middle and superior temporal gyri in the human brain (38).

Alterations in neural activity patterns within the DLPFC are noted in several neurodevelopmental and neuropsychiatric disorders (39–42), and it is hypothesized that changes in molecular signaling cascades may contribute to these alterations in activity. To gain insight into molecular dysfunction in DLPFC in the context of disease, we used our integrated molecular atlas to spatially map cell type-specific LR interactions that are associated with genetic risk for SCZ. For example, we highlight the interaction between *EFNA5* and *EPHA5* in excitatory L5/6 and L6 neuron subtypes in deep cortical layers, which is consistent with results from the most recently released SCZ genome-wide association study (GWAS) that identified enrichment of SCZ risk genes in glutamatergic neurons (43). Not only is *EFNA5* the locus of a GWAS-identified common SCZ risk variant, but it is also differentially expressed between individuals with SCZ and neurotypical controls in specific cell types (44). Spatially mapping disease-relevant LR pairs, which are often highly specific and druggable targets, can give new insights into pathophysiology and can help prioritize spatially restricted targets for therapeutic development. In combination with our interactive web resources, our highly integrated single cell and spatial transcriptomic data from neurotypical control DLPFC can be used to accelerate research across a variety of brain disorders by allowing researchers to search for relevant genes of interest, spatially register clinical gene sets, and explore disease-associated cell types for complementary assays, such as *in vitro* disease models.

Finally, spatial registration of eight DLPFC snRNA-seq datasets collected across the PsychENCODE consortium in the context of different neuropsychiatric disorders (27) revealed a convergence of excitatory, inhibitory, and non-neuronal cell types in relevant spatial domains. We observed increased confidence of inhibitory neuron mapping in our current expanded study compared to (6), likely due to the larger donor/sample number and data-driven clustering approach, which allowed for identification of finer resolution SpDs. Furthermore using ASD as an example (14), we demonstrate the utility of our data-driven molecular atlas for localizing cell-type specific DEGs to specific SpDs. For example, ASD DEGs in VIP inhibitory neurons were enriched in L3, L5, and L6-associated spatial domains, while those in SV2C inhibitory neurons were enriched only in the L5-associated domain. Together, these analyses provide anatomical context for cell type-specific gene expression changes and molecular mechanisms associated with neurodevelopmental disorders and psychiatric illness.

In summary, we provide a large-scale, highly integrated single cell and spatial transcriptomics resource for understanding the molecular neuroanatomy of the human DLPFC. We share web-based tools for the scientific community to interact with these datasets for further interrogation of molecular pathways associated with brain disorders.

## Supporting information

Table S1. Demographics of donors included in the study, data generation metrics, and SpaceRanger metrics for Visium and Visium Spatial SPG.

Table S2. snRNA-seq spatial registration correlation values and annotations.

Table S3. Summary of ligand-receptor pairs detected in data-driven cell-cell communication analysis.

Table S4. Summary of ligand-receptor pairs associated with SCZ genetic risk association.

Table S5. Summary statistics from t-tests on gene expression from artifact vs. non-artifact regions.

Table S6. P-values of differences in cell-type correlation across artifact vs. non-artifact regions.

Table S7. DEGs (FDR < 5%) for ANOVA model at broad and fine resolution for both Sp09 and Sp16 resolutions.

Table S8. DEGs (FDR < 5%) for the enrichment model at broad and fine resolution for both Sp09 and Sp16 resolutions.

Table S9. DEGs (FDR < 5%) for the pairwise model at broad and fine resolution for both Sp09 and Sp16 resolutions.

Table S10. DEGs (FDR < 5%) across Ant, Mid, Post positions for the anova, enrichment, and pairwise statistical models at the Sp09 resolution.

Table S11. snRNA-seq sample and quality control details.

Table S12. snRNA-seq cell type marker gene statistics.

Table S13. Correlation and RMSE for software spot-deconvolution results compared with CART predictions.

## Acknowledgements

We would like to thank Linda Orzolek and the Johns Hopkins Single Cell and Transcriptomics Core facility for executing all sequencing. We also acknowledge Elizabeth Engle and the Johns Hopkins Tumor Microenvironment Lab core facility for assistance with the Vectra Polaris slide scanner. We would like to thank the Joint High Performance Computing Exchange (JHPCE) for providing computing resources for these analyses. While an Investigator at LIBD, Andrew E. Jaffe helped secure funding for this work, and contributed to the conceptualization of the spatial registration framework. We thank William S. Ulrich for his assistance in deploying the Samui interactive websites. We thank Daniel R. Weinberger and members of the PsychENCODE consortium for feedback on the manuscript. We thank Amy Deep-Soboslay and her diagnostic team for curation of brain samples. We thank the neuropathology team, especially James Tooke, for assistance with tissue dissection. We thank the physicians and staff at the brain donation sites and the generosity of donor families for supporting our research efforts. Finally, we thank the families of Connie and Stephen Lieber and Milton and Tamar Maltz for their generous support of this work. Schematic illustrations were generated using Biorender.

## Authors Contributions

Conceptualization: LCT, KRM, KM, SCH

Methodology: LCT, LHM, KRM, KM, SCH

Software: AB, ASe, ASp,BG, CS, HD, LCT, LHM, MGP, MT, NE, PR, SCH

Validation: ASp, BG, KDM, KRM, LCT, LHM, MGP, MT, NE, SHK

Formal Analysis: ASp, AN, BG, HD, KRM, LCT, LHM, MGP, MT, NE, PR, SCH

Investigation: ASp, KDM, SHK, KRM, SCP, MNT

Resources: JEK, KRM, KM, LCT, SCH, TMH

Data Curation: LHM, LCT, NE

Writing-original draft: ASp, BG, KDM, KRM, KM, LCT, LHM, MGP, MT, NE, SCH, SCP, SHK

Writing-review and editing: KRM, KM, LCT, SCH, SCP

Visualization: ASp, LHM, KRM, LCT, PR, MGP, BG, SHK, NE

Supervision: AB, LCT, KRM, KM, MR, SCH, SCP

Project administration: LCT, KRM, KM, SCP

Funding Acquisition: LCT, KRM, KM, SCH

## Data and materials availability

The source data described in this manuscript are available via the PsychENCODE Knowledge Portal (https://PsychENCODE.synapse.org/). The PsychENCODE Knowledge Portal is a platform for accessing data, analyses, and tools generated through grants funded by the National Institute of Mental Health (NIMH) PsychENCODE Consortium. Data is available for general research use according to the following requirements for data access and data attribution: (https://PsychENCODE.synapse.org/DataAccess). For access to content described in this manuscript see: https://www.synapse.org/#!Synapse:syn51032055/datasets/. All figure and table files are available from https://www.synapse.org/#!Synapse:syn50908929. The source data are also publicly available from the Globus endpoint ‘jhpce#spatialDLPFC’ and ‘jhpce#DLPFC_snRNAseq’ that are also listed at http://research.libd.org/globus. The raw data provided through Globus include all the FASTQ files and raw image files. Processed data are publicly available from the Bioconductor package *spatialLIBD* version 1.11.7 (29) through the `fetch_data()` function. All source code is publicly available through GitHub and permanently archived through Zenodo at https://github.com/LieberInstitute/spatialDLPFC (45) and https://github.com/LieberInstitute/DLPFC_snRNAseq (46).

## Supplementary Materials List

Supplemental Methods.

Supplementary Figures S1 to S42.

Supplementary Table Legends.

Tables S1 to S13.

Supplemental References (48-75)

