## Supplementary Methods and Figures for "Integrated single cell and unsupervised spatial transcriptomic analysis defines molecular anatomy of the human dorsolateral prefrontal cortex"

**Supplemental Methods**

Post-mortem human tissue samples

Post-mortem human brain tissue was obtained at the time of autopsy with informed consent from the legal next of kin, through the Maryland Department of Health IRB protocol #12–24, and from the Western Michigan University Homer Stryker MD School of Medicine, Department of Pathology, the Department of Pathology, University of North Dakota School of Medicine and Health Sciences, and the County of Santa Clara Medical Examiner-Coroner Office in San Jose, CA, all under the WCG protocol #20111080. Demographics for the 10 donors are listed in **Table S1**. Details of tissue acquisition, handling, processing, dissection, clinical characterization, diagnoses, neuropathological examinations, and quality control measures have been described previously (48). Tissue blocks from the anterior (Ant), middle (Mid) and posterior (Post) positions along the rostral-caudal axis of human DLPFC were microdissected from the post-mortem brains of 10 adult neurotypical control donors (n=10 Ant, n=10 Mid, n=10 Post DLPFC blocks). Each tissue block was dissected from Brodmann Area 46 in a plane perpendicular to the pial surface to capture L1-6 and the WM.

Tissue processing and quality control

Following microdissection, tissue blocks were embedded in cold OCT and snap frozen in isopentane on dry ice and stored at -80℃. Tissue blocks were equilibrated to -20℃ before cryosectioning at a thickness of 10uM. Prior to cutting sections for Visium experiments, sections of the tissue blocks were cut to complete two quality control steps: 1) H&E staining to assess gross neuroanatomical structure, and 2) fluorescence multiplex RNAscope single molecule fluorescence *in situ* hybridization (smFISH) to ensure presence of all 6 layers and white matter (**Fig. 1**). H&E staining and RNAscope were performed according to manufacturer's instructions as previously described (49), and images were acquired using an Aperio CS2 slide scanner

(Leica) or a VectraPolaris slide scanner (Akoya Biosciences), respectively. For RNAscope, probes for established marker genes (ACD Bio) were used to identify laminae in the human cortex (6), including *SLC17A7* (Cat No. 415611), *RELN* (Cat No. 413051), *NR4A2* (Cat No. 582621), and *MBP* (Cat No. 411051).

Visium H&E data generation

One 10uM section from each of the 30 blocks was collected onto a 10x Visium Gene Expression slide, and processed according to manufacturer's instructions (10x Visium Gene Expression protocol) as previously described (6). Samples were processed in rounds by anatomical location (Ant, Mid, Post) with 4 different donors included on each slide. In brief, the workflow includes permeabilization of the tissue (18 minutes) to allow access to mRNA, followed by reverse transcription, collection of cDNA from the slide, and library construction. Libraries were quality controlled and sequenced on an Illumina NovaSeq 6000 at the Johns Hopkins Single Cell Transcriptomics core according to manufacturer's instructions at a minimum depth of 50,000 reads per Visium spot. Samples were sequenced to a median depth of 275,278,056 reads, corresponding to a median 67,211 of mean reads per spot, a median 2,257 of median unique molecular indices (UMIs) per spot, and median of 1,380 median genes per spot. After sequencing, we plotted established marker genes for neurons (*SNAP25*), white matter (*MBP*), and layer 5 (*PCP4*) to further validate the morphology of each tissue block (**Fig S1**, **Fig S2**, **Fig S3**).

Visium Spatial Proteogenomics (SPG) data generation

Immunofluorescence (IF) staining was conducted according to the manufacturer's instruction (catalog no.CG000312 Rev C, 10x Genomics). In brief, tissue blocks were cryosectioned at 10uM thickness and tissue sections were collected on a Visium Spatial Gene Expression Slide (catalog no. 2000233, 10x Genomics). Tissue was then fixed in pre-chilled methanol, treated with BSA-containing blocking buffer, and incubated for 30 minutes at room temperature with primary antibodies against the four cell-type marker proteins, NeuN for neurons, TMEM119 for microglia, GFAP for astrocytes, and OLIG2 for oligodendrocytes. For primary antibodies we used mouse anti-NeuN antibody conjugated to Alexa 488 (Sigma Aldrich, Cat# MAB377X, 1:100), rabbit anti-TMEM119 antibody (Sigma Aldrich, Cat# HPA051870, 1:20), rat anti-GFAP antibody (Thermofisher, Cat# 13-0300, 1:100), and goat anti-OLIG2 antibody (R&D systems, Cat# AF2418, 1:20). Following a total of 5 washes, secondary antibodies were applied for 30 minutes at room temperature. Detailed product information of the secondary antibodies is provided as follows: donkey anti-rabbit IgG conjugated to Alexa 555 (Thermofisher, Cat# A-31572, 1:300), donkey anti-rat IgG conjugated to Alexa 594 (Thermofisher, Cat# A-21209, 1:600), and donkey anti-goat IgG conjugated to Alexa 647 (Thermofisher, Cat# A-21447, 1:400). DAPI (Thermofisher, Cat# D1306, 1:3000, Final 1.67 µg/ml) was used for nuclear counterstaining. The slide was coverslipped with 85% glycerol supplemented with RiboLock RNase inhibitor (Thermofisher, Cat# EO0384, Final 2 U/µl) and scanned on a Vectra Polaris slide scanner (Akoya Biosciences) at 20x magnification with the following exposure time per given channel: 2.1 msec for DAPI; 143 msec for Opal 520; 330 msec for Opal 570; 200 msec for Opal 620; 1070 msec for Opal 690; 100 msec for autofluorescence (**Fig. 4A, Fig S26**).

Visium raw data processing and quality control (QC)

FASTQ and image data were pre-processed with the 10x *SpaceRanger* pipeline (version 1.3.0 for Visium H&E and 1.3.1 for Visium-SPG). Outputs of *SpaceRanger*, including the spotcounts matrix, were stored in a *SpatialExperiment* 1.6.0 (50) object (Bioconductor version 3.14) for downstream analysis. Data were filtered to remove genes that were not detected and spots with zero counts. We used *scran* 1.24.0 (Bioconductor version 3.14) (18) to calculate QC metrics, including mitochondrial expression, low library size, and low gene count. Only low library size spots were discarded: 4,866 (4.1%) spots, with 29 to 566 per sample with a median of 100 (**Fig S4**). We performed dimensionality reduction using the *scater* 1.22.0 package (51) to calculate the top 50 PCs followed by using the *Harmony* package 0.1.0 (52) to perform batch-correction for the effect of donors (**Fig S8**).

<u>Image segmentation and processing</u>

*Visium H&E*

The high-resolution images obtained on the 10x Genomics Visium platform were processed using *VistoSeg* (53), a *MATLAB*-based pipeline that integrates gene expression data with histological data. First, the software splits the high-resolution image of the entire slide into individual capture areas (.tif file format), which are used as input to 1) *Loupe Browser* (10x Genomics) for fiducial frame alignment, and 2) *SpaceRanger* 1.3.0 to extract the spot metrics/coordinates. We then used the `VNS()` function from *VistoSeg* to obtain the initial nuclei segmentations. For sections lacking tissue artifacts, we used `VNS()` nuclei segmentation outputs, and for tissue sections with artifacts like tears and folds, we applied a secondary refining step called the `refineVNS()` function, which segments more obvious nuclei from background signals. We then applied a watershed function to separate nuclei that were in close proximity and a size threshold to only retain segmented objects in the nuclei size range (30 to 4000 pixels). These final segmentations were used to obtain the nuclei count per Visium spot. We took two approaches for spot level counting with different stringencies: 1) we counted all segmented nuclei whose centroids were within a given Visium spot and 2) we calculated the proportion of pixels within a given Visium spot in which a segmented nucleus was present. A function in *VistoSeg* then uses the spot metrics and coordinates from *SpaceRanger* to integrate the segmented nuclei counts with gene expression data.

*Visium-SPG*

We leveraged the immunofluorescence (IF) images in the Visium-SPG assay to directly quantify cell type abundance and provide an imaging-based calculation of cell counts for comparison with outputs from spot deconvolution tools. Starting from the `cyto` model provided by *Cellpose* 2.0 (54), nuclear segmentations of Br2720_Ant_IF and Br6522_Ant_IF were iteratively improved in the *Cellpose* GUI, and a new model was trained until DAPI segmentations visually satisfied our accuracy standards. The final refined model was used to segment all four sections to produce masks. Segmentation masks were read into python 3.8.12, expanded to include a small region around each nucleus, and mean fluorescence intensity for the NeuN, OLIG2, and TMEM119-marked channels was quantified using `skimage.measure.regionprops_table()` from the *scikit-image (55)* v0.19.2 library. For each tissue section, an expert experimenter (co-author A.N.) used *Samui Browser (30)* to randomly select and annotate a spatially diverse set of 30 cells of each immunolabeled cell type – e.g. neuron, oligodendrocyte, astrocyte, microglia, or other (i.e. presence of DAPI, but absence of other immunofluorescent signals). The resulting dataset included 600 cells with a cell type label, and the corresponding mean fluorescence intensities from the respective channels.

To address concerns regarding possible biased selection of cells to annotate, a logistic-regression model was trained on the dataset using *scikit-learn* v1.1.1 (56), applied to predict un-annotated cells in all four sections. For each cell, the maximal probability of the model over any cell type was recorded as an indication of the model's prediction confidence. For each section and cell type, confidences were binned into four quartiles, and four cells from each such bin were randomly selected to form a list of 320 new cells to annotate. This process was designed to ensure a diverse set of cells, including some difficult-to-classify ones, were included in the full dataset, to improve model generalization. The original 600 cells were divided in an

80-20 training-test split, evenly stratified by section and cell type. The newer 320 cells were split 75-25, to ensure that 3 cells from each section, cell type, and quartile were included in the training set, with the remaining cell in the test set. A decision-tree classifier, built with `tree.DecisionTreeClassifier(max_depth = 4)` in *scikit-learn* (56), was trained, and an accuracy of 86.0% was achieved on the test data. The trained decision tree then predicted exact cell-type counts for each spot across the four Visium-IF sections, and this was compared against the results derived from *Tangram*, *Cell2location*, and *SPOTlight*.

Evaluating the impact of histology artifacts on Visium H&E data

The presence of artifacts, such as wrinkles and folds that result from tissue sectioning and sample preparation, can lead to misleading data. However, excluding spots that are believed to be artifacts diminishes sample size and the statistical power of downstream tests. To determine the impact of including spots from artifact regions in our analyses, an expert experimenter (K.R.M) manually annotated artifacts and histological layers for 3 samples: Br6522_Anterior, Br6522_Middle and Br6522_Posterior (**Fig S5A**). For each annotated sample, we used the one-sided Wilcoxon rank sum test to evaluate whether spots labeled as artifacts had greater library size, number of detected genes and estimated number of nuclei from *VistoSeg*, compared to spots that were not artifacts. Similarly, we evaluated whether spots in regions of artifacts were expected to have a lower percentage of mitochondrial gene expression. Since artifact spots were observed to have both a higher number of nuclei and higher library sizes (**Fig S5B**), we examined whether the changes in the density of nuclei had an effect on gene expression that was not accounted for by library size normalization. Specifically, we selected protein-coding genes and excluded genes expressed in fewer than 5% of all spots and fit the following regression model with a negative binomial (NB) error distribution and log-link function to estimate the effect of the presence of artifacts on the expression of each gene:

$$log(E[x_i]) = \beta_0 + \beta_a a$$

where $x_i$ is the vector of UMI counts assigned to gene $i$ and $a$ is a binary vector which indicates whether a spot is an artifact or not. Using the NB parameterization with mean $\mu$ and variance $\mu + \frac{\mu^2}{\theta}$, we estimate the intercept $\beta_0$, slope $\beta_a$ and overdispersion parameter $\theta$ for each gene, and regularize by aggregating information across genes with similar mean UMI counts using the *sctransform* v0.3.3 R package (57,58). Alternatively, we considered a negative binomial regression model with both library size and the presence of artifacts as independent effects:

$$log(E[x_i]) = \beta_0 + \beta_a a + \beta_l \, log_{10} l$$

where $l_j = \sum_i x_{ij}$ is the library size of spot *j*. When we include library size as an independent effect we observed that the regularized effect corresponding to artifacts was non-existent across all three samples (**Fig S5C**). Finally, we compared the estimate of the mean and variance of log-normalized gene expression computed across all spots to estimates obtained after excluding artifacts and found them to be similar. (**Fig S5D**).

To evaluate the impact of artifacts on spot-level downstream analyses, including clustering, we considered the complete dataset of 36,601 genes and all spots which were annotated to be found in a region with tissue. We first excluded genes with zero expression across all cells and spots which were low-quality due to inadequate library size as described previously. Since mitochondrial genes, ribosomal genes and *MALAT1* are highly expressed and believed to be technical rather than associated with a biological process of interest, we excluded them. We then log-normalized that expression data and selected the top 10% of highly variable genes to compute 50 expression PCs and obtained a UMAP embedding of the spots. We did not observe distinct clustering of the artifact spots when the data was visualized using UMAP (**Fig S6A**). Assuming that each PC (Principle Component) represents a distinct axis of variation unique to a subpopulation, we estimated the minimum number of PCs required to represent all subpopulations present, which corresponds to 10 PCs for Br6522_ant, Br6522_mid and 9PCs for Br8667_post and performed shared nearest neighbor clustering using the selected subset of PCs. For each cluster obtained, we characterized the fraction of spots which were artifacts and the relative layer proportions (**Fig S6B**). Further, we computed the variance explained by known sample characteristics including library size, percentage mitochondrial gene expression, layer of origin, and presence of artifacts using ANOVA (**Fig S6C**). Finally, we performed differential gene expression analysis for each sample and layer, comparing the expression in artifact spots to non-artifact spots using `PairwiseTTests()` and `combineMarkers()` from *scran* v1.21.1 and examined the log Fold-change in gene expression and corresponding *p*-values (**Fig S6D, Table S5**).

One possible manner in which artifacts can lead to misleading data occurs when a spot contains nuclei from two distinct biological regions of DLPFC (heterotypic spots) which would manifest either as a novel layer or lead to lack of separation between layers. In contrast, artifact spots formed due to wrinkling or folding of a single layer or transcriptionally similar layers (homotypic spots) are indistinguishable following normalization of gene expression. We artificially generated heterotypic spots across 28 potential layer combinations for Br6522_ant and Br6522_mid and 10 potential layer combinations for Br8667_post by randomly sampling weights $w_1$ and $w_2$ such that $0.3 \leq w_1, w_2 \leq 0.7$ and $w_1 + w_2 = 1$. We then computed the expression profile of a simulated heterotypic doublet by randomly selecting two spots from non-artifact regions corresponding to the two layers and computed the weighted sum where the weights are given by $w_1$ and $w_2$. We pooled simulated heterotypic doublets and observed samples, performed joint-data normalization and obtained a lower dimensional UMAP embedding (**Fig S7A**). For each spot we determined the 10 nearest neighbors using the shared nearest neighbor algorithm. Finally, we aggregated across all spots belonging to a specific artifact category and estimated the mean number of nearest neighbors that are non-artifacts,

artifacts and simulated doublets to evaluate whether artifact regions had a greater fraction of neighbors that were either alternate artifact spots or simulated doublets (**Fig S7B**). We observed that artifact spots did not disproportionately co-cluster with simulated heterotypic doublets. Alternatively, the presence of artifacts could lead to the observation of spots which have a cell-type composition that differs from non-artifact spots owing to overlapping biologically distinct layers. We used *Tangram* to assign cells of 7 distinct cell-types to both artifact and non-artifact spots in all three samples. For each spot, we summed across all cells belonging to a particular cell type to obtain the number of cells of each type present in the spot. We considered the Pearson correlation between a pair of cell-types within a layer to approximate how likely two cell types co-localized. Then we computed the cell-type correlation using all spots and compared it to the correlation obtained after excluding artifact spots to evaluate whether including artifact spots changed our beliefs about the cell type composition of spots using the `st1()` function from the *corTest* v1.0.7 R package. We then corrected the p-values obtained across all tests to obtain the FDR and observed no significant differences between to cell-type correlation computed from all spots and excluding artifact spots at FDR < 0.1 (**Fig S7C, Table S6**).

Unsupervised clustering of spatial data

To select an unsupervised clustering method for robust identification of spatial domains (SpDs) that best corresponded to cortical laminae in the Visium data, we evaluated three algorithms: shared nearest neighbors graph-based clustering (18), *BayesSpace* v1.5.1 (7), and *SpaGCN* v1.2.0 (9). These algorithms incorporated different features, including batch clustering (graph-based and *BayesSpace*), spatial awareness (*BayesSpace* and *SpaGCN*), and histology-weighted clustering (*SpaGCN*). Performance was benchmarked with our previously published DLPFC Visium data (12 sections from 3 donors; (6)) by comparing clustering accuracy against manual layer annotations using the Adjusted Rand Index (ARI), where a higher value represents greater similarity between predicted SpDs and manually annotated layers. Among the algorithms tested, *BayesSpace* most accurately identified SpDs consistent with histological cortical layers, and its performance improved with *Harmony* v0.1.0 batch correction (52) (**Fig S8**, **Fig S9**). We describe individual clustering methods below:

*Graph-based*

We performed graph-based clustering using `buildSNNGraph()` from *scran* and the Walktrap method implemented by *igraph* (17) v1.2.4.1 on 10 nearest neighbors and either 50 *Harmony* dimensions or 50 non-batch corrected principal components. Clustering was performed across all samples at once. We cut the graph at $k$ = 4 through $k$ = 28 in increments of 1 to explore different cluster resolutions.

*BayesSpace*

*BayesSpace* (7) is a spatially-aware clustering method. It is a Bayesian statistical model that relies on the spatial arrangement of Visium spots to borrow information from each spot's neighborhood for cluster assignments, thus resulting in smoother spatial clusters. To use *BayesSpace* on all samples at once, we arranged all samples into the same plane as recommended for joint clustering analysis https://edward130603.github.io/BayesSpace/articles/joint_clustering.html with a row offset of 100 per sample. We used `spatialCluster()` with `nrep = 10000` for `q` = 4 up to 28 applied on either the 50 non-batch corrected principal components or the 50 batch-corrected dimensions from *Harmony* (52). Note that *BayesSpace* is not guaranteed to result in $k$ equal to $q$ as it can merge smaller initial clusters (`init.method = "mclust"` by default) into a single spatial cluster.

*SpaGCN*

*SpacGCN* v1.2.0 (9) was applied following tutorial guidelines to assign spots to one of seven SpDs, for each sample individually. Default parameters were used for most functions, except where indicated otherwise in the tutorial. `Max_epochs = 200` was used in `SpaGCN.search_res()` instead of the default of `20`, to promote convergence and improve consistency between function executions. The model creation process, including invocations of `SpaGCN()`, `set_l()` and `train()`, was repeated until seven SpDs were found, as even with random seeds set, the process was not deterministic and was not guaranteed to find seven SpDs.

Identification of data-driven *k* for unsupervised clustering

To identify an optimal number of clusters for *BayesSpace* SpDs, we used a scalable discordance-based internal validity metric, *fasthplus* (59), to assess clustering performance for different values of *k* (**Fig S12**). Using *fasthplus* version 1.0, we calculated $H^+$ and then plotted 1-$H^+$ to identify the second inflection point of the plot as the point at which $H^+$ started to stabilize. Using this data-driven approach, we identified *k*=16 as the optimal number of clusters.

Layer-level data processing and differential expression modeling

At *k*=9, *k*=16, and *k*=28, `registration_pseudobulk()` from *spatialLIBD* v1.10.0 (29) was used to pseudo-bulk cells of the same sample ID and *BayesSpace*-determined spatial domain, and `filterByExprs()`, `cpm()`, and `calculateNormFactors()` from *edgeR* (60,61) were used to filter out lowly expressed genes and log-normalize counts.

We then identified differentially expressed genes using three models, as in our previous work (6) while adjusting for position in the anterior-posterior axis of the DLPFC, age, and sex. First, the ANOVA model used `registration_stats_anova()` from *spatialLIBD* and powered by *limma* (62) to test for differences in mean expression across $Sp_9Ds$ or $Sp_{16}Ds$ and identified robust DE (**Table S7**), with 11,341 (92.8%) DE genes (DEGs) across $Sp_9Ds$ (FDR < 0.05) and 9,042 (94.3%) across $Sp_{16}Ds$ (FDR < 0.05). Second, the enrichment model used `registration_stats_enrichment()` from *spatialLIBD* to test for differences in expression between one $Sp_kD$ versus all other $Sp_kDs$ and identified enriched genes for each $Sp_kD$: 5,931 and 4,471 unique enriched (t>0) genes for *k*=9 and 16, respectively. *CLDN5* was identified as being enriched in $Sp_9D_1$ **Fig. 2** (**Table S8**). Third, the pairwise model used `registration_stats_pairwise()` from *spatialLIBD* to test for DEGs between each pair of $Sp_kDs$ within a given resolution (36 pairs for $Sp_9Ds$ and 120 pairs for $Sp_{16}Ds$) resulting in 11,995 and 9,535 unique genes having at significant differences (FDR < 0.05) in at least one pair(**Table S9**).

Spatial registration of *Spatial Domains*

We validated and enhanced our understanding of the *BayesSpace* (7) SpDs by comparing them to the manual annotations of the pilot dataset (6) through a method of "spatial registration". We accessed the enrichment modeling results from Maynard et al. with *spatialLIBD* v1.11.4 `fetchData(type = "modeling_results")`. To compare the gene enrichments between the SpDs (Supplemental Methods: layer-level data processing and differential expression modeling) and the manual annotations, we calculated a Pearson's correlation between the enrichment t-statistics for each SpD and each manually annotated layer pair using only the union of the top 100 genes from each manual layer with *spatialLIBD* `layer_stat_cor(model = "enrichment", top_n=100)` (**Fig. 1E, Fig. 2B**).

Next to annotate SpDs with their best layer identity or hybrid-layer identity, we used `annotate_registered_clusters(cutoff_merge_ratio = 0.1)`. This function first finds the top correlation value for each SpD, if it's greater than the `confidence_threshold` of 0.25, the SpD's annotation has a good confidence (if not it is assigned poor confidence). Then the proportion of the difference between the top correlation and next highest correlation is compared to the `cutoff_merge_ratio`, if the proportion is lower the new layer is added to the list of associated layers. This process repeats until the ratio is higher than the cutoff as shown in this equation: `(cor`$_{current}$ `- cor`$_{next}$`) / cor`$_{next}$ `< cutoff_merge_ratio.`

This histological layer association is denoted for a given $Sp_kD_d$ by adding "~*L*" (e.g. $Sp_{09}D_{07}$~L6). Note that this SpD may not be identical to a histological layer as cytoarchitecture was not used to define layer boundaries. Our goal here was to highlight the most strongly associated histological layer to aid in biological interpretation.

Anatomical-level data processing and differential expression modeling

Using the same strategy and pseudo-bulked data as for the spatial domain differential expression analysis (Supplemental Methods: Layer-level data processing and differential expression modeling), we identified differentially expressed genes using three models while adjusting for $Sp_9$Ds, age, and sex. First, the ANOVA model used `registration_stats_anova()` from *spatialLIBD* to test for differences in mean expression across positions in the anterior-posterior axis of the DLPFC and identified 1,220 (9.98%) DEGs (**Table S10**). Second, the enrichment model used `registration_stats_enrichment()` from *spatialLIBD* to test for differences in expression between one position against the two other positions and identified 512 enriched (t>0) DEGs. Third, the pairwise model used `registration_stats_pairwise()` from *spatialLIBD* to test for DEGs between each pair of positions (3 pairs) resulting in 1,486 (12.2%) unique genes having significant differences (FDR < 0.05) in at least one pair.

snRNA-seq data collection and sequencing

Eighteen of the 30 tissue blocks were selected for single nucleus RNA-sequencing (snRNA-seq); the two "morphologically best" blocks were selected for each donor. "Morphologically best" was defined by: inclusion of all cortical layers, white matter, and few surface defects according to the layer definitions from the Visium data. Each selected block was sectioned using a Leica CM3050 at 100μm. Ten, 100μm sections were collected in lo-bind tubes for each respective sample, and these tissue sections were stored at -80°C until time of experiment. Nuclei preparations were conducted according to the 10x Genomics customer-developed “Frankenstein” nuclei isolation protocol as previously described in (63), with modifications designed to optimize the protocol for use with cryosections. Briefly, chilled EZ lysis buffer (MilliporeSigma #NUC101) was added to the lo-bind microcentrifuge tube containing cryosections, the tissue was broken up via pipette mixing, this lysate was transferred to a chilled glass dounce, the tube was rinsed with additional EZ lysis buffer, which was added to the respective dounce. The tissue was further homogenized using part A and B pestles for ~10 strokes each, and the homogenate was then strained through a 70μm cell strainer. After lysis, samples were centrifuged at 500g at 4°C for five minutes, supernatant was removed, and the sample was resuspended in EZ lysis buffer and re-centrifuged. After removing the supernatant, wash/resuspension buffer (PBS containing 0.5% BSA (Bovine Serum Albumin, Jackson ImmunoResearch #001-000-162) was added to the pellet. Upon resuspension, the samples were spun again and this wash process with wash/resuspension buffer was done three times. After the final wash and resuspension, propidium iodide (PI, 1:500, ThermoFisherScientific #P1304MP) was added, and the sample was strained through a 35μm cell filter attached to a FACS (fluorescence activated cell sorting) tube. For each sample, ~9000 PI+ nuclei were sorted in purity mode on a Bio-rad S3e Cell Sorter into 10X Genomics Reverse Transcription reagents without enzyme. Enzyme and water were added to bring the sample to the full volume, and the Chromium Next GEM Single Cell 3’ Reagent Kits v3.1 (Dual Index) kit was used to create libraries for sequencing according to the revision A protocol provided by the manufacturer (10X Genomics). Samples were processed in 5 rounds ranging from 1-4 samples per round.

snRNA-seq quality control, clustering, and annotation

Samples were sequenced to a median depth of 323,114,123 reads, corresponding to a median 55,811 mean reads per nucleus, median of 3,334 mean unique molecular indices (UMIs) per nucleus, and median 1,959 median genes per nucleus. We processed the sequencing data with 10x Genomics *Cell Ranger* v6.1.1, using human reference genome GRCh38. In total 26,605,806 (min=604,278, max=2,290,670 per sample) droplets were sequenced. Empty droplets were excluded using *DropletUtils* v1.14.2 `emptyDrops()` function (64) with a 30,000 iterations for the Monte Carlo *p*-value calculation, and a lower bound on UMI determined for each sample by the "knee point" calculated by *DropletUtils* `barcodeRanks()` function (65) plus 100, this value ranged from 219 to 910. Droplets with a significant deviation from ambient profile (FDR < 0.05) were kept, this resulted in a dataset of 84,756 nuclei (2,989 to 6,269 per sample).

For quality control (QC), nuclei were assessed for high percent mitochondrial reads, low total counts per nuclei, and low number of detected features. We applied an adaptive 3 median absolute deviation (MAD) threshold to each sample with *scater* v1.22.0 `isOutlier()` (51). If a nucleus failed any of the three QC tests, it was excluded. Next, doublet scores were computed by sample from the top 1000 highly variable genes with *scDblFinder* v1.12.0 (66) `computeDoubletDensity()`. This metric was later assessed at the cluster level (**Fig S19A**). Following quality control, a total of 77,604 (2,661 to 5,911 per sample) nuclei across 19 tissue blocks from 10 donors were included in the study (**Fig S18**, **Table S11**).

To perform dimension reduction, we first selected the top 2000 deviant genes from the raw count values with *scry* 1.6.0 `devianceFeatureSelection()` (67). Next, reduced dimensions were calculated with generalized linear models principal component analysis (GLM-PCA) (67), with *scry* `nullResiduals()` and *scater* 1.22.0 `runPCA()` (51). Initial reduced dimension plots showed evidence for batch effect (**Fig S20A**, **Fig S21A**). To correct for batch effects, we applied *Harmony* 0.1.0 `RunHarmony()` (52) to the GLM PC's with "Sample" as the group variable as it is a subset of both sequencing batch and individual. This returned corrected Harmony PCs, which upon visual inspection seemed to successfully correct for differences across different variables (**Fig S20B**, **Fig S21B, Fig S22**).

Next, we followed the hierarchical clustering procedure outlined in (63) in the batch corrected PC space, we performed graph-based clustering with scran v1.22.0 `buildSNNGraph()` with `k=20` and *igraph* 1.3.1 `cluster_walktrap()` to identify 296 preliminary nuclei clusters. After pseudo-bulking over the preliminary clusters, hierarchical clustering (hc) revealed 29 clusters. The clusters were annotated for broad cell type identity using established marker genes (63,68). When multiple cell types appeared to belong to the same broad cell type category, cell types were appended with a number (e.g. Excit_01) (**Fig S23**). One large cluster of cells did not have a clear cell type identity, and may have been driven by high mitochondrial rates and low UMIs (**Fig S19B)**. Examining the marker expression of this hc cluster at the preliminary cluster level showed that two out of six preliminary clusters had high expression for Oligo marker genes, and were therefore re-classified as Oligos. The other four preliminary clusters could not be assigned a clear cell type, so they were classified as a 30th "Ambiguous" cluster and excluded (**Fig. 3A**) resulting in a set of 56,447 nuclei. Doublets did not drive clustering in this dataset as no clusters had particularly high median doublet scores (**Fig S19A).**

<u>snRNA-seq spatial registration</u>

To add anatomical context to our cell populations, we performed spatial registration on the 29 hc cell type clusters with several functions from *spatialLIBD* v1.9.19. The enrichment statistics were calculated with `registration_wrapper()` with position, age, and sex used as covariates. These statistics were correlated against the manually annotated histological layers from

Maynard et al., as well as the $Sp_9D$ and $Sp_{16}D$ *BayesSpace* clusters with `layer_stat_cor()`, then annotated with `annotate_registered_clusters()` to assign the most strongly correlated histological layer or SpD. For $Sp_9D$ and $Sp_{16}D$ *BayesSpace* clusters, a stricter `cutoff_merge_ratio = 0.1` was used to identify more specific SpDs annotations (**Fig. 3B**, see details in Supplemental Methods: Spatial registration of *Spatial Domain*s). The histological layer assignments based on (6) were used to create a “layer level” cell type annotation for excitatory neuron clusters (e.g. Excit_L3). Thirteen, excitatory neuron clusters were grouped by their layer level assignments to seven layer resolution populations, three excitatory neurons clusters had poor annotation confidence and were classified as spatially ambiguous and excluded from the downstream analysis. Other non-excitatory cell types were grouped at broad resolution (**Fig. 3C, Fig S24**, **Table S2**). The spatially annotated set of 54,394 (921 to 5043 per sample) nuclei across 19 tissue blocks from 10 donors were used in downstream analysis.

snRNA-seq Azimuth validation

To support community efforts in using a common cell type nomenclature (69), we also took an alternative clustering approach and used the reference-based mapping tool *Azimuth* (https://azimuth.hubmapconsortium.org/ (20)), using *Azimuth* v0.4.5 `RunAzimuth(ref = "humancortex")`, to assign nuclei to subclasses (e.g. L5 extratelencephalon-projecting [E]T or L5 intratelencephalon-projecting [IT]) based on reference data derived from human motor cortex (21). We showed that a majority of our hierarchical cluster- and layer-resolution snRNA-seq clusters strongly correlate with those from *Azimuth*, with the exception of L4-associated excitatory neuron clusters (**Fig S25**), which is consistent with the absence of L4 in the agranular motor cortex reference dataset used by *Azimuth*.

Spot deconvolution benchmarking

Given that individual Visium spots often contain multiple cell types, we performed spot deconvolution to better understand the cellular composition of spots mapping to unsupervised spatial domains. While several algorithms have been developed to predict cell type proportions within individual Visium spots using single cell reference data (8,22,23,70,71), they have not yet been comprehensively benchmarked in brain tissues due to lack of a suitable reference dataset containing 1) paired Visium and snRNA-seq data from the same donors, 2) known cell type abundances in each Visium spot, 3) manually annotated Visium regions enriched in specific cell types. Therefore, we generated the first gold standard spot deconvolution dataset for 4 broad cell types in postmortem human DLPFC using the Visium Spatial Proteogenomics assay (hereafter referred to as Visium-SPG), which replaces H&E histology with immunofluorescence staining to label proteins of interest with fluorescent dyes (**Fig. 4A**). We selected 4 out of the 30 tissue blocks from independent donors and performed immunofluorescent staining for established cell type-specific proteins, including NEUN (marking neurons), OLIG2 (marking oligodendrocytes), GFAP (marking astrocytes), and TMEM119 (marking microglia). Following multispectral fluorescence imaging (**Fig S26**), we proceeded with the standard Visium protocol to generate corresponding gene expression libraries from the same tissue section (**Fig. 4B**, Supplemental Methods: Visium Spatial Proteogenomics (SPG) data generation).

Next, we benchmarked 3 recently published spot deconvolution algorithms: *SPOTlight*, *Tangram*, and *Cell2location* (8,22,23) using our Visium-SPG and snRNA-seq data. First, we identified a set of 25 robustly expressed marker genes for snRNA-seq clusters at both broad and layer-level resolution. For both the broad and layer-level cell-type resolutions, the `get_mean_ratio2()` function from the *DeconvoBuddies* v0.99.0 R package was applied to rank each gene in the snRNA-seq data in terms of its suitability as a marker for each possible target cell type. This method, which we call "mean ratio", determines the ratio of expression between a target cell type and the next-highest-expressing cell type. This is contrasted with a more conventional one-versus-all marker-finding strategy comparing the logarithm of the expression for a target cell type with the logarithm of the mean taken across all other cell types, an approach we denote “std.logFC” (**Fig S27**). Mitochondrial genes were excluded from the ranking process. The top-25-ranked markers by mean ratio were taken for each cell type, and it was verified the mean ratio exceeded 1 for each gene, to ensure greater expression in the target cell type than all others. This resulted in a total of 150 broad-resolution markers and 300 layer-level markers, used downstream for spot deconvolution (**Fig. 3D, Table S12**). We verified that Visium-SPG gene expression data reproduced expected laminar gene expression patterns (**Fig S28**). To confirm the utility of the snRNA-seq-derived marker genes for spot deconvolution, we evaluated their proportion of nonzero expression in Visium data (**Fig. 4B, Fig S29**) and confirmed that 25 genes per cell type were sufficient to identify laminar patterns (**Fig S30**). Using L5 excitatory neurons as an example, we found consistent spatial localization between L5 marker gene *PCP4* and the top 25 Excit_L5 marker genes identified in snRNA-seq data (**Fig. 4B**, **Fig S29)**.

Applying spot deconvolution softwares

Having verified the selection of robust marker genes for each cell type, we then ran *SPOTlight* (22), *Tangram* (8), and *Cell2location (23)* algorithms and calculated the predicted cell type counts per spot. *Tangram* v1.0.2, *Cell2location* v0.8a0, and *SPOTlight* v1.0.0 were applied with default parameters following the respective tutorials provided by their authors, with a few exceptions. The marker genes derived from the snRNA-seq data were used as training genes for all three tools, though for *SPOTlight*, ribosomal genes were excluded following tutorial recommendations, resulting in the exclusion of 6 and 2 markers from the layer-level and broad resolutions, respectively. At both resolutions, the cell-type-signature model was trained for 400 epochs rather than the default of 250 as this increase in training steps seemed to improve convergence of the model, which was recommended by the authors. Additionally, after noticing significant variation between *SPOTlight* results run with different random seeds subsetting to the default of 100 cells per cell type in the single-cell reference data, the final *SPOTlight* results were generated without any subsetting. This is consistent with the recommendations of the authors to use more than 100 cells for data with finer cell types, as was the case for our layer-level resolution data. *Tangram* was provided total cell counts per spot derived through segmentation with *Cellpose* v*2.0* (54,72) (Supplemental Methods: Image segmentation and processing), while *Cell2location* was provided an average count per spot, taken across all four sections. Importantly, these algorithms are not constrained to exactly match these input counts, and distort their values to differing extents, which is reflected in their outputs of cell type abundances (**Fig S31**). While *Tangram* and *Cell2location* predict cell-type abundances, *SPOTlight* predicts cell-type proportions, and is not subject to this distortion. Therefore, to more directly compare *SPOTlight* to *Tangram* and *Cell2location*, we multiplied the cell-type proportions outputted by *SPOTlight by* the *Cellpose*-derived counts to obtain abundances.

Evaluating performance of spot deconvolution methods

To quantify algorithm performance, we took 2 complementary approaches: 1) evaluating the localization of laminar cell types to their expected cortical layer and 2) comparing predicted cell type counts to actual counts obtained from immunofluorescent images. First, spots were manually assigned to L1-L6 or the WM using a combination of marker gene expression and cytotectonic architecture (**Fig. 4C**, **Fig S32**). Then for each cell type, we calculated the average predicted counts across manually annotated cortical layers. As expected, both *Tangram* and *Cell2location* show the highest predicted counts for L5 excitatory neurons in spots manually annotated to L5 (**Fig. 4C**). Across all cell types, *Tangram* and *Cell2location* showed similar layer-mapping performance; however, *Tangram* made more conservative predictions compared to *Cell2location* and performed more consistently across broad and layer level resolutions (**Fig S32**, **Fig S33**).

Next, using the immunofluorescent images, we calculated the actual number of neurons, astrocytes, microglia, and oligodendrocytes per spot by segmenting individual nuclei (72) and implementing a classification and regression tree (CART) approach (73) in *scikit-learn* (56) to categorize nuclei into the 4 immunolabeled cell types (**Fig S34**) (Supplemental Methods: Image segmentation and processing). We then compared the predicted cell type counts per spot from these softwares to the CART-calculated number of immunolabeled cells for both broad and layer level resolutions (**Fig. 4D**, **Fig S35**, **Fig S36**, **Table S13**). Counts for each software tool and CART-quantifiable cell type were summed across each Visium-SPG section. Totals for each software method (gene expression) were compared against CART-predicted (immunofluorescence) totals using Pearson correlation and root mean squared error (RMSE) (**Fig S37A**). Next, counts for each particular software tool and predicted by the CART were normalized to add to 1 across the four cell types. This allowed the calculation of the Kulback-Lieber divergence (KL divergence) from each software tool's predictions to the CART predictions. This treats the CART-predicted cell-type composition as a ground-truth probability distribution that each software tool is attempting to estimate. Given the challenge of deconvolving transcriptionally fine cell types, all softwares generally performed better (higher correlation, lower RMSE, and lower KL-divergence) at broad resolution compared to layer-level resolution. However, *Tangram* showed the most consistent performance at both broad and layer level resolution across all cell types and samples (**Fig. 4E**). This can at least partially be explained by the fact that *Tangram* spatially maps individual cells to perform deconvolution, while *Cell2location* and *SPOTlight* compute gene expression profiles for each cell type, which are then used for deconvolution. Since the same single-cell reference was used for deconvolution across all samples, *Tangram* was constrained to fix the proportions of cell types across sections, while *Cell2location* was able to more accurately capture dynamic changes in cell type proportions (**Fig S39**). In summary, our Visium-SPG benchmark analyses confirmed that both *Tangram* and *Cell2location* are suitable for spot deconvolution of Visium DLPFC data, but each tool has important caveats for data interpretation.

#### Visualization with *Samui Browser*

To facilitate interaction with H&E and IF images along with the transcriptome, we used a performant and interactive web-based data visualization tool for spatially-resolved transcriptomics data, *Samui Browser* (30). All Visium-SPG data and spot deconvolution results, including fluorescent images, nuclei segmentation, gene expression data, and cell type proportions, are available for further exploration at http://research.libd.org/spatialDLPFC/#interactive-websites.

#### Clinical gene set enrichment analysis

Differentially expressed genes for Autism Spectrum Disorder at the cell type level were reported by (14). For genes with a reported FDR < 0.05, we performed an enrichment analysis using BayesSpace $Sp_9Ds$ with *spatialLIBD* v1.11.4 `gene_set_enrichment()` (**Fig. 6C**). We performed the same enrichment analysis on differentially expressed genes for post traumatic stress disorder (PTSD) and major depressive disorder (MDD) on bulk RNA-seq reported through personal communication (PEC study 6, **Fig. 6**).

#### SCZ risk ligand-receptor occurrence

A SCZ risk gene list was obtained from the OpenTargets platform (https://www.opentargets.org/) under keyword “Schizophrenia.” (24) The top resulting list was filtered to only include genes with genetic association score > 0.1. To compare the occurrence of ligands and receptors in SCZ risk genes, we obtained a reference list of brain-expressed genes from GTEx data (https://storage.googleapis.com/gtex_analysis_v7/rna_seq_data/GTEx_Analysis_2016-01-15_v7_RNASeQCv1.1.8_gene_median_tpm.gct.gz). To create the brain-expressed gene list, tissues were filtered to those starting with 'Brain' and genes were filtered to have expression greater than zero. ENSEMBL genes were converted to symbol genes using the *mygene* library. Reference data for ligand or receptor classification assignment was obtained from the Omnipath database of interactions (25), importing only genes with categories 'ligand' or 'receptor'. A randomized set of genes equal in size to the number of SCZ risk genes obtained from Open Targets was sampled from the brain-expressed reference gene list. Ligand and receptor incidence was determined using the ligand/receptor reference data. This process was performed iteratively 10,000 times, with % incidence of ligands and receptors visualized as histograms. *p*-values were calculated by dividing the number of bootstrapped values greater than the inquiry value by the total number of statistics.

<u>SCZ risk-associated LR interactions of interest</u>

Among SCZ risk genes, ligand–receptor (LR) interactions where both interactors had an Open Targets genetic association score > 0.1 were prioritized (**Fig. 5A, Table S4**).

<u>Data-driven cell-cell communication analysis of snRNA-seq data</u>

Independently, we performed cell-cell communication analysis using all available normalized and annotated (at layer level resolution) snRNA-seq data via *LIANA* v0.1.8 (12). Ligand–receptor (LR) analysis was performed using `liana_wrap()`. We used `liana_aggregate()` to find consensus ranks of different methods. Interesting LR pairs were selected with the threshold `aggregated_rank<=0.01`. Complex heatmap and chord diagram (74,75) were used to display communication frequencies between cell type pairs (**Fig. 5A**) and cell communication pattern between *EFNA5* and *EPHA5* (**Fig. 5B**) respectively.

Cell specificity assessment of targets

Gene expression of *EFNA5*, *EPHA5* and *FYN* was visualized using the *scanpy* implementation for plotting dotplot (**Fig. 5C**). Cell specificity statistics in the preprocessed and annotated snRNA-seq data were calculated with the *tspex* python library. Two statistics were calculated: the tau statistic for determining whether a gene was specifically expressed, and the SPM statistic to identify the cell types gene expression was likely to be specific to. These results were visualized as *clustermaps* (**Fig S40B-C**).

snRNA-seq intracellular co-expression

For interactions predicted to occur intracellularly, we estimated the percent of nuclei co-expressing (raw counts > 1) both interactors of interest per cell type. These values were then visualized in the form of a clustermap (**Fig. 5D**).

Spatial co-expression

Preprocessed and annotated snRNA-seq and Visium data were used to verify co-expression of intracellular interactors of interest. Any nucleus (snRNA-seq) or spot (Visium) with raw counts > 0 was considered to be an expressor of a target of interest. When both the ligand and receptor had raw counts > 0, they were considered to be co-expressed in that nucleus or spot.

Cellular neighborhood network analysis

Spatial colocalization network analysis was performed using the deconvoluted spatial data for both *Tangram* and *Cell2location* results. In this analysis, the top cell types (**Fig. 5G**) with most likely contributions to a given spot were assumed to be co-localized in that spot. Co-localization relationships were scored in an adjacency matrix for spots where both the ligand and receptor were expressed. The adjacency matrix values were then divided by the absolute sum of the matrix. This was also performed for spots where neither the ligand nor the receptor were expressed. We took the ratio between both adjacency matrices and visualized this as a heatmap (**Fig. 5I**, **Fig S40 D-E**, **H-I**).

Spatial registration of PsychENCODE and other external snRNA-seq datasets

We leveraged the uniformly processed snRNA-seq data from eight PsychENCODE consortium (PEC) datasets (CapstoneII_Dataset_scRNAseq_HybridAnnotations_BICCN+Ma-etal-2022). All datasets were run through the same processing pipeline and annotated using consistent nomenclature (47). PEC studies are: 1) CommonMind Consortium (CMC), 2) DevBrain snRNA-seq, 3) IsoHub, 4) the snRNA-seq from this study generated at LIBD, 5) Multiome from the DLPFC, 6) PTSD from the Brainomics project, 7) schizophrenia and bipolar disorder with multi-seq (SZBDMulti-Seq), and 8) an ASD study from UCLA (UCLA-ASD). Spatial registration was completed with tools from *spatialLIBD* v1.11.4. The data was pseudo-bulked by cell type and sample, using `registration_pseudobulk()`, with `var_registration = "cellType"`. Clusters with less than 10 nuclei were excluded. The block correlation was computed by `registration_block_cor()`, on a model built with `registration_model()`. Enrichment statistics were calculated with `registration_stats_enrichment()`, utilizing the block correlation. This process is identical to using `registration_wrapper()` but was done piece wise to maximize efficiency as this analysis was completed with the full dataset (**Fig S42**) and a dataset filtered to include only neurotypical controls (**Fig. 6A**).

Enrichment statistics for the snRNA-seq dataset was completed with `registration_wrapper()`, after filtering to only neurotypical controls (14).

These statistics were correlated against the manually annotated histological layers from (6), as well as the $Sp_9D$ and $Sp_{16}D$ *BayesSpace* clusters with `layer_stat_cor()` and then annotated with `annotate_registered_clusters()`. For $Sp_9D$ and $Sp_{16}D$ *BayesSpace* clusters, a stricter `cutoff_mere_ratio = 0.1` was used to identify more specific spatial domain annotations (**Fig. 6A-B**).

Statistics

Statistical tests were performed with R versions 4.1.1 to 4.2.2 with Bioconductor versions 3.14, 3.15, and 3.16.0, as well as python versions 3.8.12 to 3.10.8. In addition to software versions listed in the Supplemental Methods, R session information was recorded with `sessioninfo::session_info()` for many specific analyses on log files or scripts themselves and is available on GitHub and Zenodo (Data and materials availability).

**Supplementary Table Legends**

**Table S1.** Demographics of donors included in the study, data generation metrics, and SpaceRanger metrics for Visium (n=30) and Visium Spatial Proteogenomics (Visium-SPG, n = 4) slides.
**Table S2.** snRNA-seq spatial registration correlation values and annotations.
**Table S3.** Summary of ligand-receptor pairs detected in data-driven cell-cell communication analysis.
**Table S4.** Summary of ligand-receptor pairs associated with SCZ genetic risk association.
**Table S5**. Summary statistics from t-tests on gene expression from artifact vs. non-artifact regions.
**Table S6**. P-values of differences in cell-type correlation across artifact vs. non-artifact regions.
**Table S7.** DEGs (FDR < 5%) for ANOVA model at broad and fine resolution for both Sp09 and Sp16 resolutions.
**Table S8.** DEGs (FDR < 5%) for the enrichment model at broad and fine resolution for both Sp09 and Sp16 resolutions.
**Table S9.** DEGs (FDR < 5%) for the pairwise model at broad and fine resolution for both Sp09 and Sp16 resolutions.
**Table S10.** DEGs (FDR < 5%) across Ant, Mid, Post positions for the anova, enrichment, and pairwise statistical models at the Sp09 resolution.
**Table S11.** snRNA-seq sample and quality control details.
**Table S12.** snRNA-seq cell type marker gene statistics.
**Table S13.** Correlation and RMSE for software spot-deconvolution results compared with CART predictions.

### Supplementary Figures

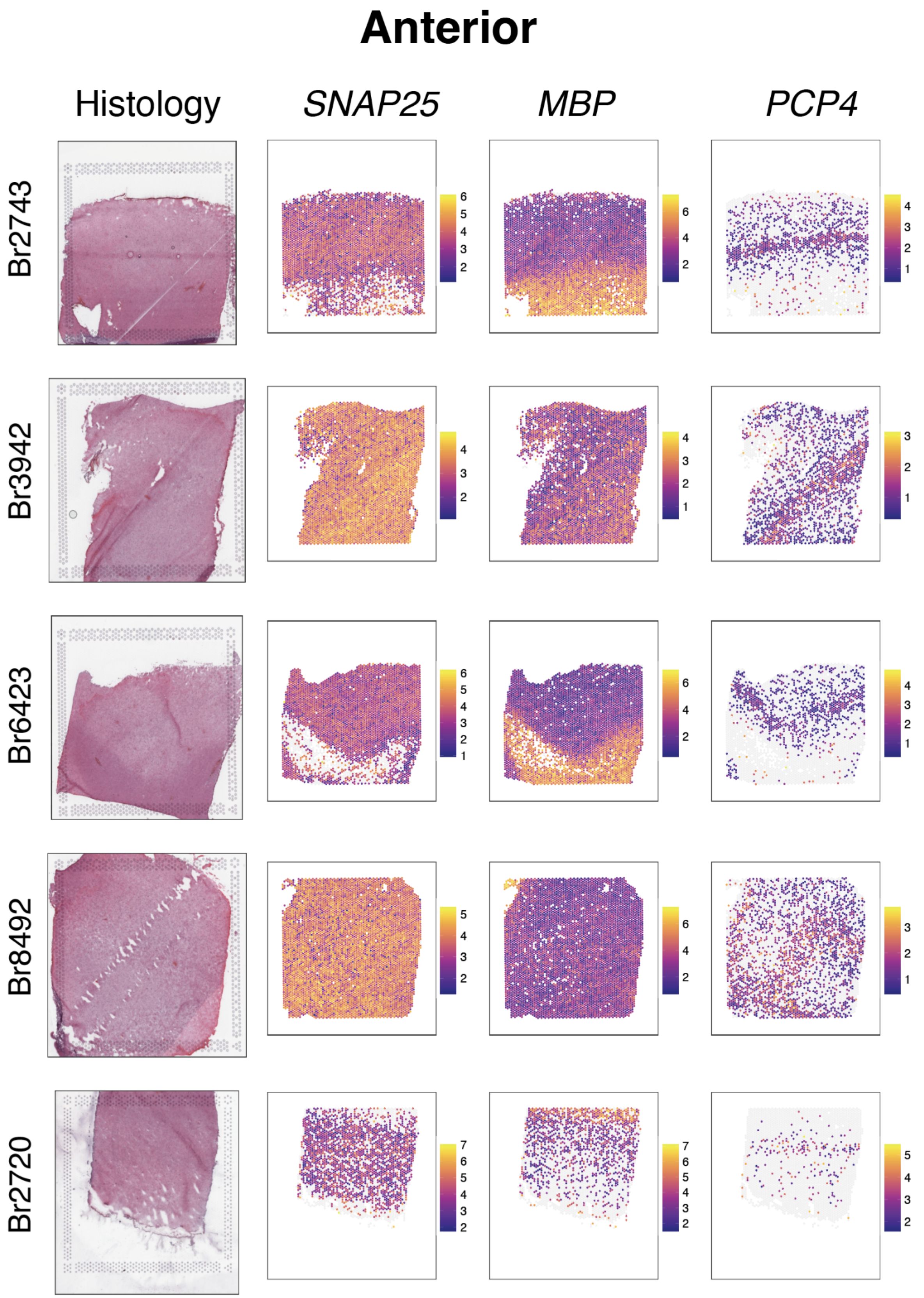
Anterior
Histology
SNAP25
MBP
PCP4
Br2743
Br3942
Br6423
Br8492
Br2720

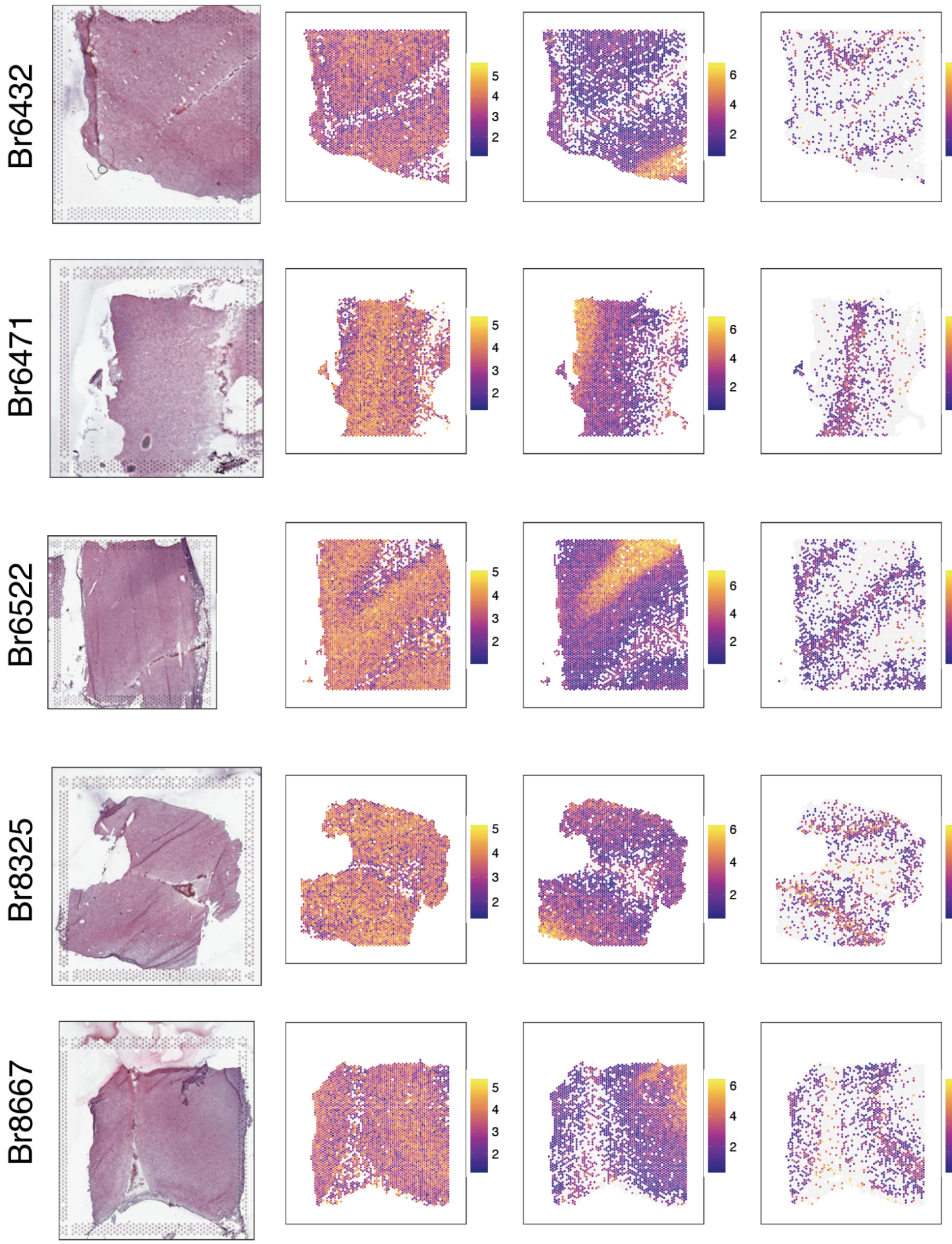


**Supplementary Figure 1. Expression of *SNAP25*, *MBP,* and *PCP4* confirm spatial orientation of anterior position DLPFC samples.** H&E histology, log transformed normalized expression (logcounts) for *SNAP25* (enriched in gray matter), *MBP* (enriched in white matter), and *PCP4* (enriched in layer 5) across 10 neurotypical control subjects (one subject per row).

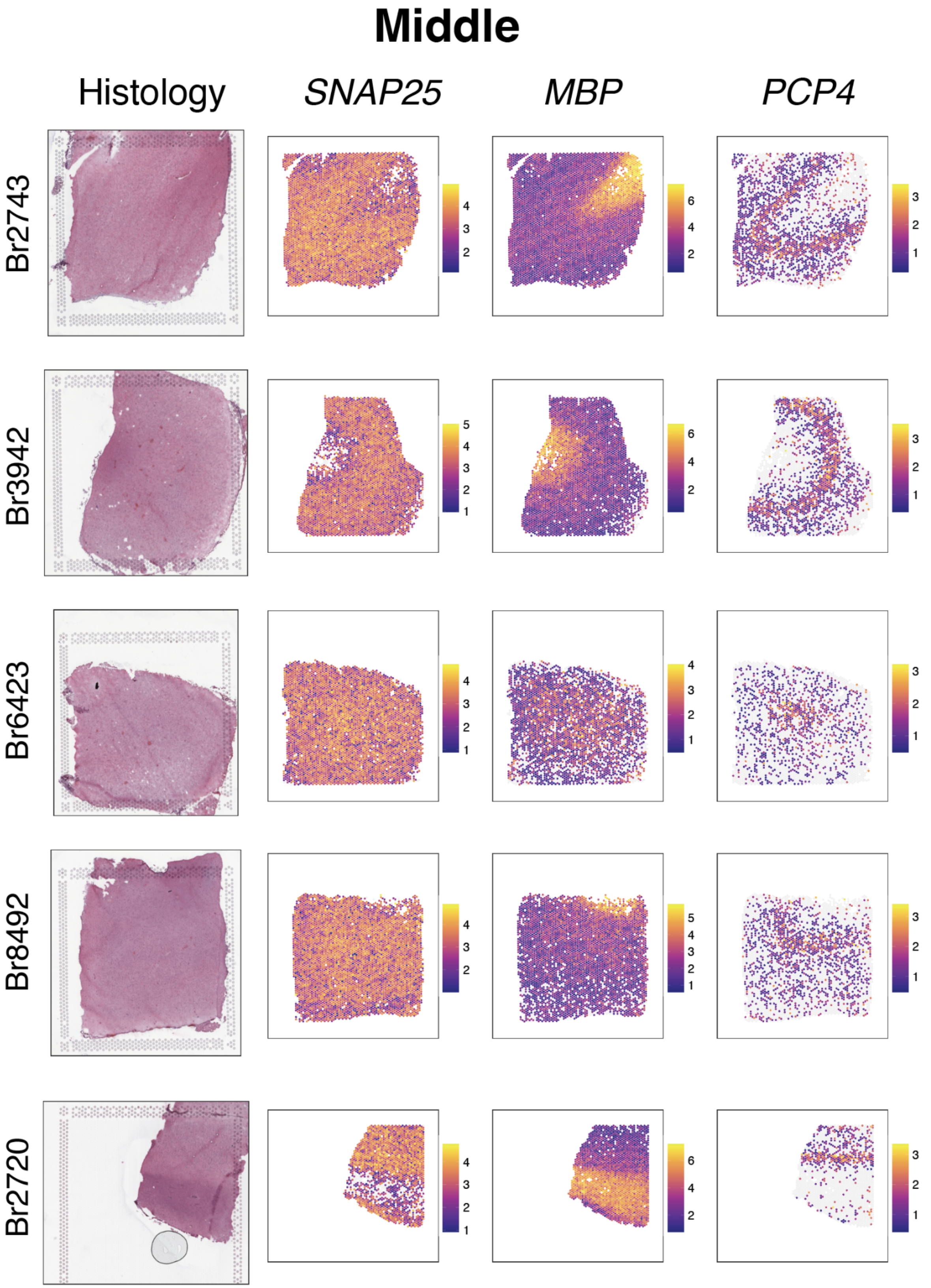

Middle
Histology
SNAP25
MBP
PCP4
Br2743
Br3942
Br6423
Br8492
Br2720

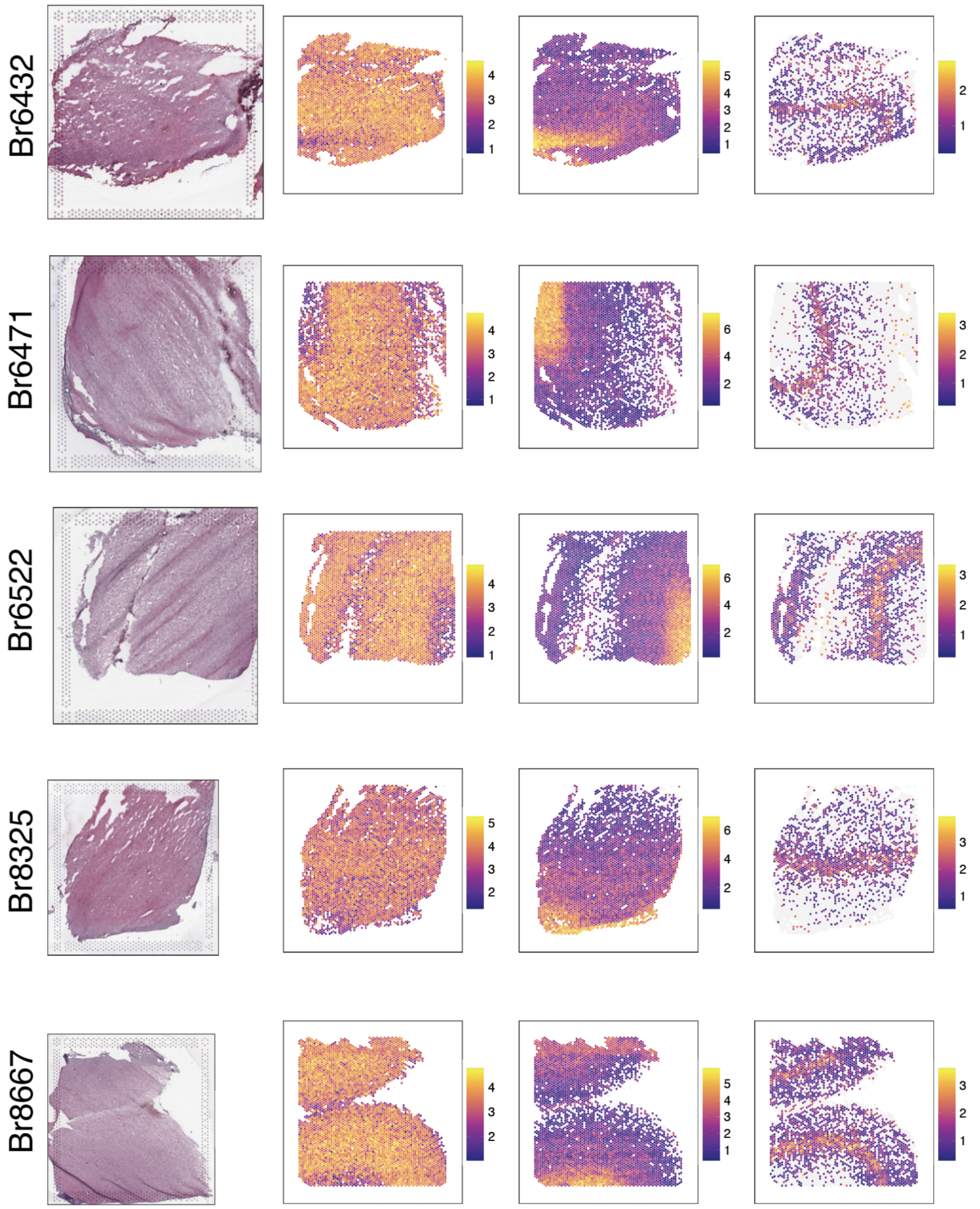


**Supplementary Figure 2. Expression of *SNAP25*, *MBP,* and *PCP4* confirm spatial orientation of middle position DLPFC samples.** H&E histology, log transformed normalized expression (logcounts) for *SNAP25* (enriched in gray matter), *MBP* (enriched in white matter), and *PCP4* (enriched in layer 5) across 10 neurotypical control subjects (one subject per row).

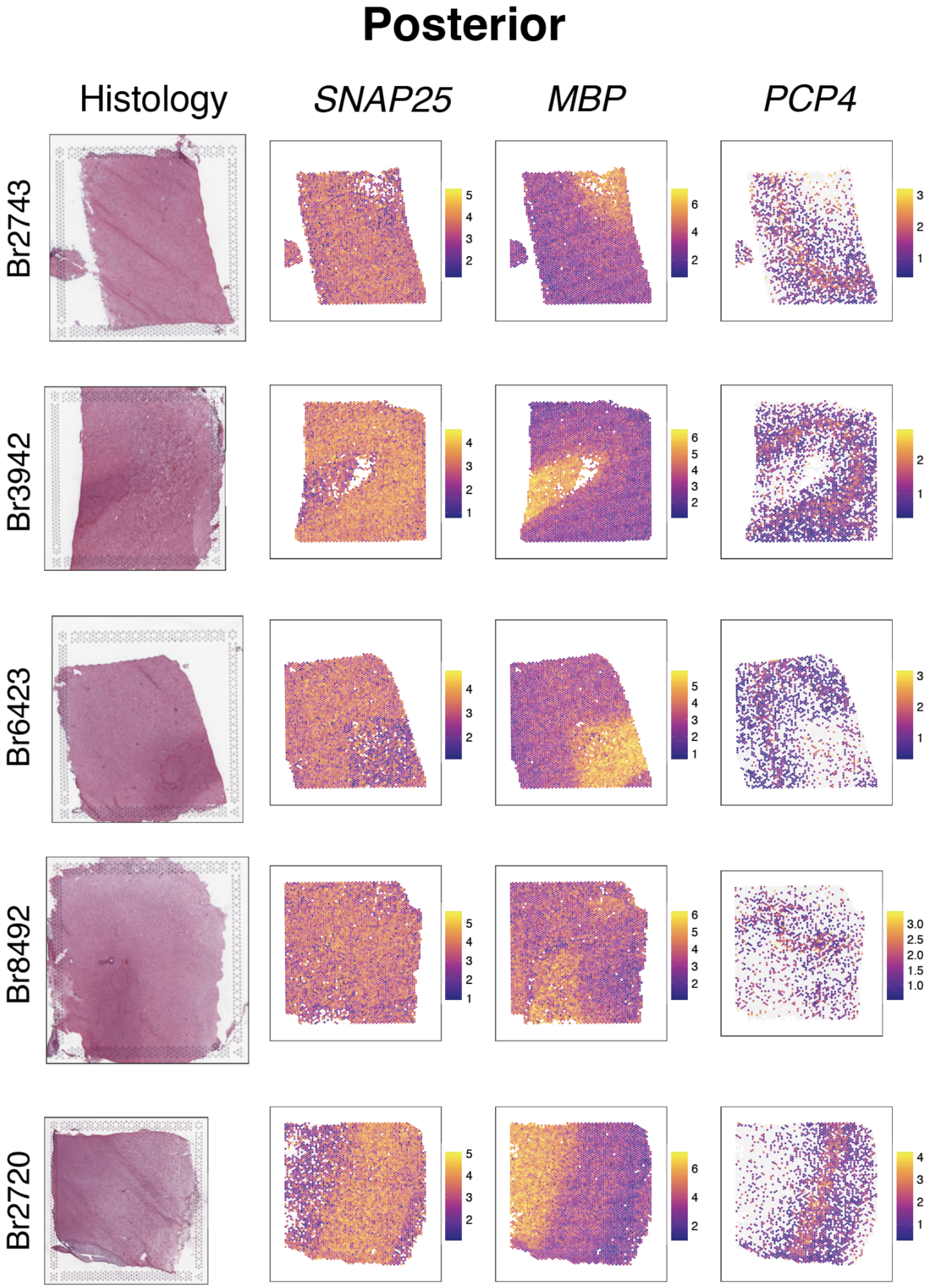
Posterior
Histology
SNAP25
MBP
PCP4
Br2743
Br3942
Br6423
Br8492
Br2720

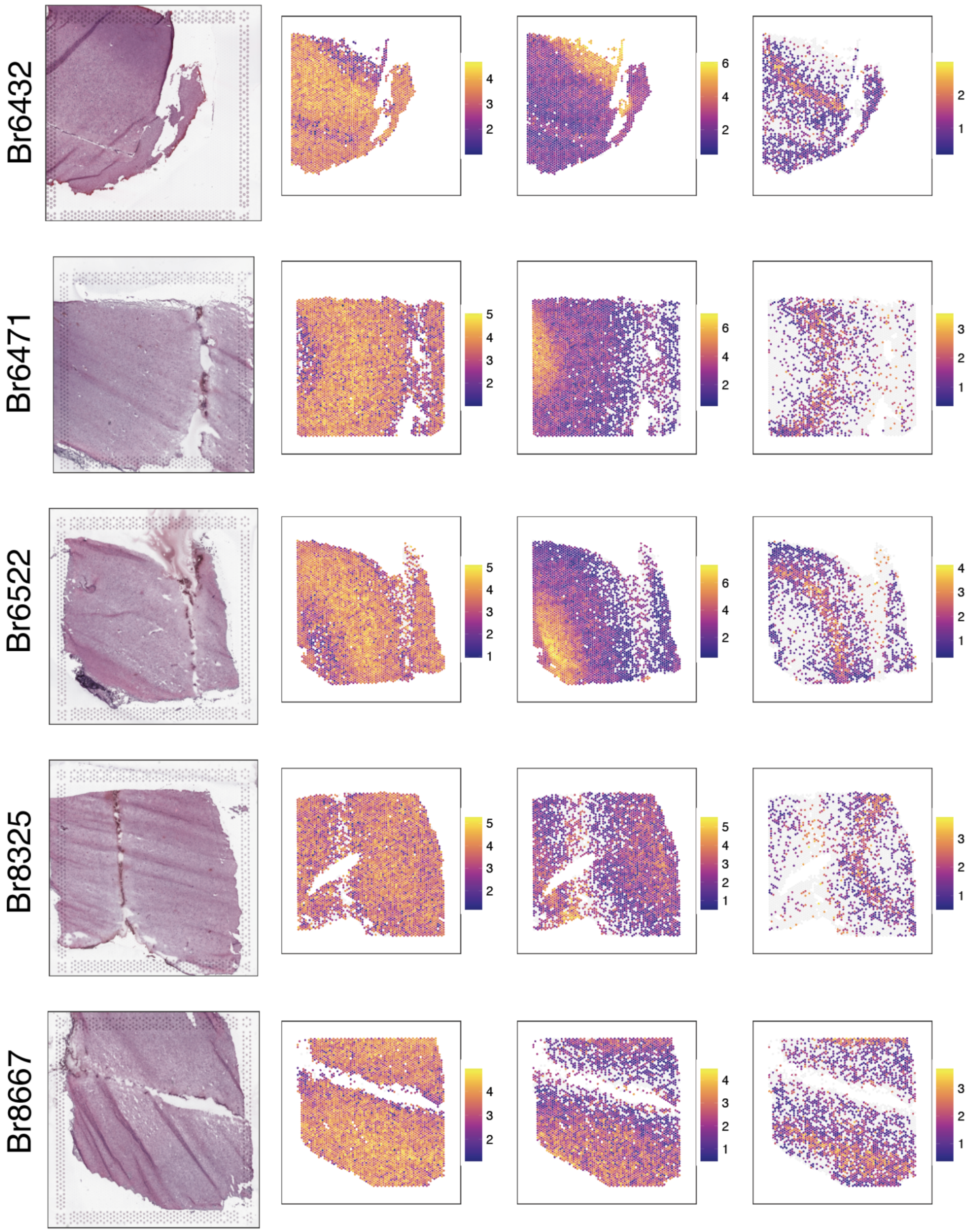


**Supplementary Figure 3. Expression of *SNAP25*, *MBP*, and *PCP4* confirm spatial orientation of posterior position DLPFC samples.** H&E histology, log transformed normalized expression (logcounts) for *SNAP25* (enriched in gray matter), *MBP* (enriched in white matter), and *PCP4* (enriched in layer 5) across 10 neurotypical control subjects (one subject per row).

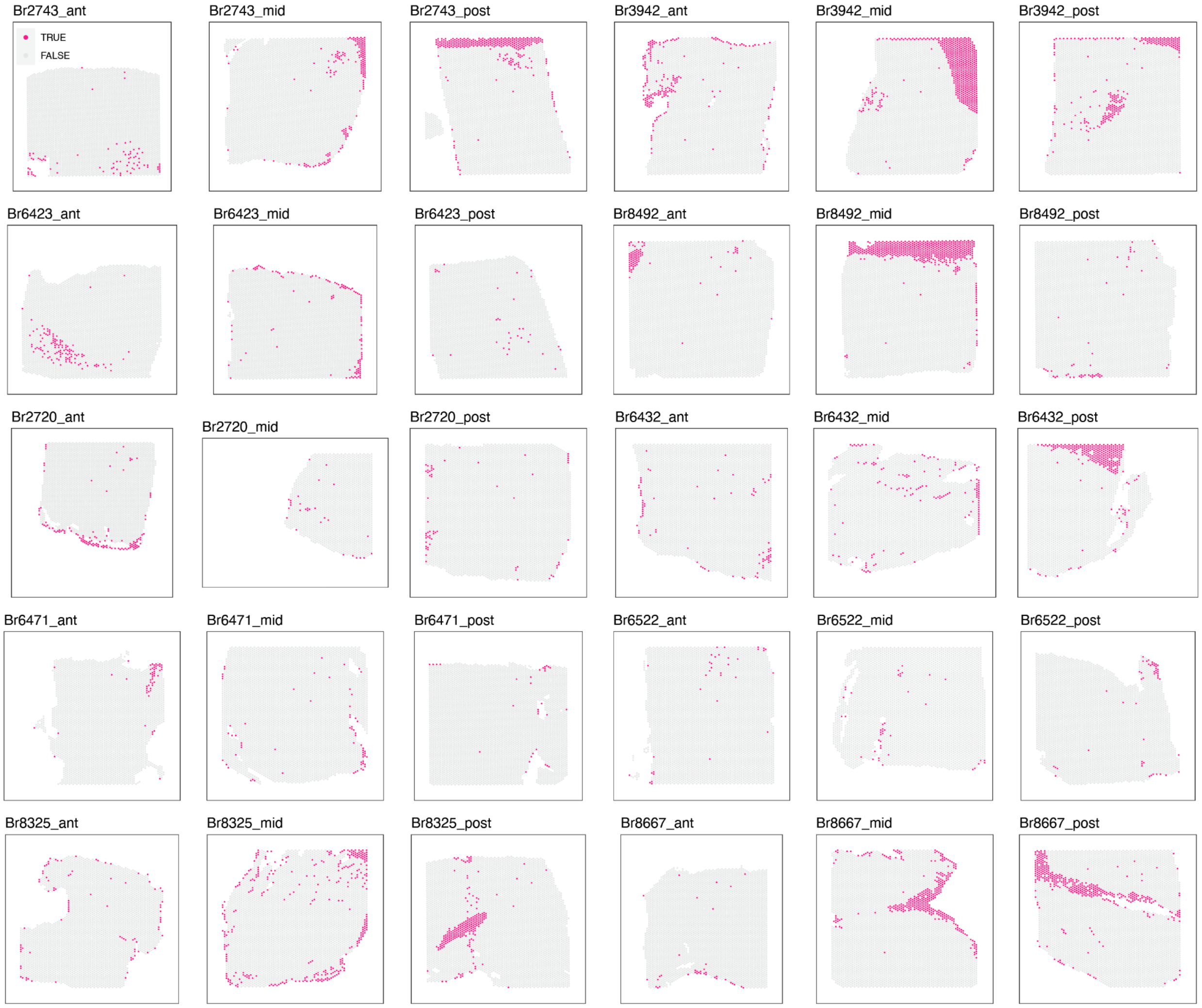


**Supplementary Figure 4**. **Quality control (QC) filtering of spots based on low library size**. Spots excluded from analysis for having a low library size as identified by *scran* for all 30 samples across the anterior-posterior axis of the DLPFC. Excluded spots are labeled in pink as TRUE.

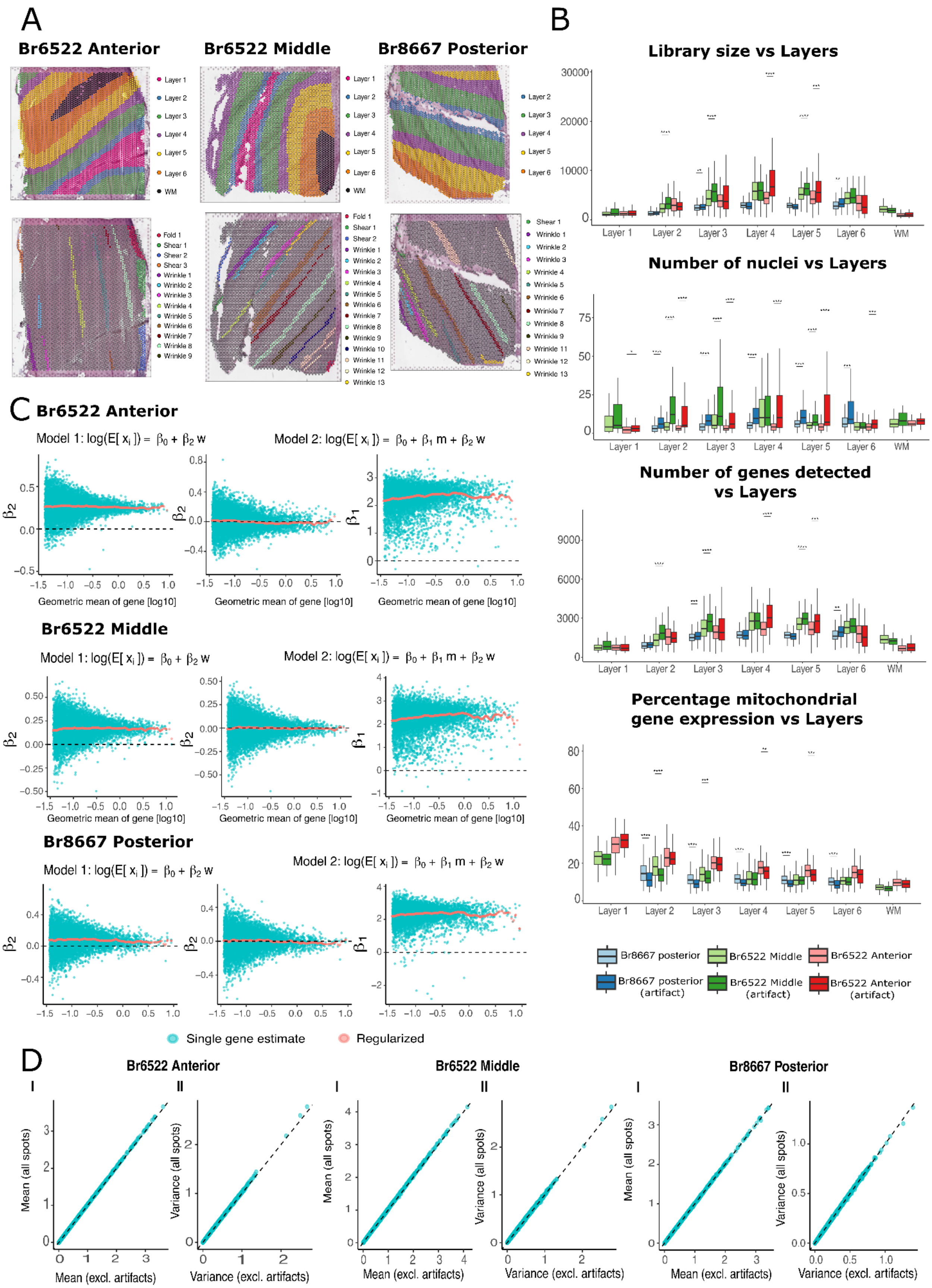

A
Br6522 Anterior
Br6522 Middle
Br8667 Posterior
Layer 1
Layer 2
Layer 3
Layer 4
Layer 5
Layer 6
WM
B
Library size vs Layers
Number of nuclei vs Layers
Number of genes detected vs Layers
Percentage mitochondrial gene expression vs Layers
Br8667 posterior
Br6522 Middle
Br6522 Anterior
Br8667 posterior (artifact)
Br6522 Middle (artifact)
Br6522 Anterior (artifact)
C
Br6522 Anterior
Model 1: log(E[ $x_i$ ]) = $\beta_0 + \beta_2$ w
Model 2: log(E[ $x_i$ ]) = $\beta_0 + \beta_1$ m + $\beta_2$ w
Geometric mean of gene [log10]
Br6522 Middle
Br8667 Posterior
Single gene estimate
Regularized
D
Br6522 Anterior
Br6522 Middle
Br8667 Posterior
I
II
Mean (all spots)
Mean (excl. artifacts)
Variance (all spots)
Variance (excl. artifacts)

**Supplementary Figure 5. Quality control (QC) assessment of impact of artifacts on gene-level downstream analyses.** (**A**) Manual annotations of spots from three tissue sections Br6522 anterior, Br6522 middle and Br8667 posterior corresponding to the presence of artifacts and the layer of origin (**B**) Comparison of common QC metrics between artifact spots and non-artifact spots for each layer in a sample. Library size was significantly greater in artifact spots for Layer 2 Br6522 middle (p=1.16e-05), Layer 3 Br8667 posterior (p=8.00e-03) and Br6522 middle (p=4.93e-11), Layer 4 Br6522 anterior (p=7.21e-08), Layer 5 Br6522 middle (p=8.54e-08), Br6522 anterior (p=6.42e-04), Layer 6 Br8667 posterior (p=4.00e-03). The number of nuclei detected in artifact spots was greater for Layer 1 Br6522 anterior (p=1.92e-02), Layer 2 Br8667 posterior (p=2.8e-06), Br6522 middle (p=1.67e-06), Br6533 anterior (p=9.33e-10), Layer 3 Br8667 posterior (p=2.34e-45) Br6522 middle (p=2.80e-20) Br6522 anterior (p=9.83e-18), Layer 4 Br8667 posterior (p=7.70e-18) and Br6522 anterior (p=1.04e-7), Layer 5 Br8667 posterior (p=2.64e-20) Br6522 middle (p=1.76e-09) Br6522 anterior (p=1.46e-15), Layer 6 Br8667 posterior (p=2.96e-04), Br6522 anterior (p=9.77e-04). The number of detected genes was significantly greater in artifact spots for Layer 2 Br6522 middle (p=3.04e-06), Layer 3 Br8667 posterior (p=1.70e-04), Br6522 middle (p=1.11e-10), Layer 4 Br6522 anterior (p=6.64e-8), Layer 5 Br6522 middle (p=9.17e-08), Br6522 anterior (p=1.72e-04), Layer 6 Br8667 posterior (p=1.00e-03). The percentage expression of mitochondrial genes was significantly lower in artifact spots for Layer 2 Br8667 posterior (p=3.60e-06), Br6522 middle (p=7.20e-07), Layer 3 Br8667 posterior (p=5.86e-22), Br6522 middle (p=1.20e-04), Layer 4 Br8667 posterior (p=2.86e-11) and Br6522 anterior (p=4.00e-03), Layer 5 Br8667 posterior (p=9.64e-15), Br6522 anterior (p=1.47e-04), Layer 6 Br8667 posterior (p=2.91e-06). (**C**) Comparison of the effect of artifacts on gene expression without including and including library size as an independent effect using negative binomial regression with a log-link function. Artifacts have a non-zero regularized effect size on gene expression across a range of mean gene expression values which reduces to zero when library size is included as an independent effect. (**D**) Comparison of mean gene expression and variance of gene expression inferred from log-transformed and normalized expression data computed using all spots (y-axis) and excluding artifact spots (x-axis). Excluding mitochondrial genes, ribosomal genes and *MALAT1* estimates of the mean and variance of gene expression are similar when artifact spots are included.

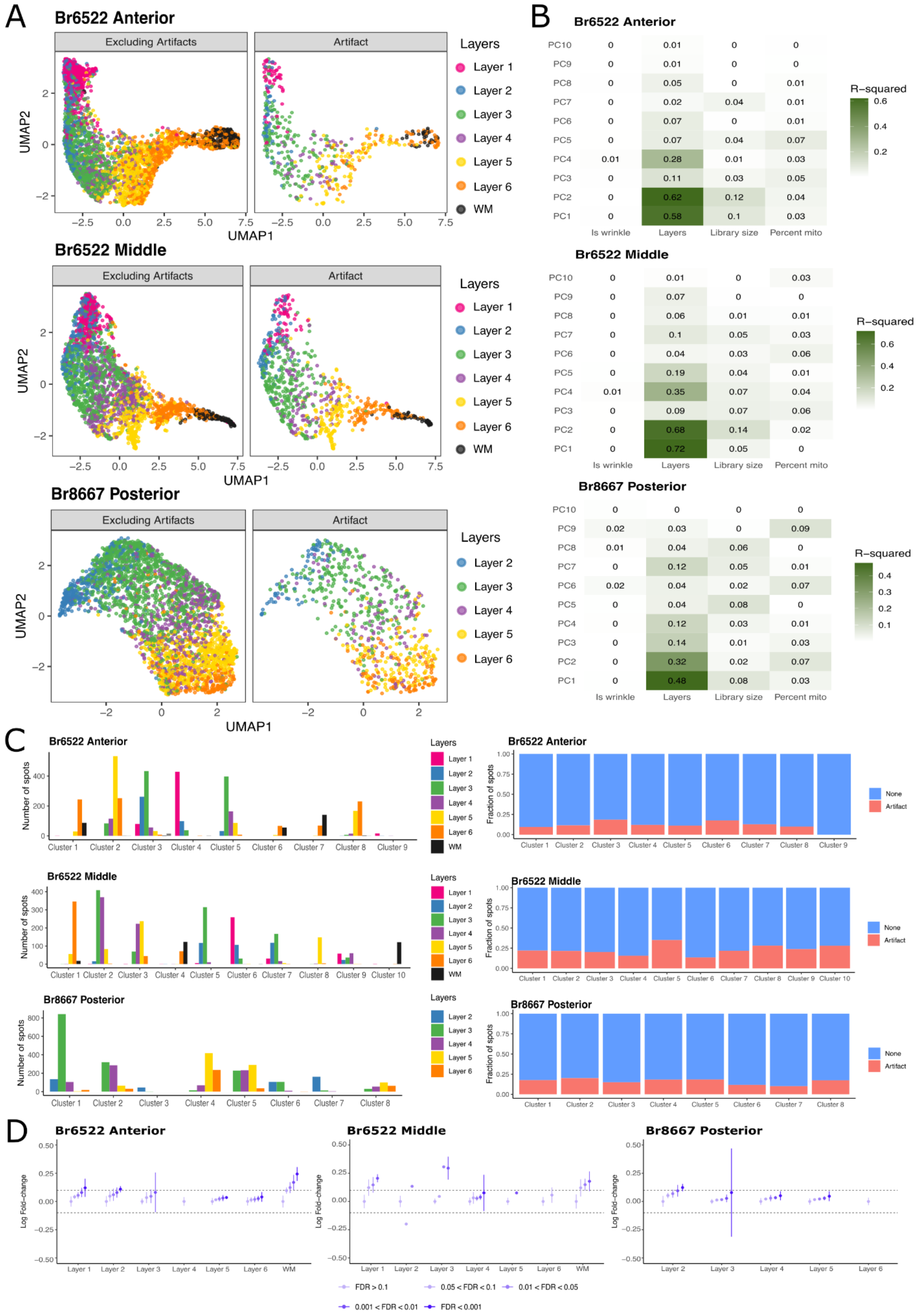

A
Br6522 Anterior
Excluding Artifacts
Artifact
UMAP1
UMAP2
Layers
Layer 1
Layer 2
Layer 3
Layer 4
Layer 5
Layer 6
WM
Br6522 Middle
Br8667 Posterior
B
Br6522 Anterior
R−squared
Is wrinkle
Layers
Library size
Percent mito
PC1
PC2
PC3
PC4
PC5
PC6
PC7
PC8
PC9
PC10
Br6522 Middle
Br8667 Posterior
C
Br6522 Anterior
Number of spots
Fraction of spots
None
Artifact
Br6522 Middle
Br8667 Posterior
D
Br6522 Anterior
Br6522 Middle
Br8667 Posterior
Log Fold−change
FDR > 0.1
0.05 < FDR < 0.1
0.01 < FDR < 0.05
0.001 < FDR < 0.01
FDR < 0.001

**Supplementary Figure 6. Quality control assessment of impact of artifacts spot-level downstream analyses.** (**A**) UMAP embedding of non-artifact and artifact spots from Br6522_ant, Br6522_mid and Br8667_post. (**B**) Percentage of variance of top 10 PCs explained by known sample characteristics. Variance of the top 4 PCs in all 3 samples Br6522_ant, Br6522_mid and Br8667_post is driven by layer of origin. 1% of the variance of PC4 is accounted to the presence of artifacts in Br6522_ant and Br6522_mid. 1-2% of the variance of PC5, PC8 and PC9 is explained by the presence of artifacts. (**C**) Frequency of spots belonging to each layer that are present in clusters obtained from shared nearest neighbors and relative proportion of non-artifact and artifact spots. Spots from adjacent layers such as layers 6 and WM, layer 5 and layer 6 and layer 4 and layer 5 tend to co-cluster implying transcriptionally similarity between spots from adjacent layers. The fraction of spots belonging to artifacts is similar across all clusters inferred from the 3 samples. (**D**) Distribution of log fold-change of gene expression (Median +/- 3MAD) in non-artifact spots compared to artifact spots for each layer and sample binned by FDR. Across all samples the absolute log fold change was not greater than 1 at FDR < 0.01 in any layer.

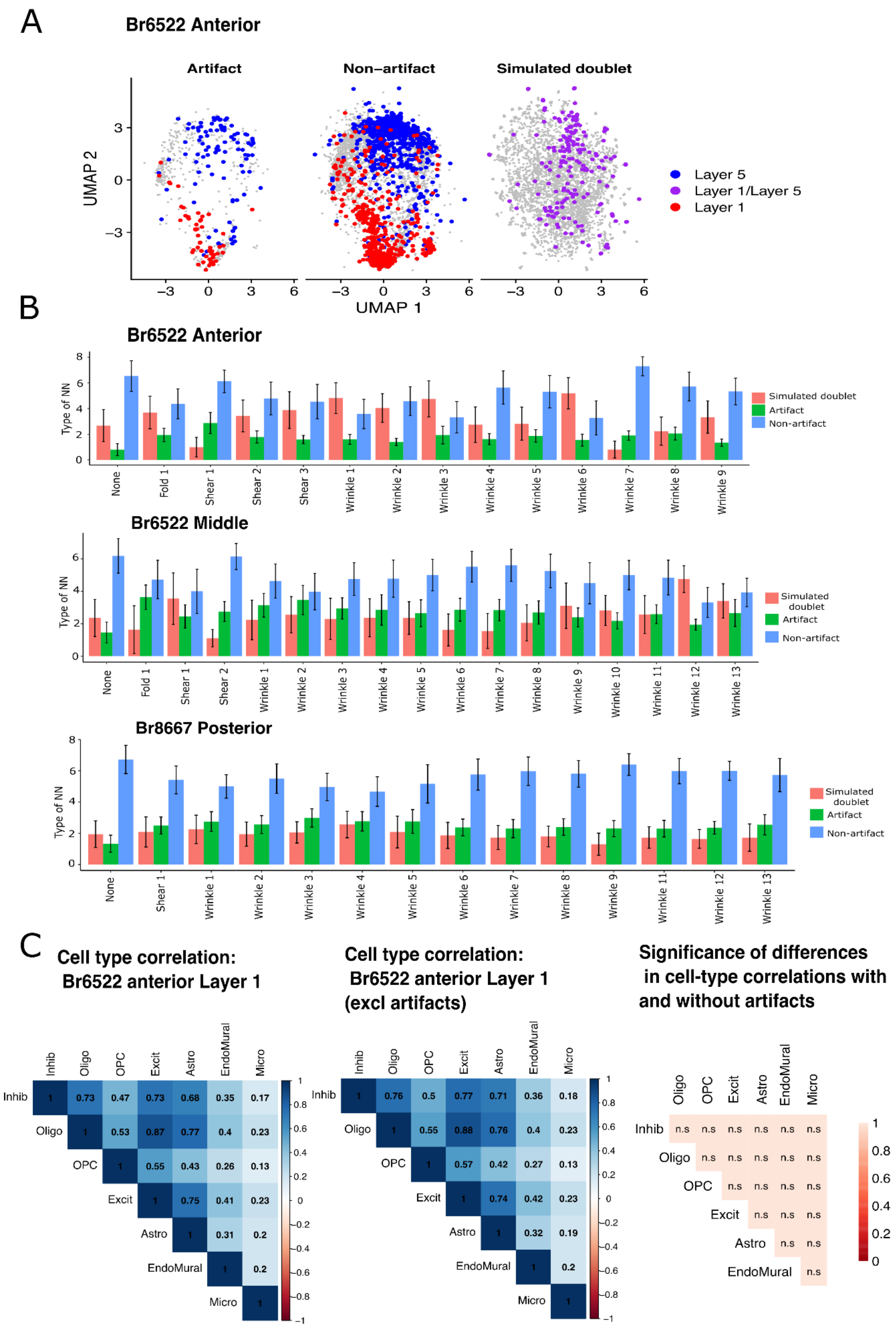
A
Br6522 Anterior
Artifact
Non−artifact
Simulated doublet
UMAP 2
UMAP 1
Layer 5
Layer 1/Layer 5
Layer 1
B
Br6522 Anterior
Type of NN
Simulated doublet
Artifact
Non-artifact
None
Fold 1
Shear 1
Shear 2
Shear 3
Wrinkle 1
Wrinkle 2
Wrinkle 3
Wrinkle 4
Wrinkle 5
Wrinkle 6
Wrinkle 7
Wrinkle 8
Wrinkle 9
Br6522 Middle
Type of NN
Simulated doublet
Artifact
Non-artifact
None
Fold 1
Shear 1
Shear 2
Wrinkle 1
Wrinkle 2
Wrinkle 3
Wrinkle 4
Wrinkle 5
Wrinkle 6
Wrinkle 7
Wrinkle 8
Wrinkle 9
Wrinkle 10
Wrinkle 11
Wrinkle 12
Wrinkle 13
Br8667 Posterior
Type of NN
Simulated doublet
Artifact
Non-artifact
None
Shear 1
Wrinkle 1
Wrinkle 2
Wrinkle 3
Wrinkle 4
Wrinkle 5
Wrinkle 6
Wrinkle 7
Wrinkle 8
Wrinkle 9
Wrinkle 11
Wrinkle 12
Wrinkle 13
C
Cell type correlation: Br6522 anterior Layer 1
Cell type correlation: Br6522 anterior Layer 1 (excl artifacts)
Significance of differences in cell-type correlations with and without artifacts
Inhib
Oligo
OPC
Excit
Astro
EndoMural
Micro
n.s

**Supplementary Figure 7**. **Identifying heterotypic artifact spots** (**A**) Joint UMAP embedding of observed non-artifact, artifact and simulated heterotypic spots. Heterotypic spots that were obtained as a linear combination of expression profiles from Layer 1 and Layer 5 form a continuous spectrum between two discrete layers and leads to a lack of separation between clusters specific to a biological layer. (**B**) Comparison of the number of spots which were non-artifacts, artifacts and simulated heterotypic doublets among the ten shared nearest neighbors of spots belonging to annotated tissue regions specifying the presence of an artifact. No significant and consistent differences were observed between non-artifact spots and artifact spots in the number of nearest neighbors that were artifacts. Therefore thresholding the fraction of shared nearest neighbors which were simulated artifacts would lead to a decrease in the number of non-artifact spots and not completely eliminate artifact spots. (**C**) Pearson correlation between 7 constituent cell types across spots belonging to sample Br6522_ant Layer 1 before and after excluding artifact spots computed from Tangram cell assignments. None of the differences in cell-type correlation were significantly different at FDR < 0.01 across all samples and layers.

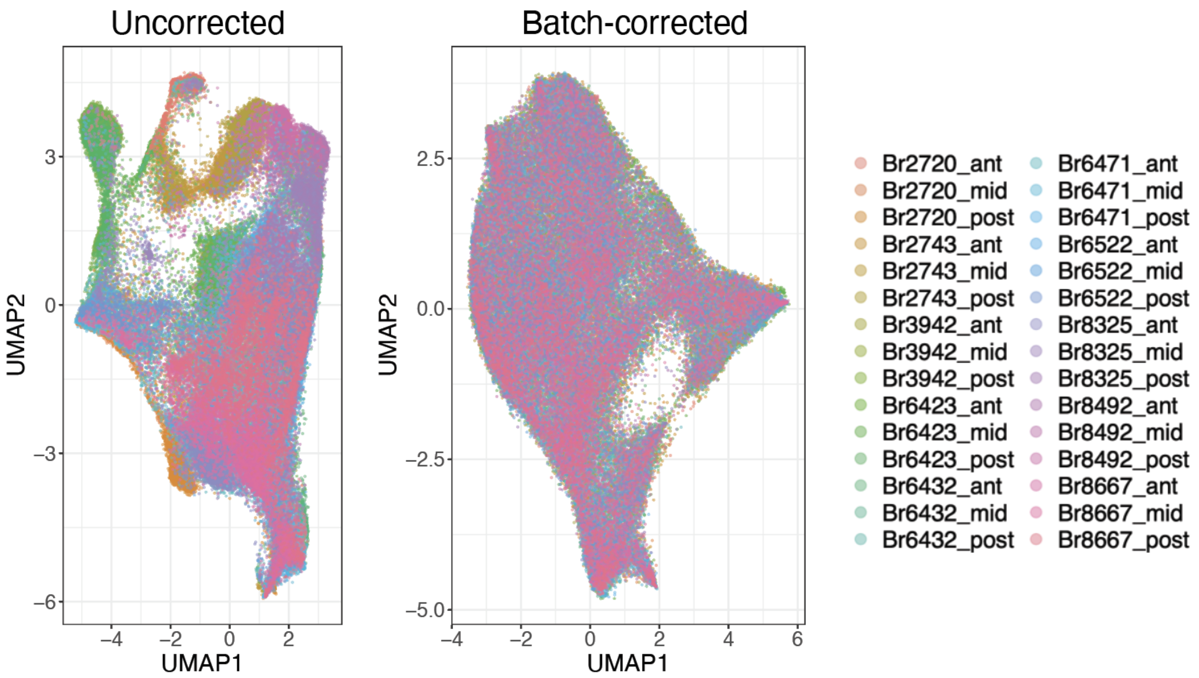


**Supplementary Figure 8. Batch correction of Visium data across samples.** UMAP of all spots before batch correction (left) and after batch correction (right) colored by sample ID. Batch correction with *Harmony (52)* reduces both donor and technical effects.

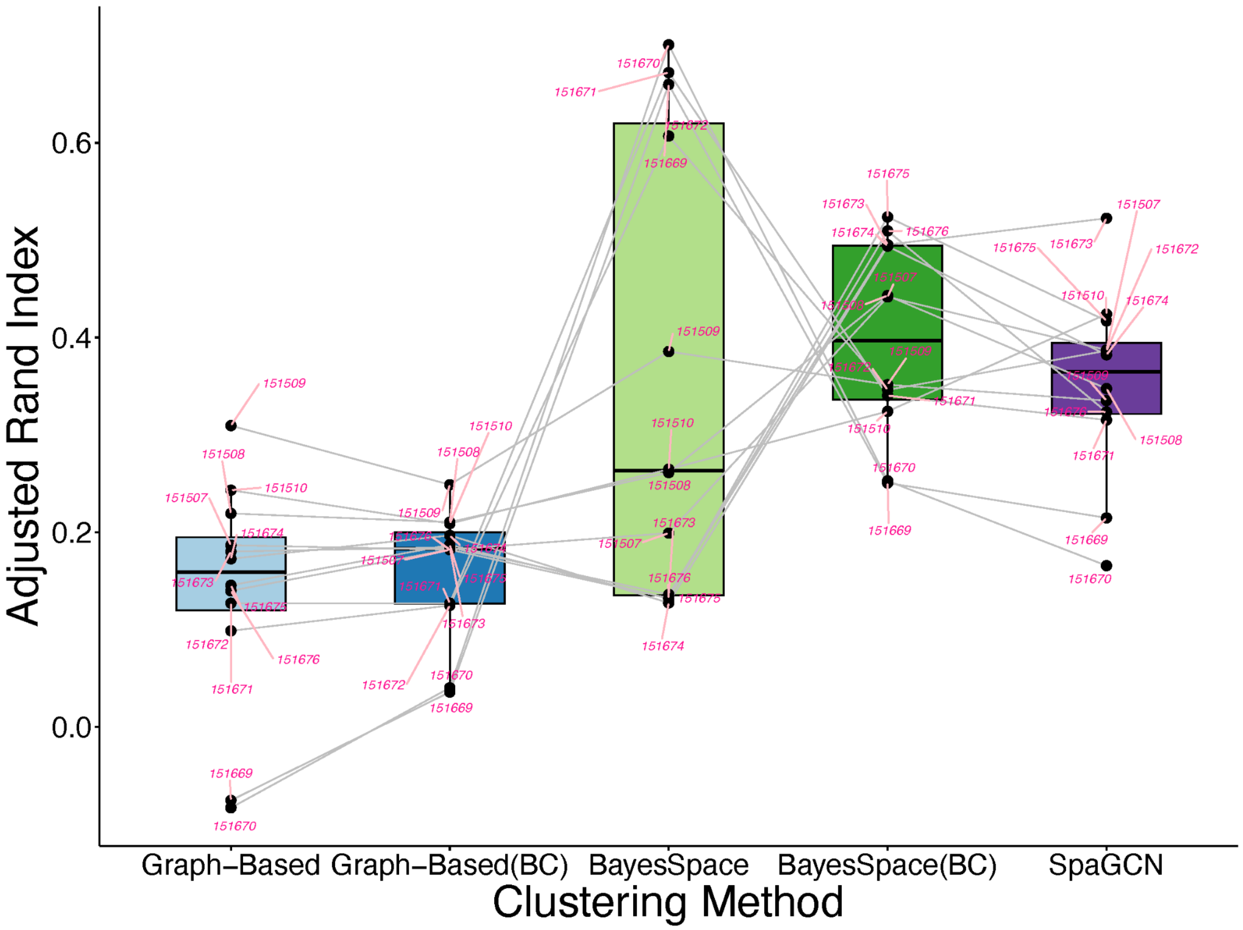


**Supplementary Figure 9**. ***BayesSpace* with batch correction computationally identifies laminar spatial domains with highest accuracy.** Summary of clustering accuracy in 12 samples from (6) using different clustering methods (graph-based, *BayesSpace*, *SpaGCN*) with or without batch correction (BC) with *Harmony* where applicable. *SpaGCN* does not perform clustering across samples and therefore batch correction could not be applied. Method performance was evaluated and compared using the Adjusted Rand Index (ARI), which measures the similarity between the predicted cluster labels and manually annotated cortical layers where larger numbers correspond to improved performance. Batch correction greatly improved the median clustering accuracy for the *BayesSpace* algorithm, which outperformed graph-based clustering and individual sample clustering with *SpaGCN*. Sample identifiers are shown in pink and connected across methods by gray lines. Boxes show the first, second (median), and third quartiles.

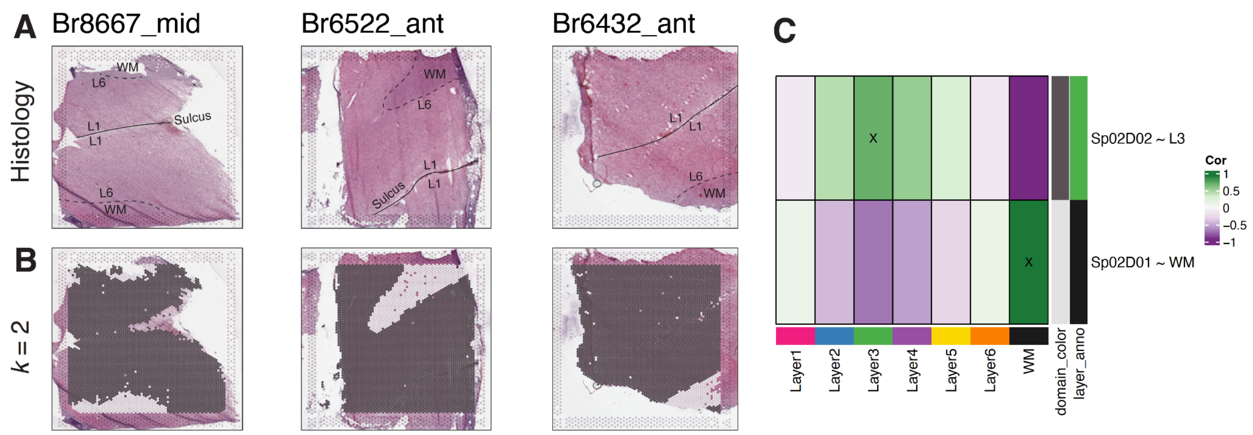


**Supplementary Figure 10**. **Clustering with *BayesSpace* at *k*=2 separates white matter and gray matter.** (**A**) H&E staining of three representative DLPFC sections from **Fig. 2** with annotations for white matter (WM) and Layers (L)1 and L6 in the gray matter. Sulci separating folds between gyri are marked with a solid line. (**B**) *BayesSpace* clustering at *k*=2 separates white matter ($Sp_2D_1$) from gray matter ($Sp_2D_2$). (**C**) Spatial registration heatmap depicting correlation between enrichment *t*-statistics computed on unsupervised $Sp_2Ds$ (x-axis) against those from manually annotated histological layers from (6) (y-axis). Higher confidence annotations ($\rho > 0.25$) are marked with an "X".

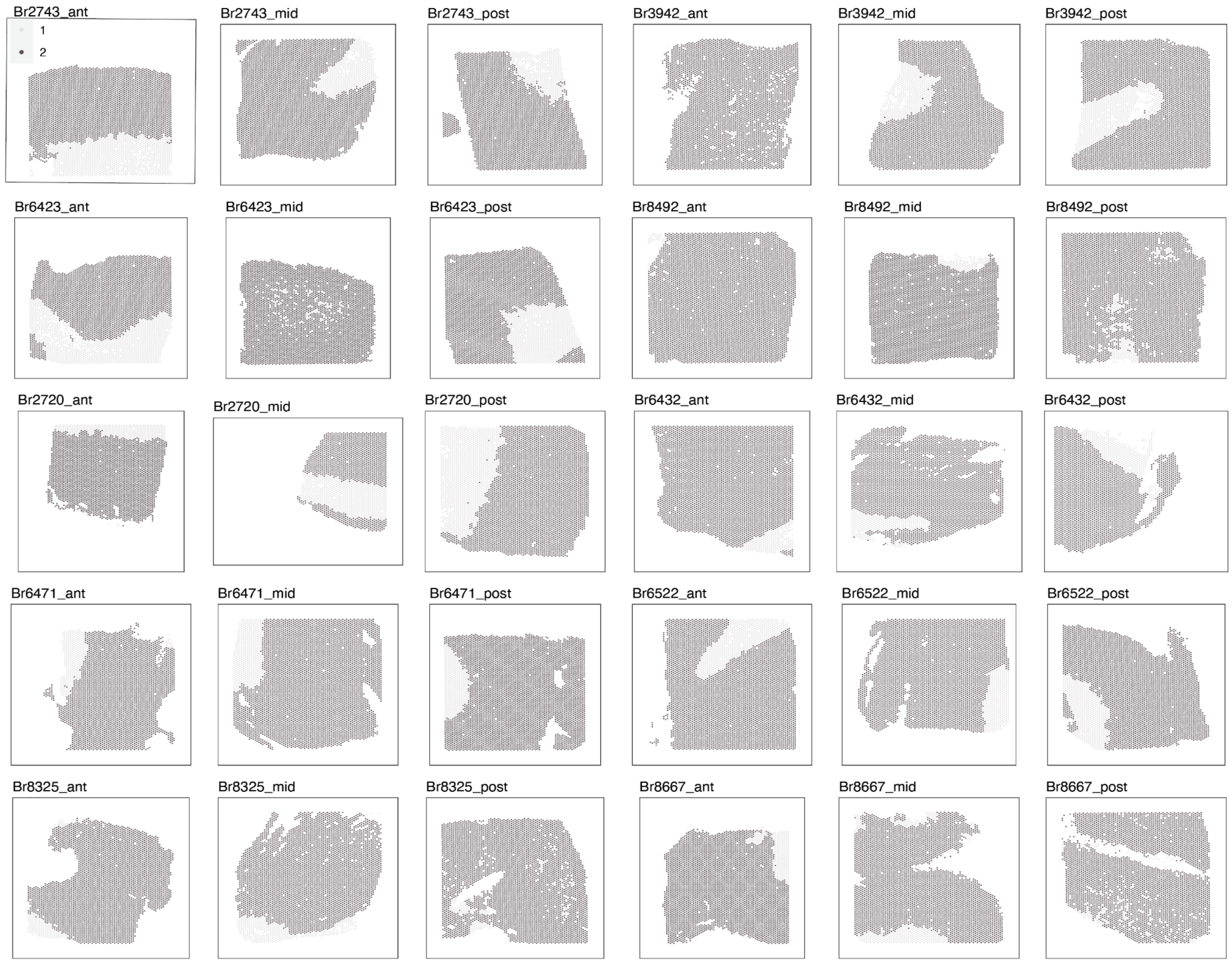


**Supplementary Figure 11**. **Unsupervised clustering with *BayesSpace k*=2 for all 30 DLPFC samples.** Spotplots depicting the cluster labels for spatial domains identified with *Bayespace k*=2 ($Sp_2Ds$), which associate with white matter (light gray) and gray matter (dark gray) as shown in **Fig S10**.

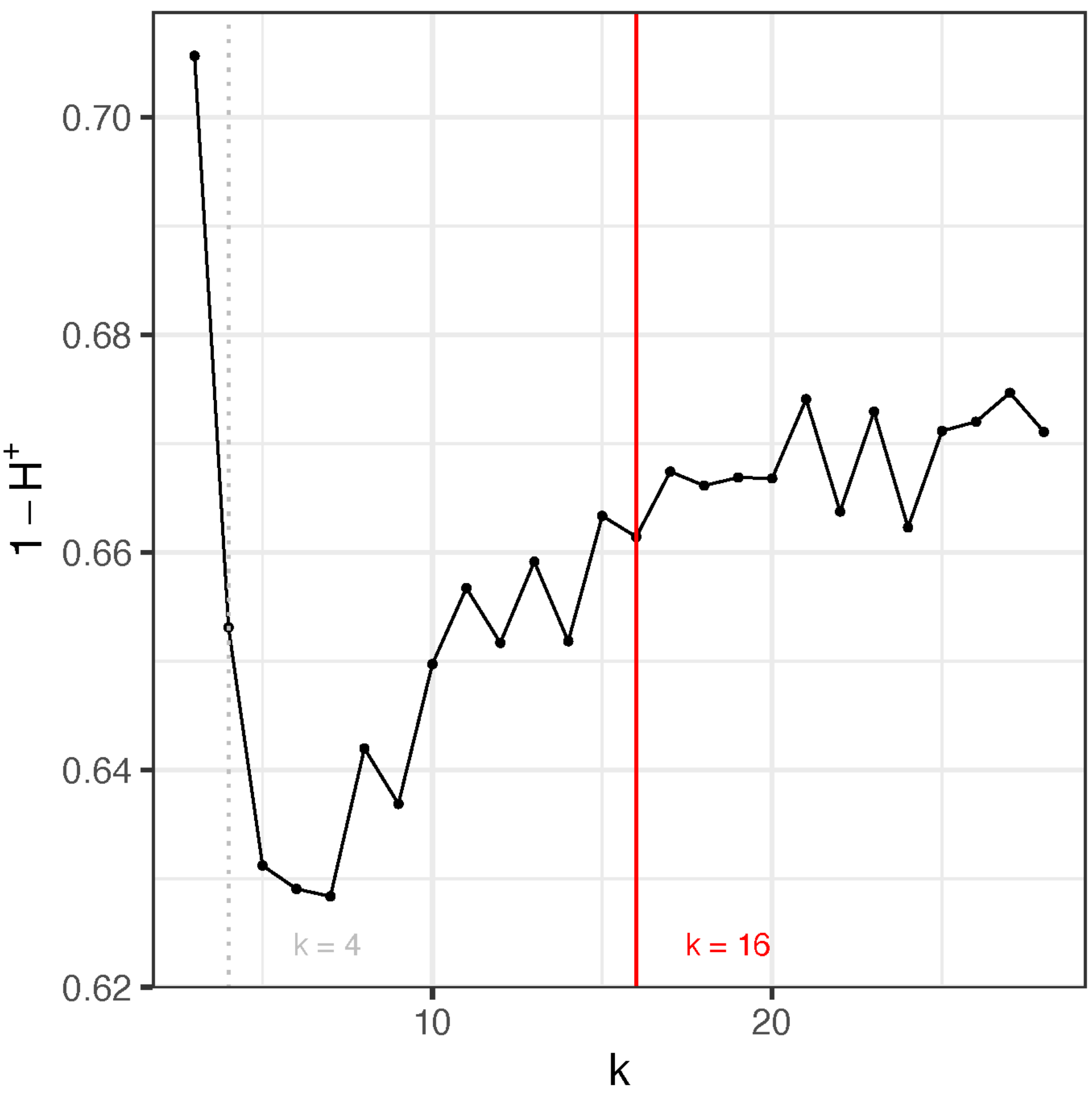


**Supplementary Figure 12. Data-driven identification of the optimal number of unsupervised clusters (k) in DLPFC samples using *fasthplus (H+)*.** To identify an optimal number of clusters, a discordance internal validity metric was used to assess clustering performance for different values of *k* (59). We plotted the 1 minus the discordance measure ($H^+$) for all values of *k* (4 to 28). The second inflection point at k=16 highlights the value of k at which discordance is decreasing at a negligible rate.

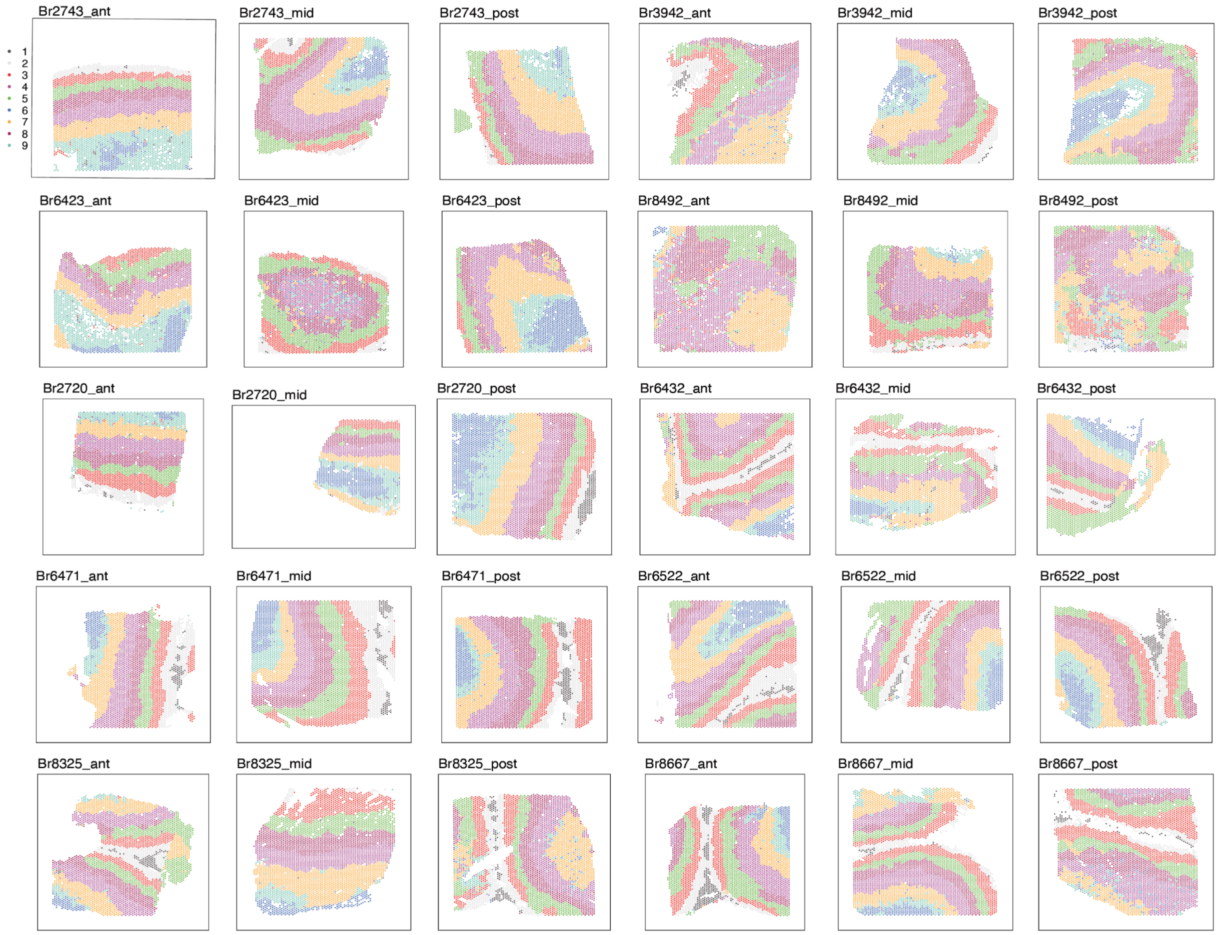


**Supplementary Figure 13. Unsupervised clustering with *BayesSpace k*=9 for all 30 DLPFC samples.** Spotplots depicting the cluster labels for spatial domains identified using *Bayespace k*=9 ($Sp_9Ds$), which most accurately recapitulated the classic histological layers.

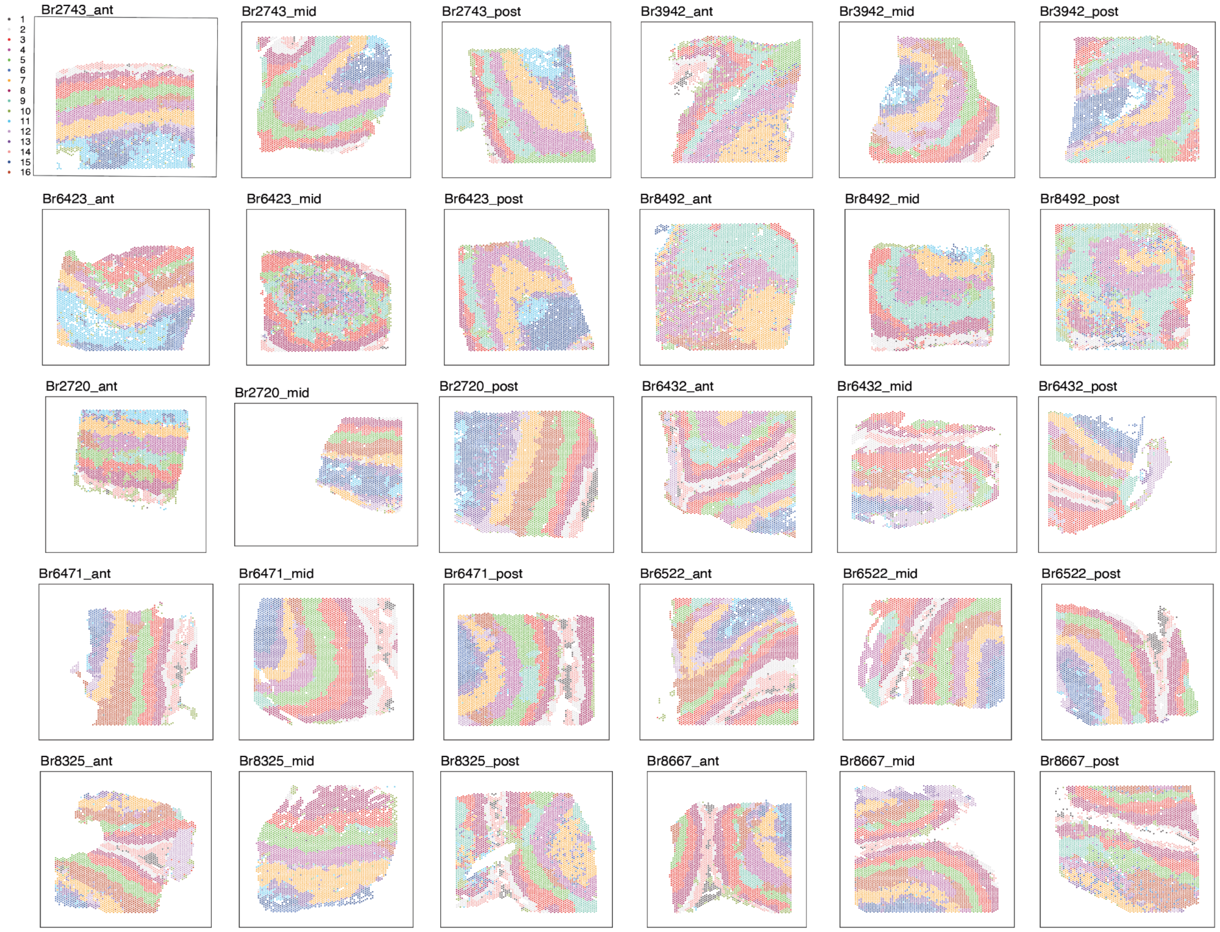


**Supplementary Figure 14. Unsupervised clustering with *BayesSpace k*=16 for all 30 DLPFC samples.** Spotplots depicting the cluster labels for spatial domains identified with *Bayespace k*=16 ($Sp_{16}Ds$), are highly correlated with multiple histological layers suggesting molecularly-defined sublayers.

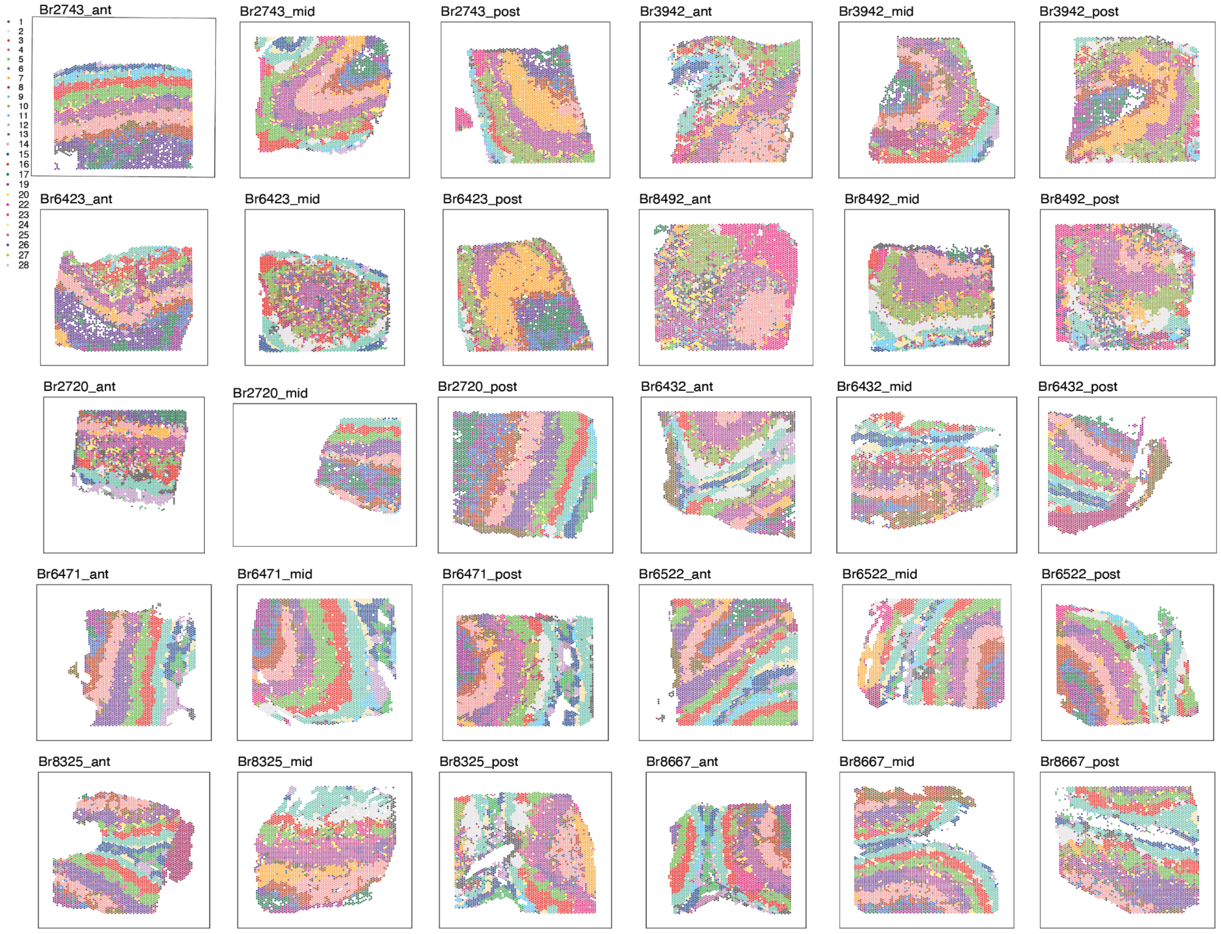


**Supplementary Figure 15. Unsupervised clustering with *BayesSpace k*=28 for all 30 DLPFC samples.** Spotplots depicting the cluster labels for spatial domains identified with *Bayespace k*=28 ($Sp_{28}Ds$), which represent both laminar and non-laminar domains. Note that clusters 18 and 21 are not assigned any spots, as `BayesSpace::spatialCluster(q = 28)` sets the maximum number of clusters, but does not guarantee returning that exact number of clusters.

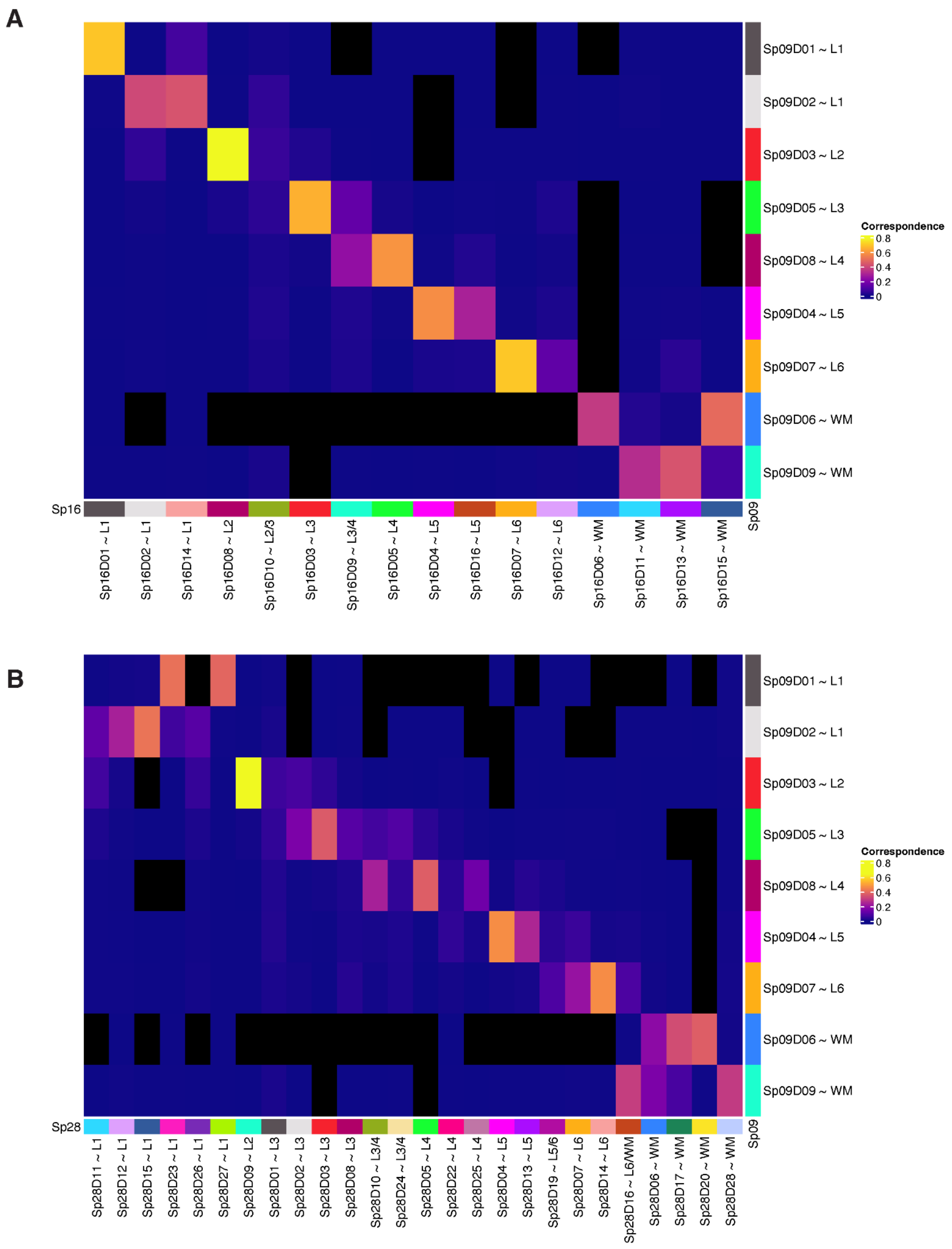


**Supplementary Figure 16. Relationship between *BayesSpace* spatial domains (SpDs) at different resolutions.** (**A**) Correspondence between $Sp_9$Ds and $Sp_{16}$Ds. (**B**) Correspondence between $Sp_{16}$Ds and $Sp_{28}$Ds. Correspondence was computed using *bluster* `linkClustersMatrix()` with default arguments, which is equivalent to the Jaccard Index. Spatial domains are labeled according to the spatial registration and annotation results from **Fig. 2**. Row and columns are ordered corresponding to the DLPFC histological layers. Values that are exactly 0 are shown in black to visually differentiate them from values close to 0.

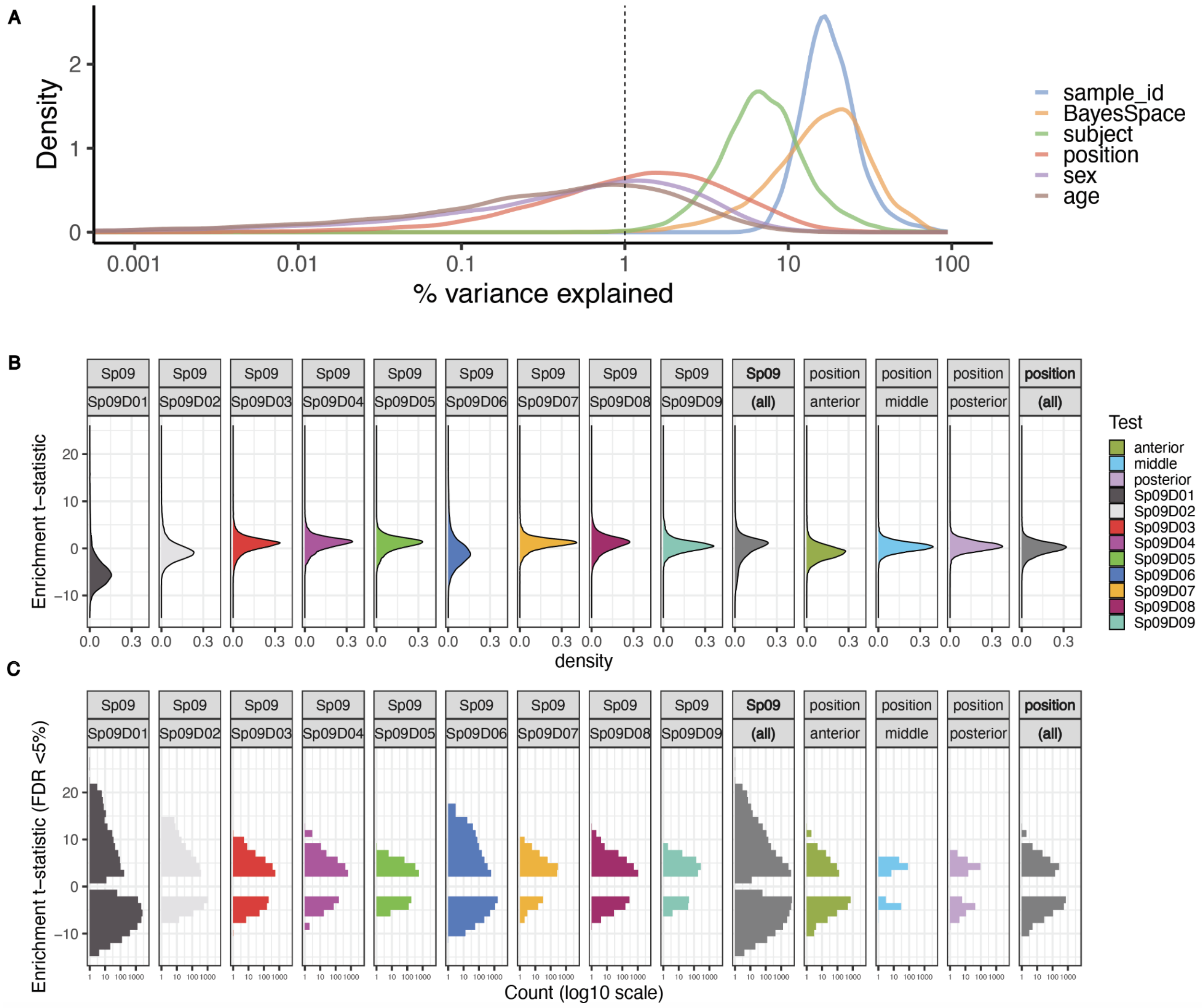


**Supplementary Figure 17**. **Summary of expression differences at $Sp_9$ resolution.** (**A**) Density curves for the percent of variance explained at the gene level for several variables at the $Sp_9$D resolution. The $Sp_9$Ds (*BayesSpace*, orange line) contribute the highest percent of variance and more than the anterior-posterior axis of the DLPFC (position, red line). (**B**) Densities of enrichment t-statistics for either each $Sp_9$D or each position compared to the rest of the same category. Overall position (all) t-statistics are closer to 0 than the $Sp_9$Ds. (**C**) Frequency histograms for the enrichment t-statistics for either each $Sp_9$D or each position, after subsetting to FDR<5%. While all tests have FDR<5% genes, there are far more for the $Sp_9$Ds than the position categories: 11,338 depleted (t<0) and 5,931 enriched (t>0) unique genes for $Sp_9$Ds compared with 1,402 depleted and 512 enriched unique genes across anterior-posterior position of the DLPFC.

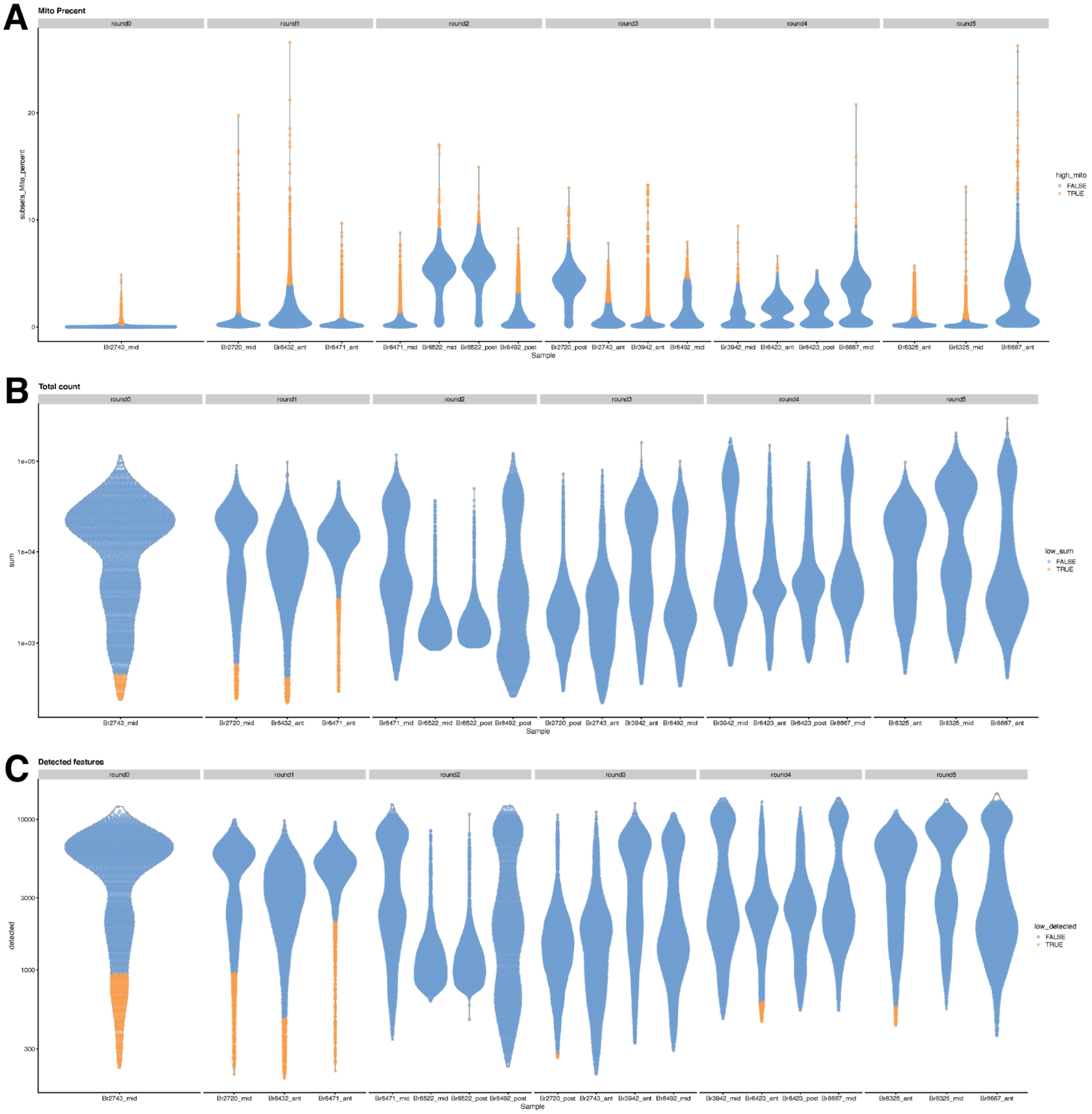


**Supplementary Figure 18. Assessment of quality control (QC) metrics across individual samples.** (**A**) Percent mitochondrial reads. (**B**) Total counts per nucleus. (**C**) Number of detected features per nucleus. Nuclei not passing a per-sample threshold set at 3 median-absolute-deviations away from median (thresholds are higher for A and lower for B and C) are labeled in orange and were dropped from analysis.

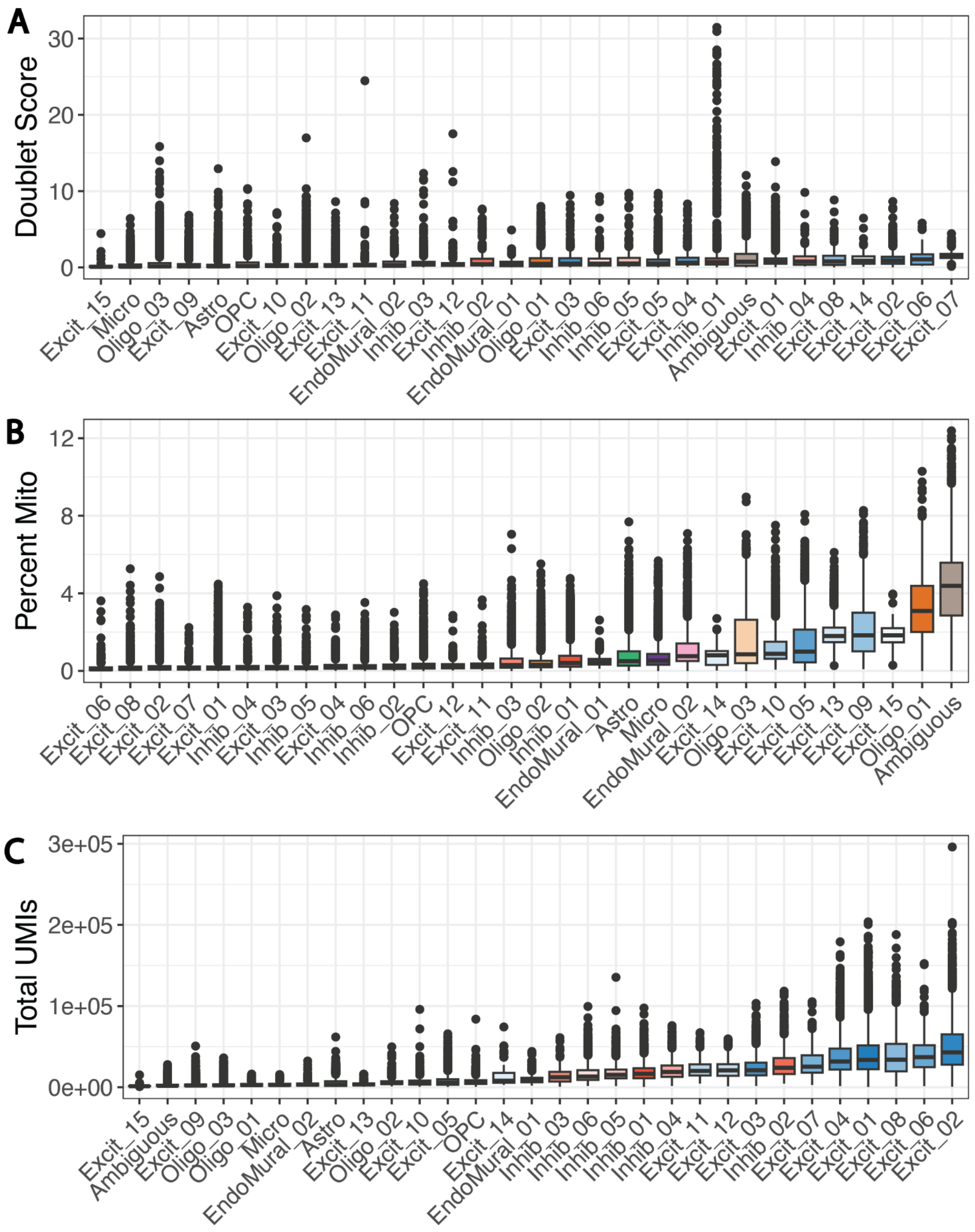


**Supplementary Figure 19. Assessment of quality control metrics across hierarchical clustering (hc) cell types.** (**A**) Doublet Scores. (**B**) Percent mitochondrial reads. (**C**) Total Unique Molecular Identifiers (UMIs) per nuclei. High percent mitochondrial genes and low total UMI's may have driven the Ambiguous cluster. No clusters were dropped based on these metrics.

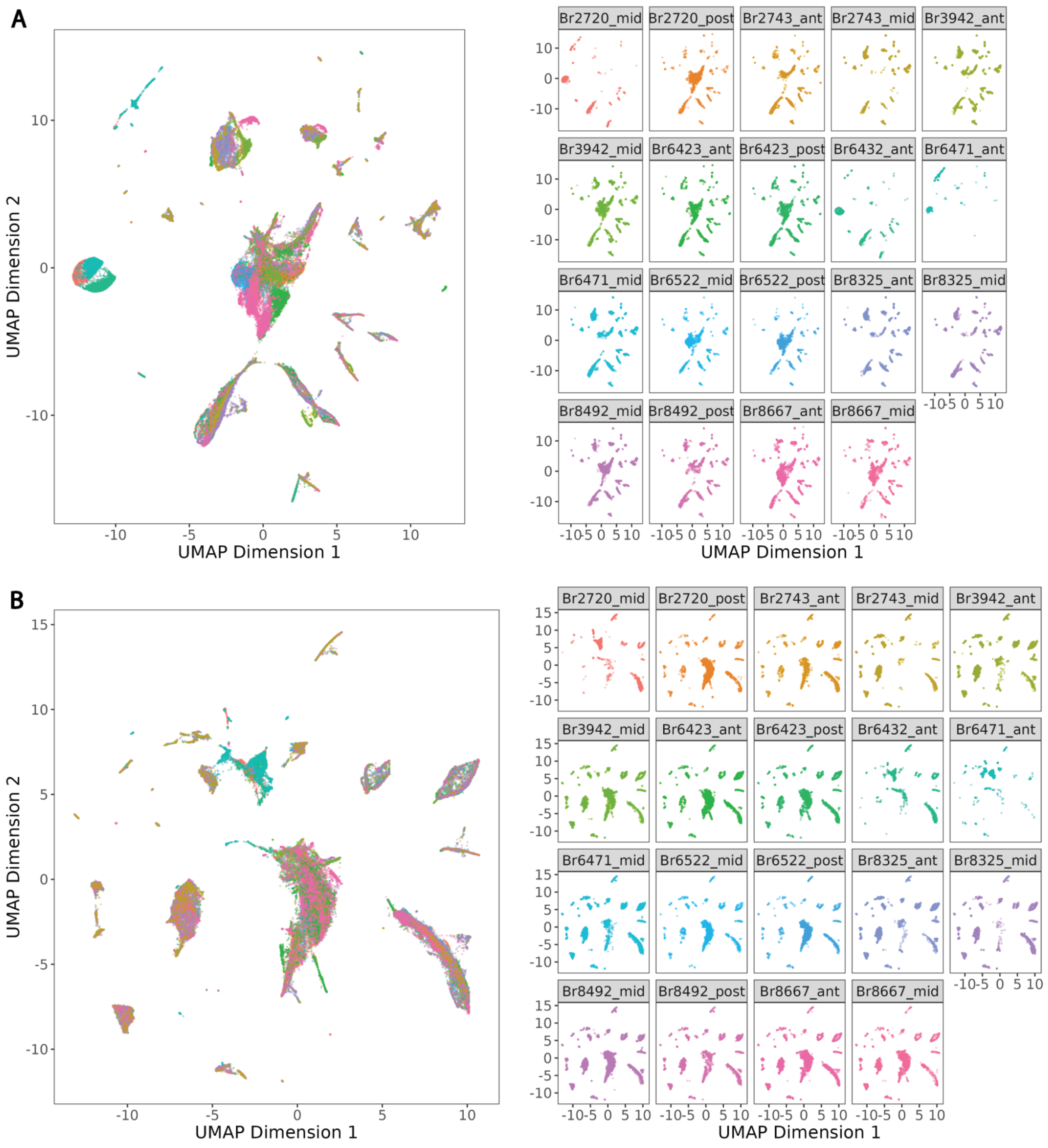


**Supplementary Figure 20. Batch correction of snRNA-seq data visualized by UMAP.** (**A**) UMAP of principal components (PCs) pre-batch correction colored by sample. (**B**) UMAP of PCs post-batch correction with *Harmony*, colored by sample. Batch correction reduced sample effect.

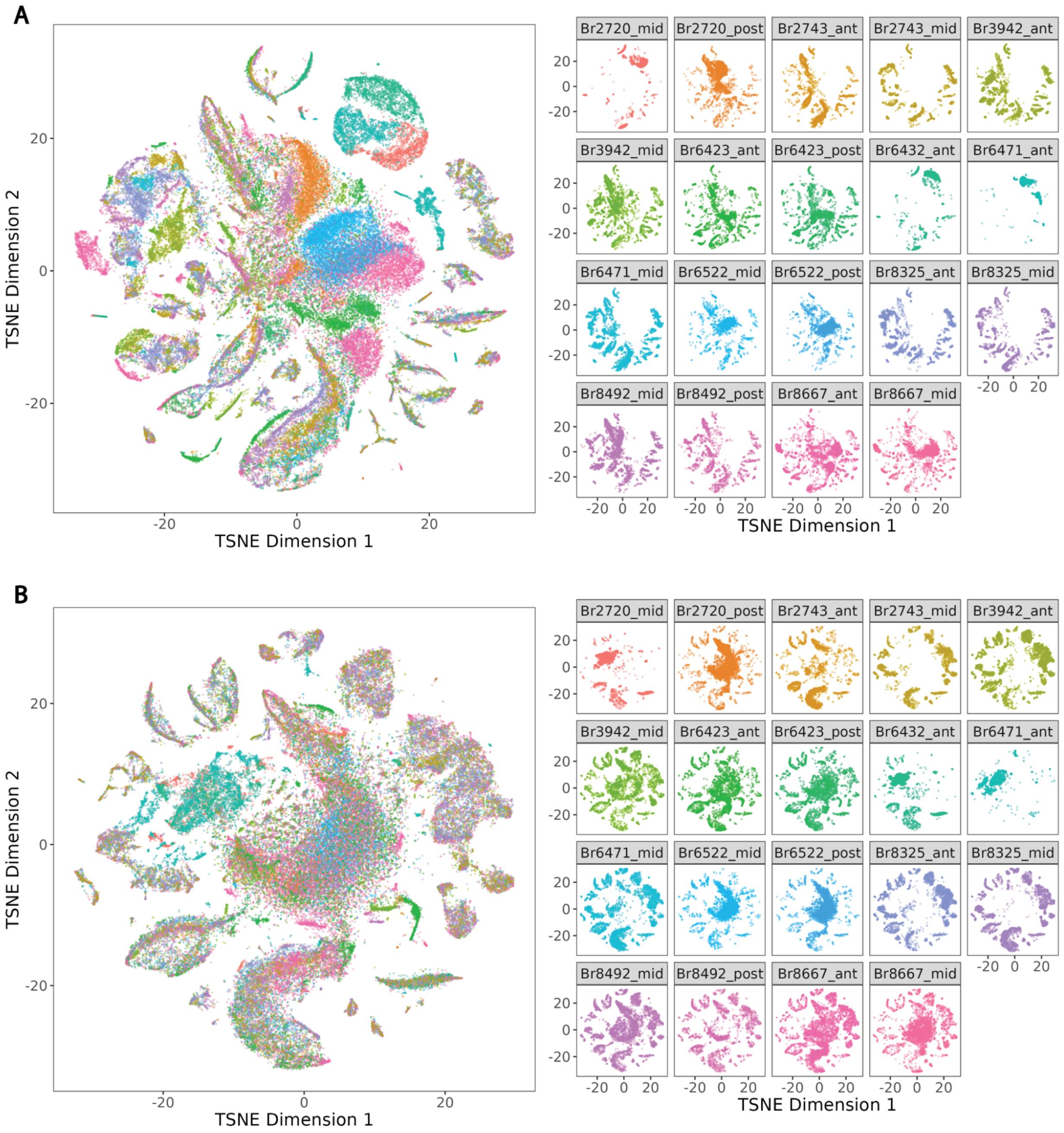


**Supplementary Figure 21. Batch correction of snRNA-seq data visualized by t-SNE.** (**A**) t-SNE of principal components (PCs) pre-batch correction colored by sample. (**B**) t-SNE of PCs post-batch correction with *Harmony*, colored by sample. Batch correction reduced sample effect.

A

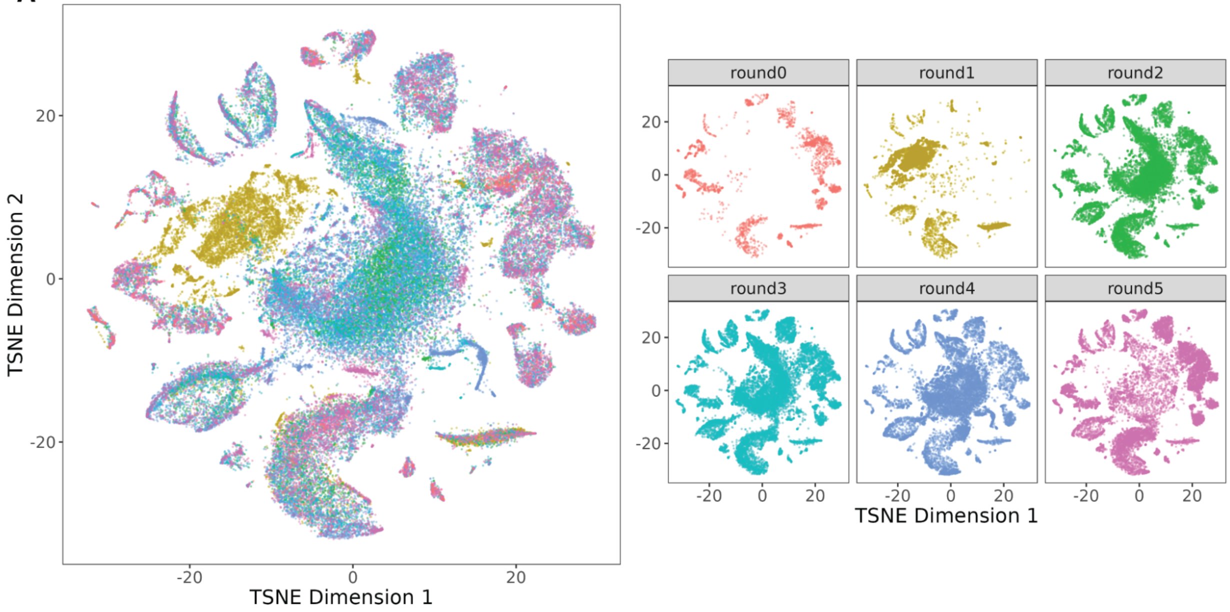


B

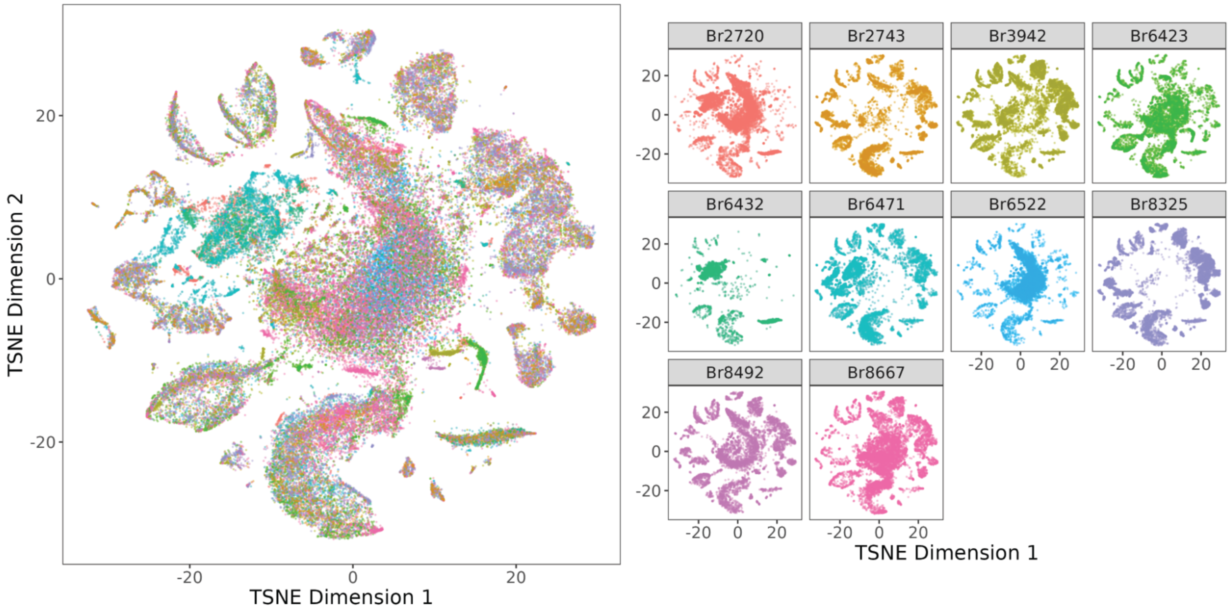


**Supplementary Figure 22. Batch correction of snRNA-seq data removes technical effects across rounds and donors.** (**A**) t-SNE of principal components (PCs) post-batch correction with *Harmony*, colored by sequencing round and (**B**) colored by donor.

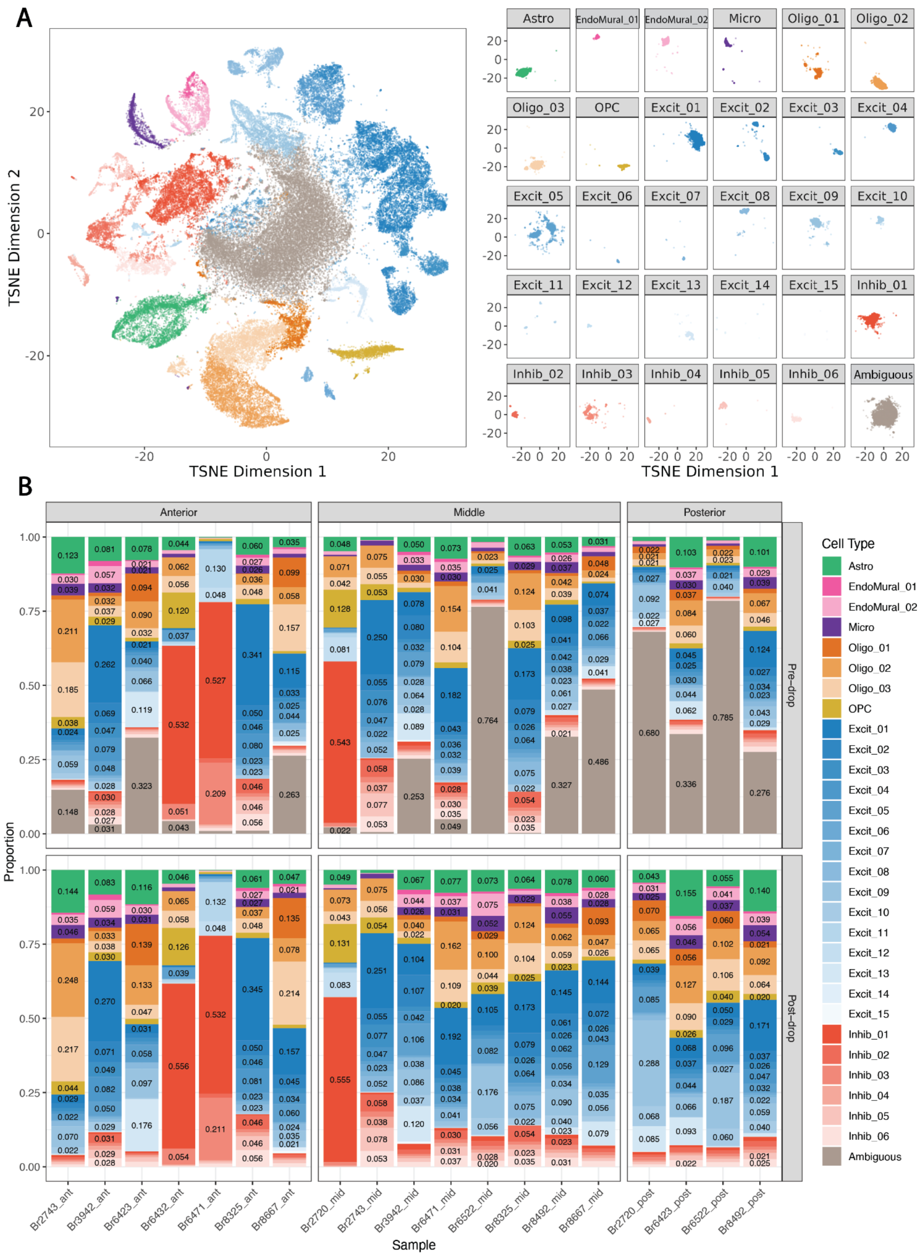


**Supplementary Figure 23. Identification of broad and fine resolution cell type clusters across anterior-posterior axis of DLPFC.** (**A**) t-SNE plot of 77,604 nuclei across 30 annotated clusters, including 29 identified cell types and one ambiguous cluster excluded from further analysis. Related to **Fig. 3** (**B**) Cell type proportion barplot by sample for hierarchical cluster (hc) cell type annotations, pre- and post- dropping 21,157 nuclei from ambiguous cluster.

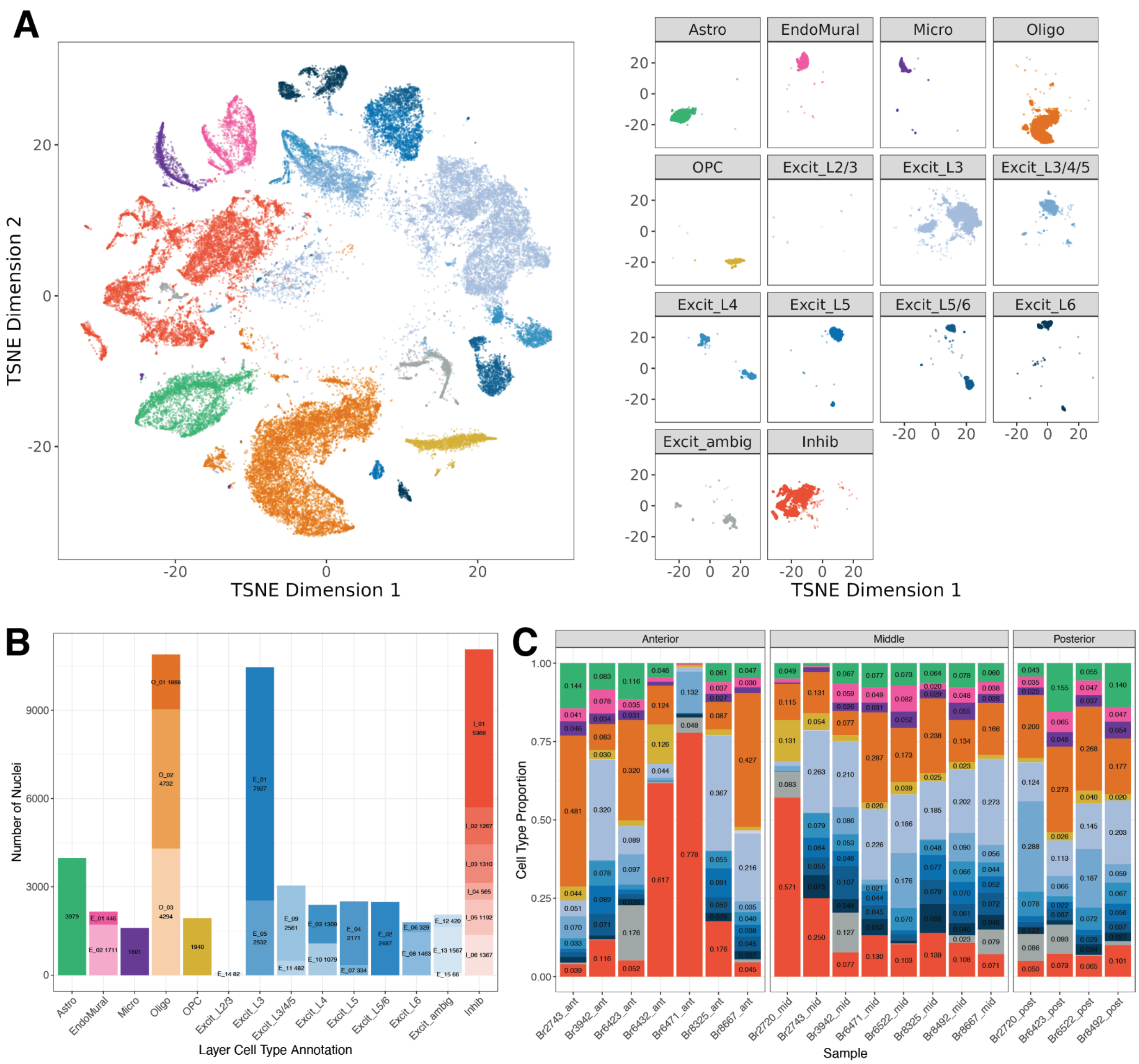


**Supplementary Figure 24. Layer-level annotation of snRNA-seq clusters** (**A**) t-SNE plot colored by layer-level annotation (related to **Fig. 3).** (**B**) Breakdown of the number of nuclei within each cluster at layer resolution, and relation to hierarchical cluster (hc) annotation. (**C**) Cell type composition barplot of each sample at the layer-level annotation.

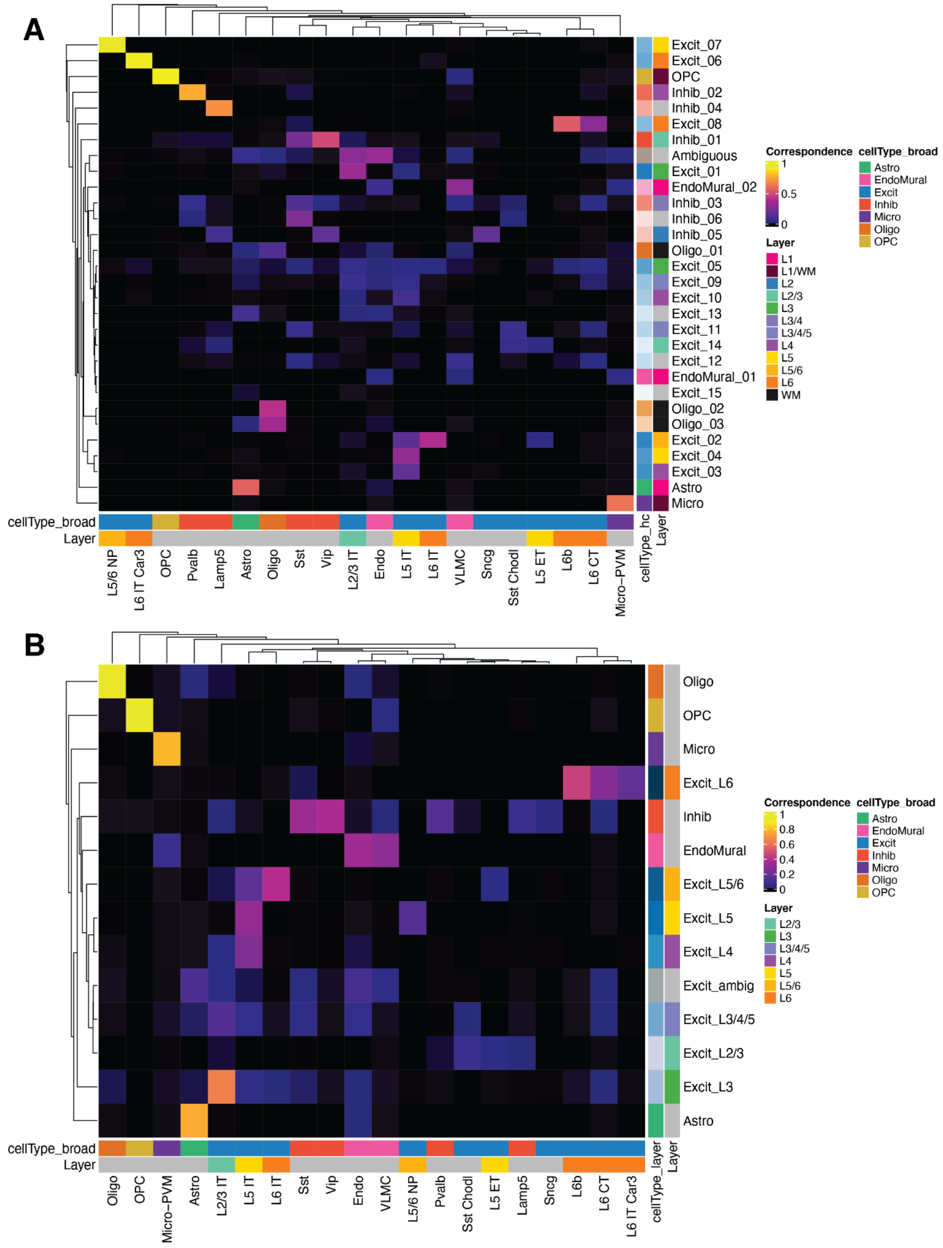


**Supplementary Figure 25. Comparison of cell clusters annotated using reference-based tool *Azimuth* versus hierarchical clustering (hc).** Correspondence of cell clusters annotated using a reference dataset of human motor cortex with *Azimuth* tool (20,21) versus cell type clusters at the (**A**) hc resolution (right annotation color bar shows hc cell type and layer associations annotated during spatial registration) and (**B**) layer-level resolution (right annotation bar shows layer-level cell type, and associated layer assignments for Excit cell types). Correspondence was computed using *bluster* `linkClustersMatrix()` with default arguments, which is equivalent to the Jaccard Index. Values that are exactly 0 are shown in black to visually differentiate them from values close to 0. Bottom annotation bar shows the broad cell type category for the *Azimuth* cell types and associated layer annotations for both plots.

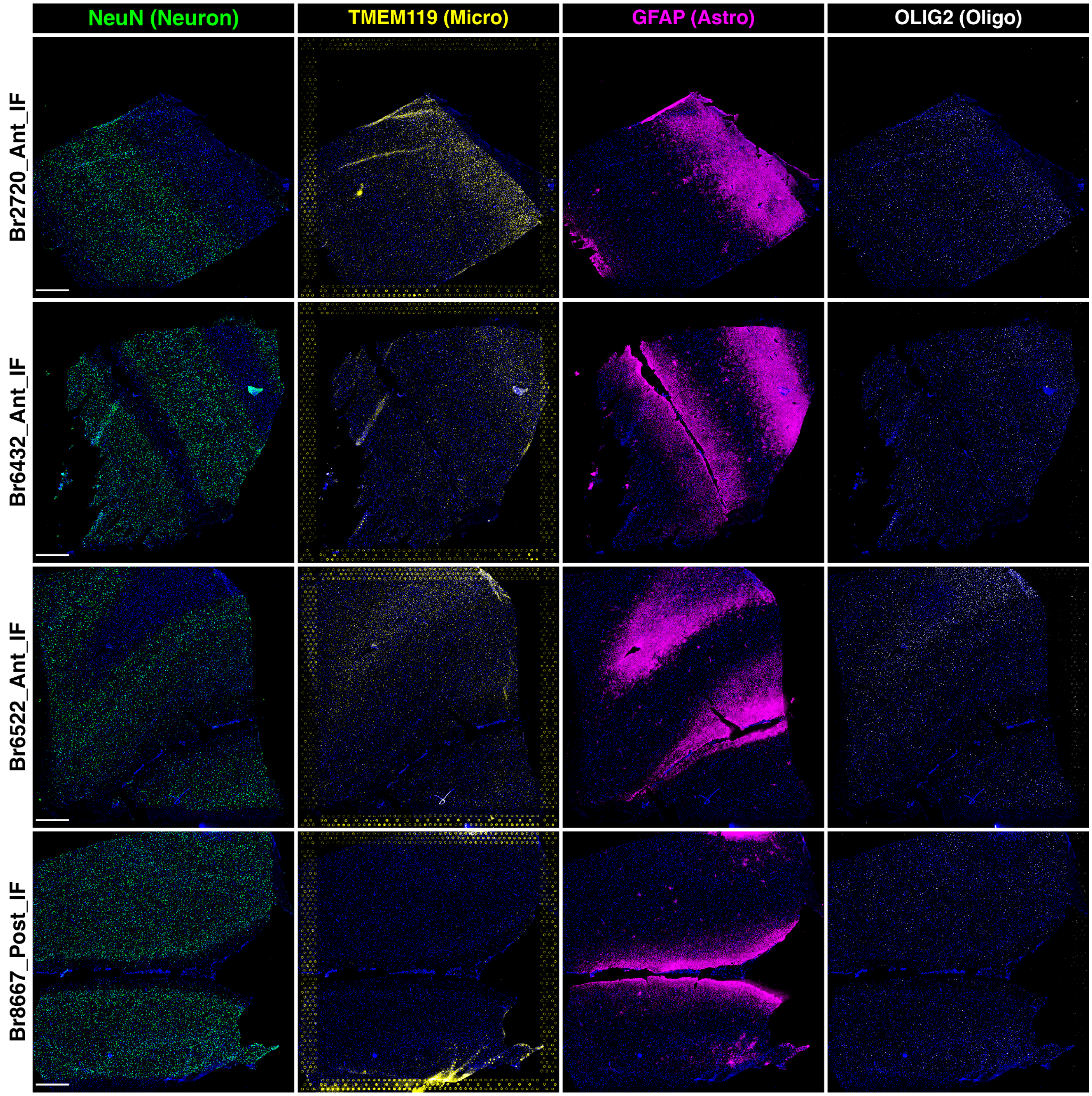


**Supplementary Figure 26**. **Multiplex immunofluorescent staining of DLPFC tissue sections in Visium-SPG assay.** Rows depict a DLPFC tissue section from each of 4 individual donors. Columns show individual fluorescent channels staining for proteins marking 4 broad cell types, including NeuN (labeling neurons), TMEM119 (labeling microglia), GFAP (labeling astrocytes), and OLIG2 (labeling oligodendrocytes). Nuclear DAPI staining is depicted in blue. Scale bar is 1mm.

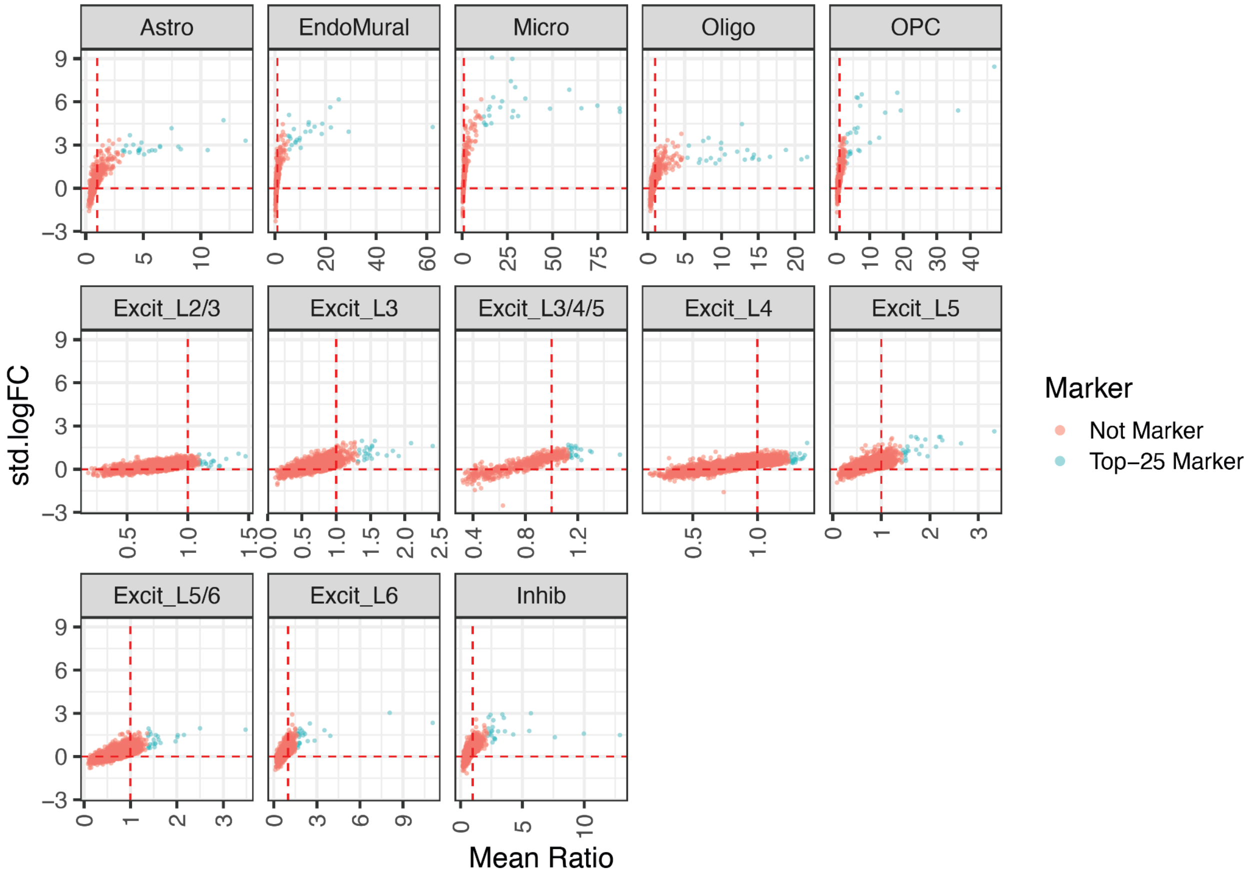


**Supplementary Figure 27. Distribution of standard log-fold change vs. mean ratio of all genes for layer-level cell types.** For each cell type, blue points are the 25 genes with the largest mean ratios, which were chosen as marker genes and later used as input into spot-deconvolution methods. Red points are the remaining genes, which are not considered cell type markers and therefore not used in downstream analyses.

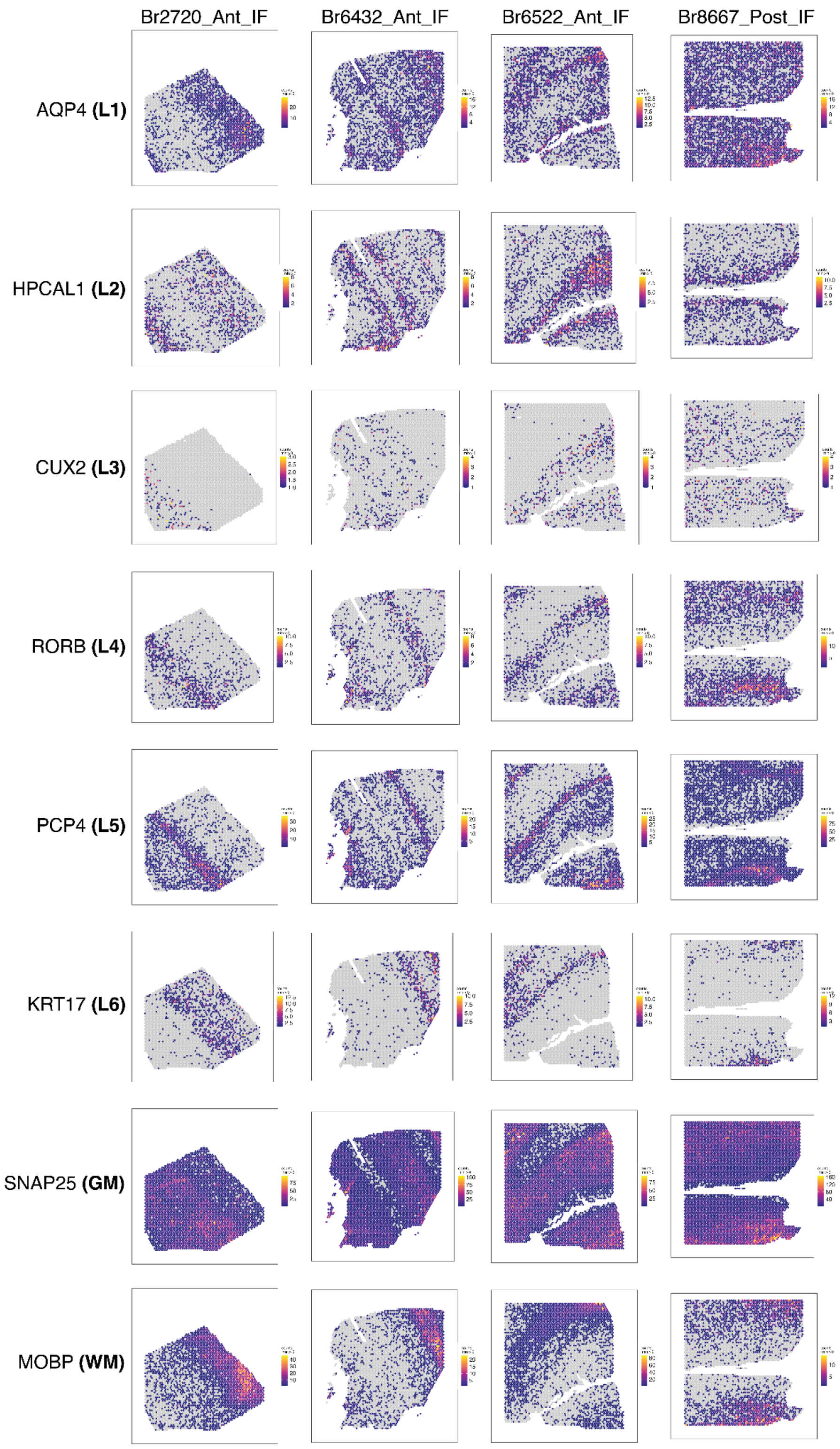


**Supplementary Figure 28. Spatial expression of DLPFC layer marker genes in the four Visium-SPG tissue sections.** Spotplots depicting expression (raw counts) for previously identified layer marker genes in the DLPFC (6) confirming the spatial orientation and laminar integrity of Visium-SPG data.

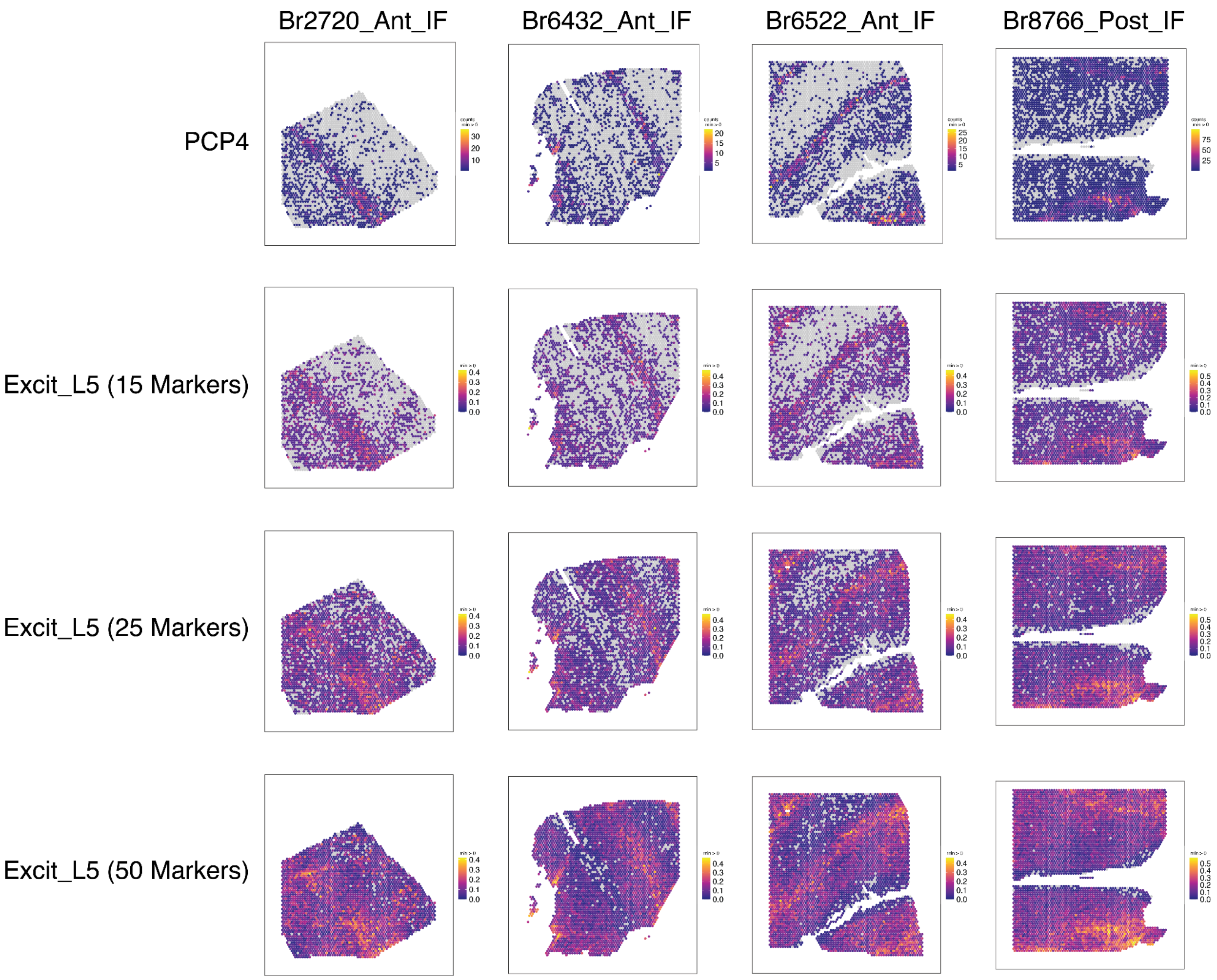


**Supplementary Figure 29. Determining the optimal number of gene markers for spot deconvolution.** Comparison of *PCP4* counts to L5 excitatory neuron marker gene expression with increasing number of markers. The top row shows spotplots for raw counts of *PCP4*, a classic marker for cortical L5, for all four Visium-SPG sections. Remaining rows show spotplots depicting the proportion of Excit_L5 marker genes with non-zero counts at each spot, when selecting the top 15, 25, or 50 marker genes. The choice of 25 marker genes per cell type results in an informative signal without significant expression outside of the expected layer. In (**Fig. 4B**), proportion of the top 25 markers with nonzero expression are compared against software-estimated Excit_L5 counts.

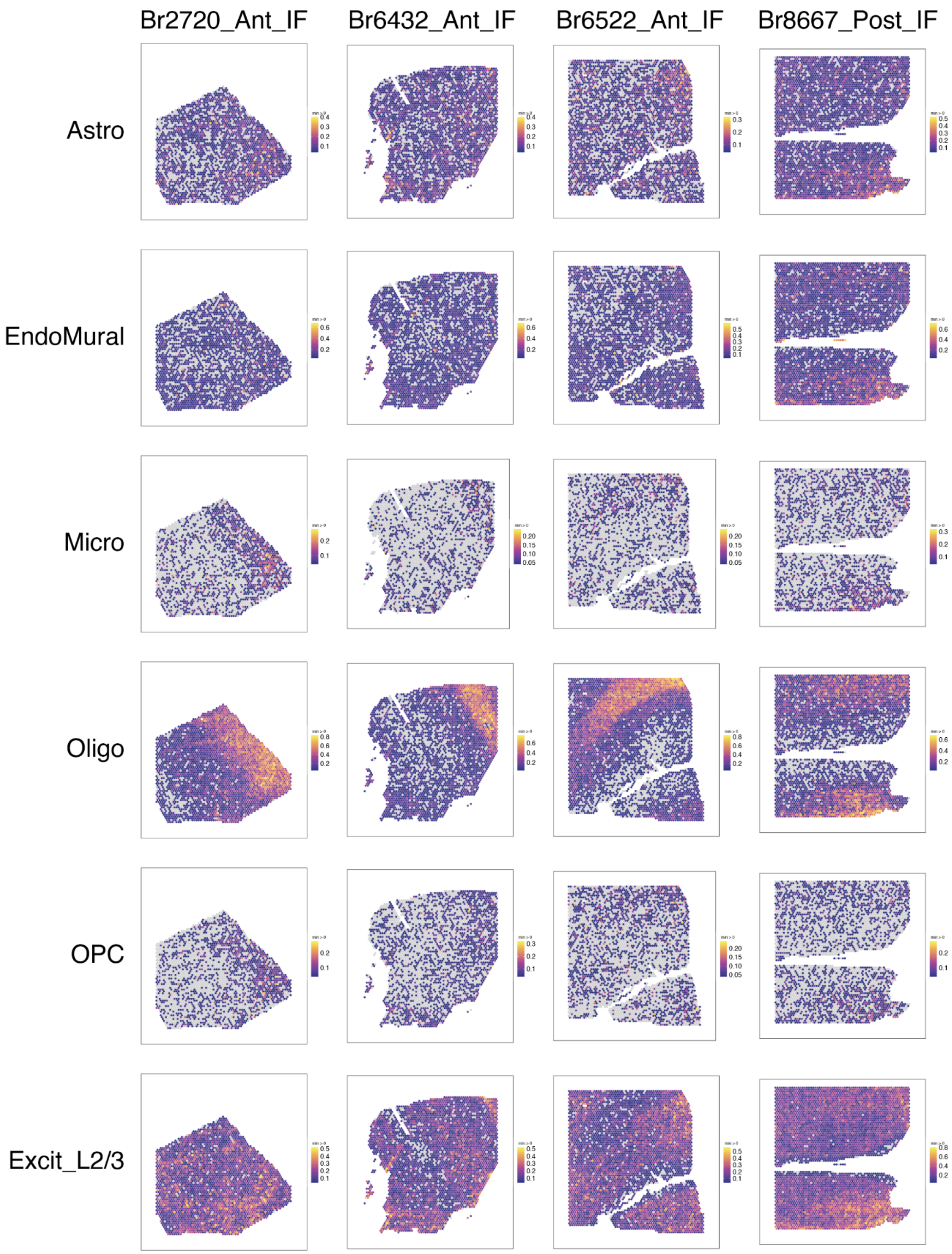
Br2720_Ant_IF
Br6432_Ant_IF
Br6522_Ant_IF
Br8667_Post_IF
Astro
EndoMural
Micro
Oligo
OPC
Excit_L2/3

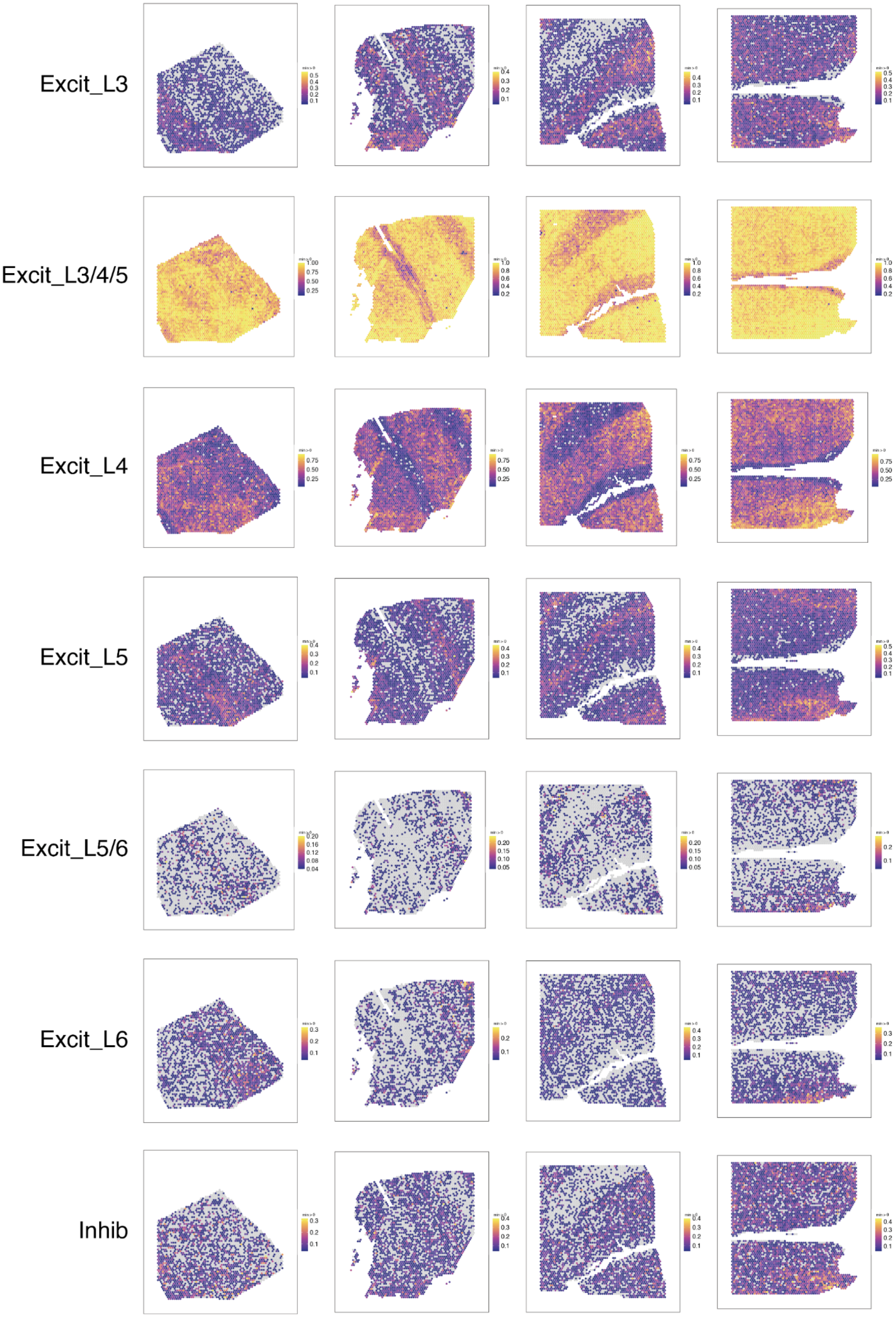
Excit_L3
Excit_L3/4/5
Excit_L4
Excit_L5
Excit_L5/6
Excit_L6
Inhib

**Supplementary Figure 30. Spatial validation of marker genes for layer-level cell types.** Proportion of marker genes with non-zero expression for layer-level cell types in the four Visium-SPG samples. In a given spot, the proportion of the top 25 marker genes for each cell type having at least one count was computed. This analysis was performed to validate the spatial integrity of marker genes. For example, as expected, the largest proportion of oligodendrocyte marker genes is in white matter.

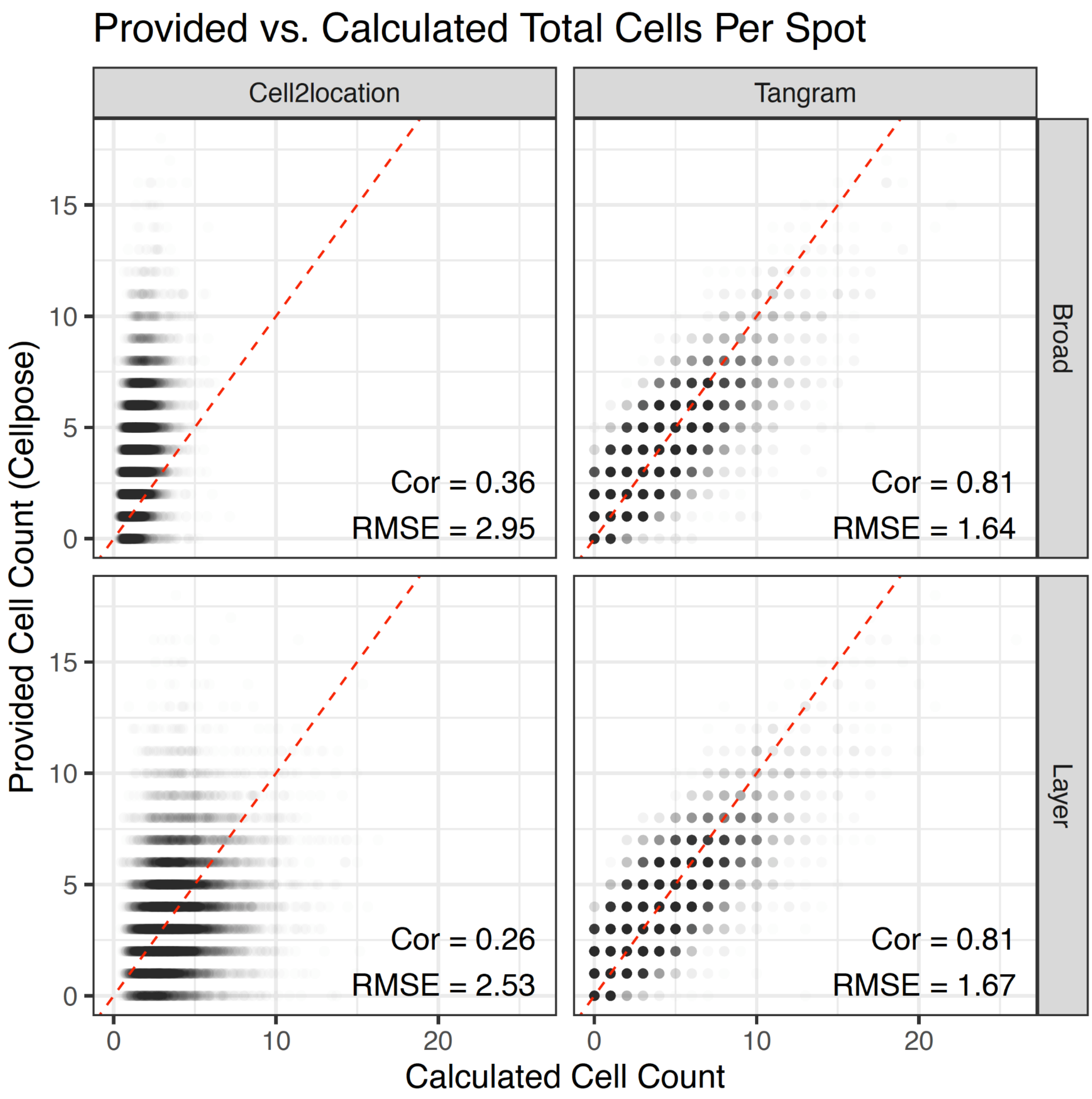


**Supplementary Figure 31. Comparison of provided versus calculated cell counts for *Tangram* and *Cell2location*.** Scatterplot of output vs. input cell counts per spot for *Tangram* and *Cell2location* when performing spot deconvolution at broad and layer-level cell type resolutions. Both software methods use an input number of cells per spot to guide deconvolution, but are not constrained to exactly match this number when outputting cell counts. For example, a spot reported by *Cellpose* to contain two cells may be assigned three cells by *Tangram*'s output. For our data, *Tangram* appears to more closely match input cell counts by spot compared to *Cell2location* as indicated by higher correlation and lower RMSE values. *SPOTlight* is excluded from this analysis as it outputs cell type proportions rather than counts, and consequently is unable to distort input counts as observed with *Tangram* and *Cell2location*.

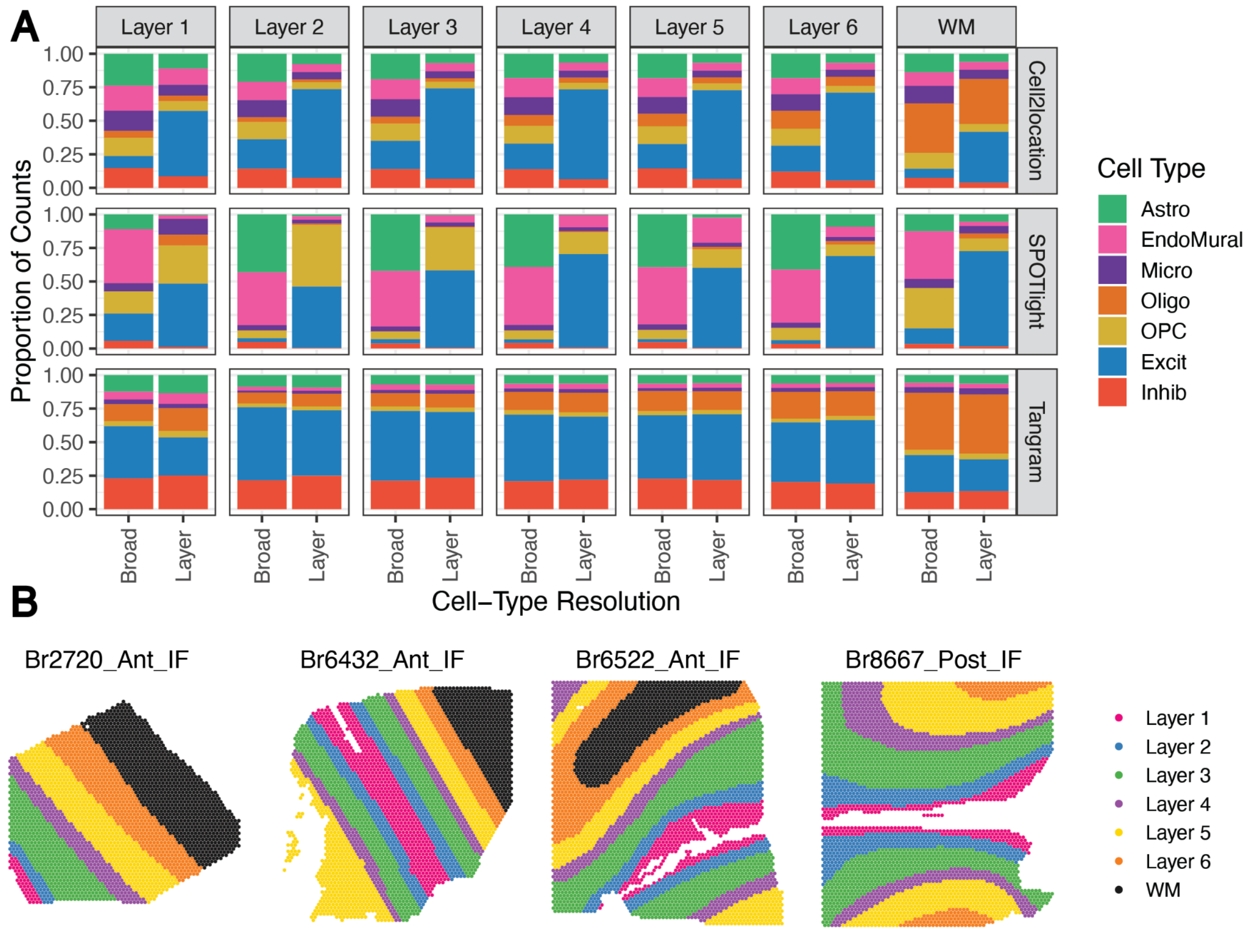


**Supplementary Figure 32. Distribution of cell type proportions across manually annotated histological layers in Visium-SPG data.** (**A**) For each histological layer, comparison of cell type proportions between broad and layer-level cell type resolutions. Layer-level excitatory cell types were collapsed onto broad resolution (i.e. Excit_L5, Excit_L6, etc were collapsed to Excit) and the proportion of each cell type was determined for each manually annotated layer. An average was taken across all four Visium-SPG sections, weighting each section equally to obtain proportion values that sum to 1.0. *Tangram* shows the most consistent distribution of cell types by layer between cell-type resolutions, which is expected as there should theoretically be a perfect match between broad and layer-level proportions. (**B**) Manual annotation of DLPFC layers for all four Visium-SPG sections. This annotation was also leveraged to benchmark spot-deconvolution software more thoroughly for layer-level cell types (**Fig. 4C**).

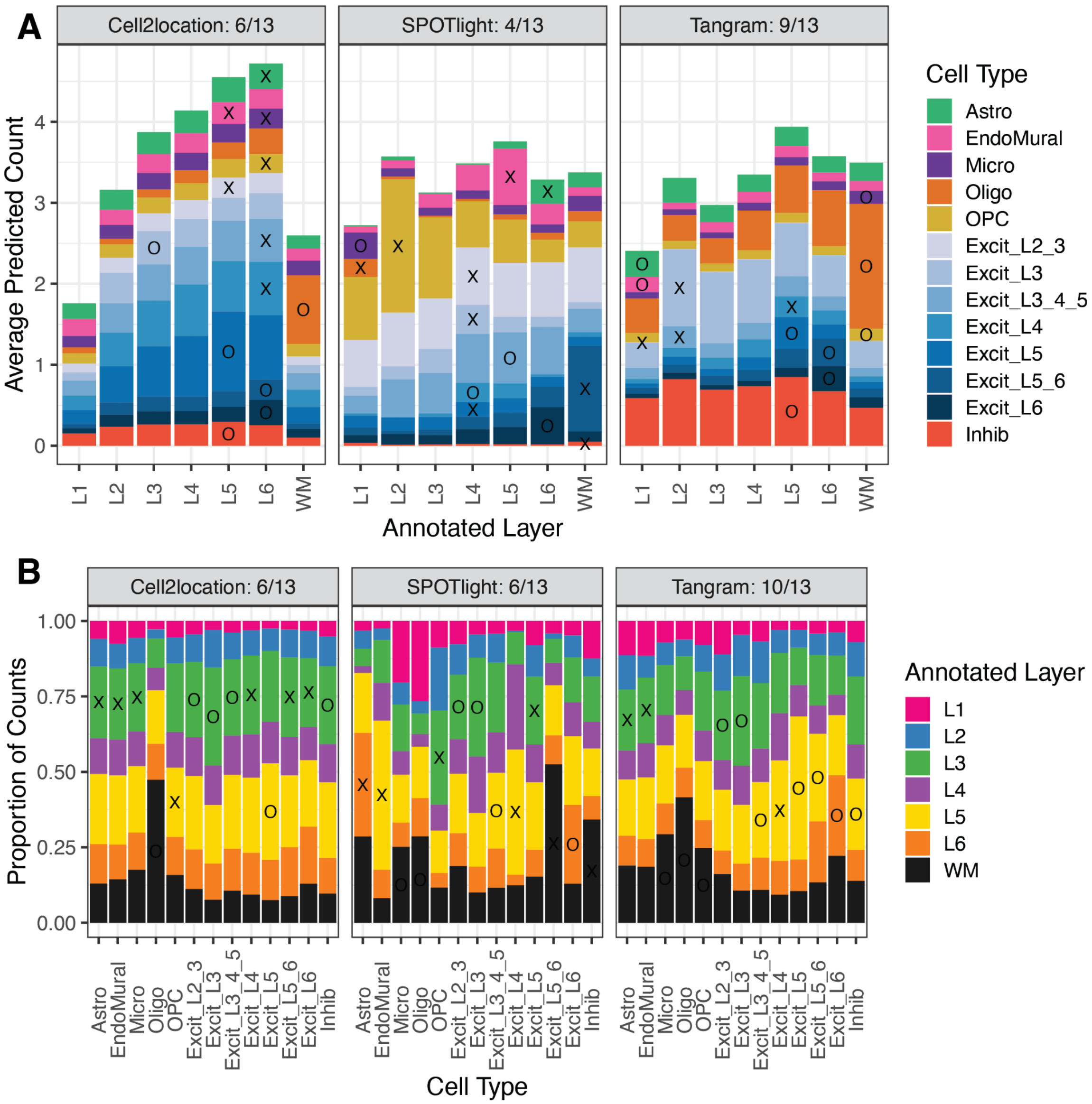


**Supplementary Figure 33**. **Distribution of layer-level cell types across manually annotated histological layers**. (**A**) Average predicted counts of each layer-level cell type by histological layer. In each histological layer, software-predicted counts were averaged across spots manually annotated to that layer, and then averaged across all four Visium-SPG sections. For each method and cell type, an “X” or “O” is placed in that layer with maximal average count. An “O” is placed when the cell type has a maximum in the expected layer (i.e.“Excit_L3” having the highest count in L3); an “X” is placed otherwise. The number of “O”s is summed across each method, and printed in the facet titles, to provide a single metric for the spatial accuracy of cell-type distributions by layer. A similar analysis was performed, but normalizes for cellular density across layers (**Fig. 4C**). (**B**) Proportion of counts belonging to each manually annotated layer by layer-level cell type. Within each column (cell type), the size of each colored bar indicates the fraction of all cells (averaged across all four Visium-SPG sections) that belongs to each manually annotated layer. As in (A), “X’s” and “O’s” are placed once per column for each cell type in the layer with maximal value, and “O”s are tallied for each method to provide a summary score in the facet title.

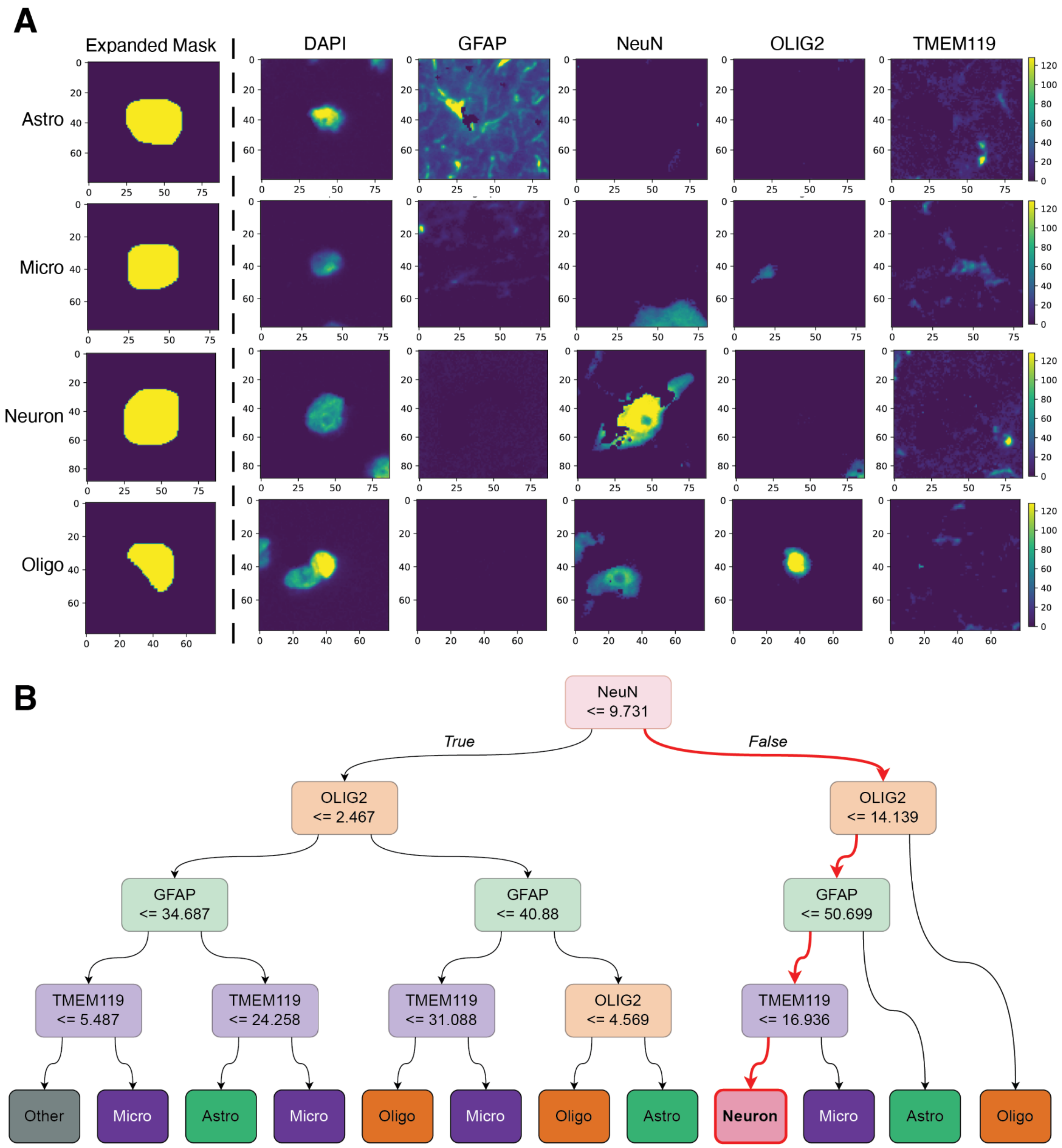


**Supplementary Figure 34. Decision-tree framework for CART-classification of cell types using Visium-SPG immunofluorescence intensities.** (**A**) Fluorescence intensities for randomly chosen cells of each CART-classified type from Br2720_Ant_IF. For each segmented nucleus, masks are slightly dilated to encompass some surrounding cell cytoplasm (Supplemental Methods: Image segmentation and processing), shown in the leftmost column. Fluorescence, taking values between 0 and 127, for the DAPI channel and other channels marking each cell type is shown in the remaining columns. The decision tree uses the mean of the values overlapping the expanded mask for classification. CART classifications are used as one means of benchmarking the accuracy of spot-deconvolution software (**Fig. 4D**, **Fig S35**, **Fig S36**) (**B**) The decision-tree structure, which is traversed downwards. Each colored box represents a decision, made by comparing the mean fluorescence of a single channel for a given segmented cell against a learned threshold (indicated within each box). Red arrows highlight the series of decisions made to classify an example neuron: mean NeuN fluorescence must exceed 9.731, OLIG2 fluorescence must be less than or equal to 14.139, GFAP fluorescence must be less than or equal to 50.699, and TMEM119 fluorescence must be less than or equal to 16.936.

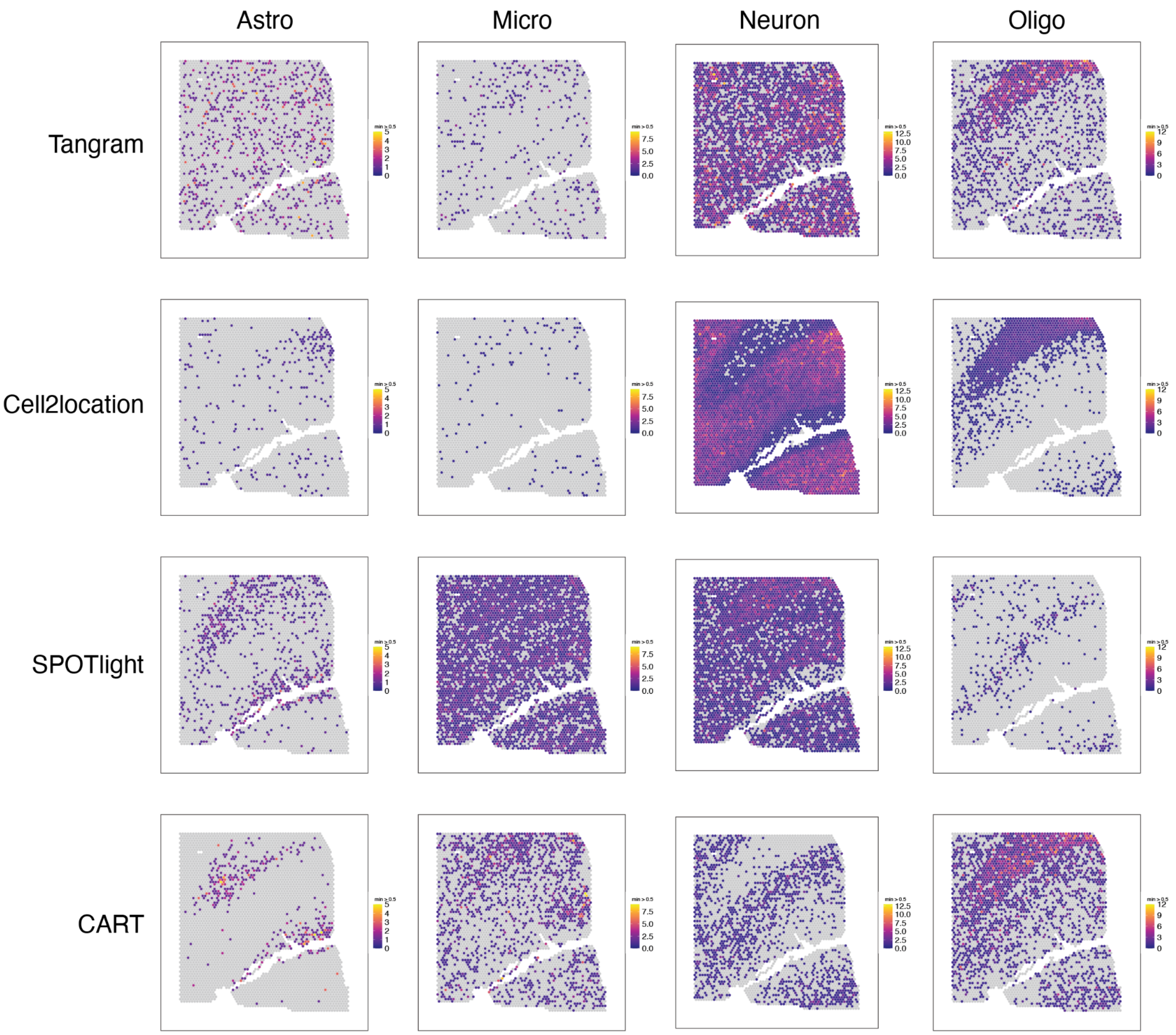


**Supplementary Figure 35. Spatial comparison of cell type counts by spot deconvolution method for Br6522_Ant_IF.** The top three rows are spotplots showing layer-level cell type counts for each Visium-SPG sample, as estimated by *Tangram*, *Cell2location*, and *SPOTlight*. Cell type counts were collapsed onto the resolution used by the classification and regression tree (CART) classifier (i.e. Astro, Micro, Neuron, Oligo), which was determined by the 4-plex immunofluorescent labeling. Spotplots of CART-predictions are in the bottom row.

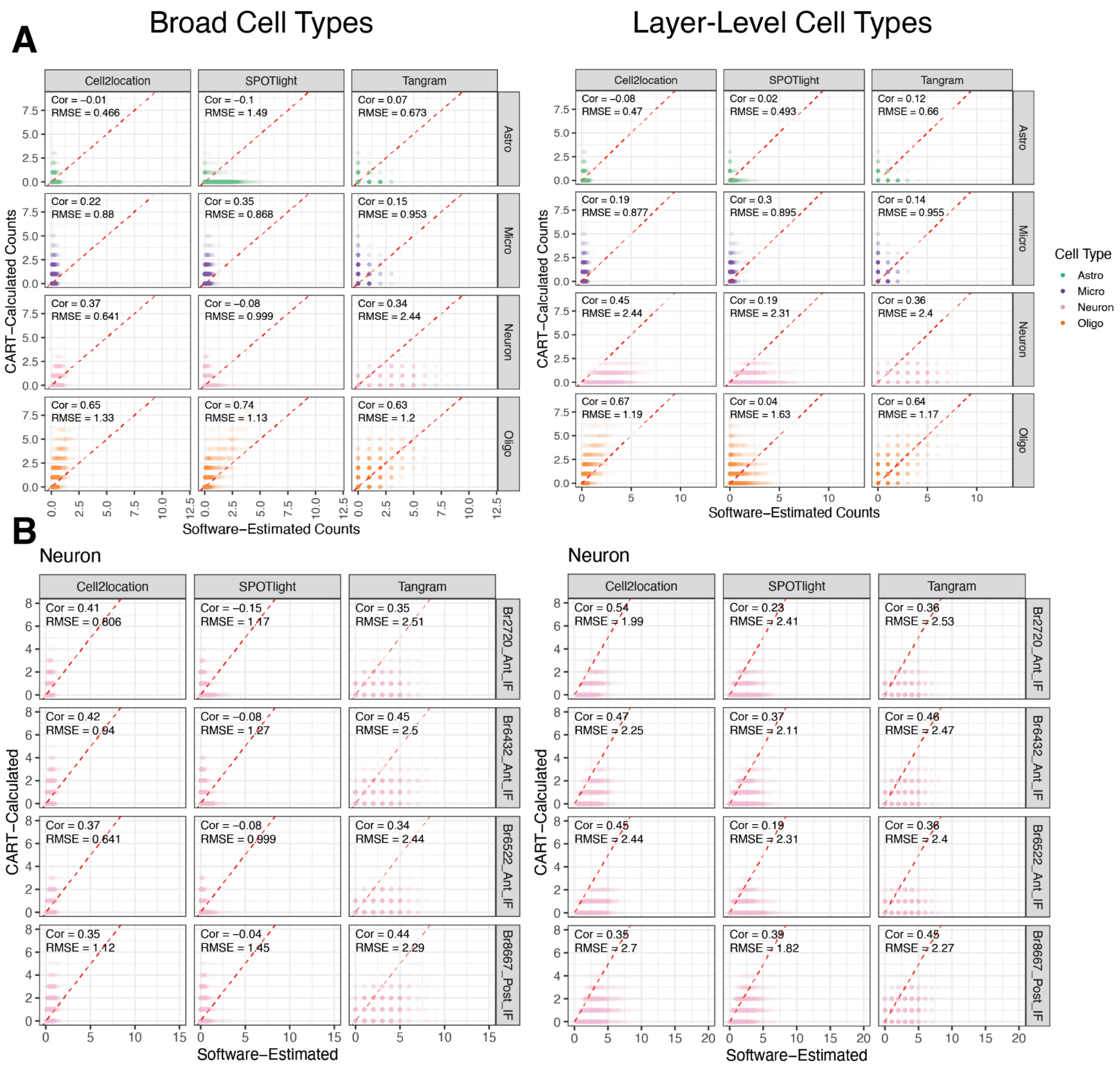


**Supplementary Figure 36. Spot-level benchmark of spot deconvolution results.** (**A**) CART-calculated vs. software-estimated counts for 4 immunolabeled cell types (Astro, Micro, Neuron, Oligo) at broad and layer-level resolutions for Br6522_Ant_IF. Each point represents the number of cells of the given cell type present in a particular spot, both as estimated by each software method and as determined by the decision-tree classifier using the IF image as input. Pearson correlation and root mean squared error (RMSE) are used as metrics to summarize the relationship between CART-calculated and software-estimated counts for a particular software method and cell type. (**B**) CART-calculated vs. software-estimated counts for neurons in all four Visium-SPG sections at broad and layer-level cell type resolutions. Correlation and RMSE values are also calculated for the other cell types, forming a more comprehensive assessment of concordance between CART and software estimates (**Fig. 4D**).

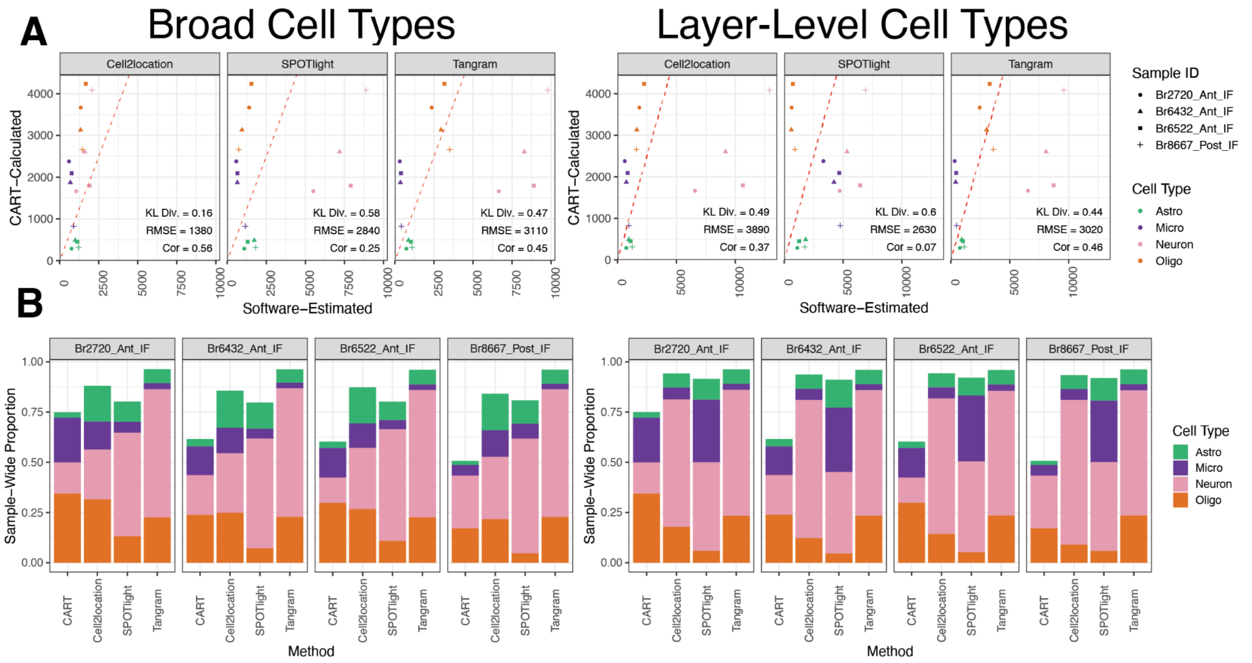


**Supplementary Figure 37**. **Predicted section-wide cell type proportions for Visium-SPG samples at broad and layer-level resolution**. (**A**) Predicted proportions from each software, including *Tangram*, *Cell2location*, and *SPOTlight* were compared to the CART-predicted proportions of each sample to assess general concordance of cell type composition estimates. Cell types at broad and layer-level resolution were collapsed to the 4 immunolabeled cell types quantifiable by CART. Kulback-Leibler divergence, RMSE, and correlation were computed to quantify performance of each of the three methods (Supplemental Methods: Evaluating performance of spot-deconvolution methods) (**B**) The information in (**A**) presented as a barplot. Totals do not add to 1 because CART can classify cells as “other” (i.e. negative for Astro, Micro, Neuron, and Oligo likely representing endothelial and mural cells), which were input as an “endomural” cell type to the three deconvolution methods.

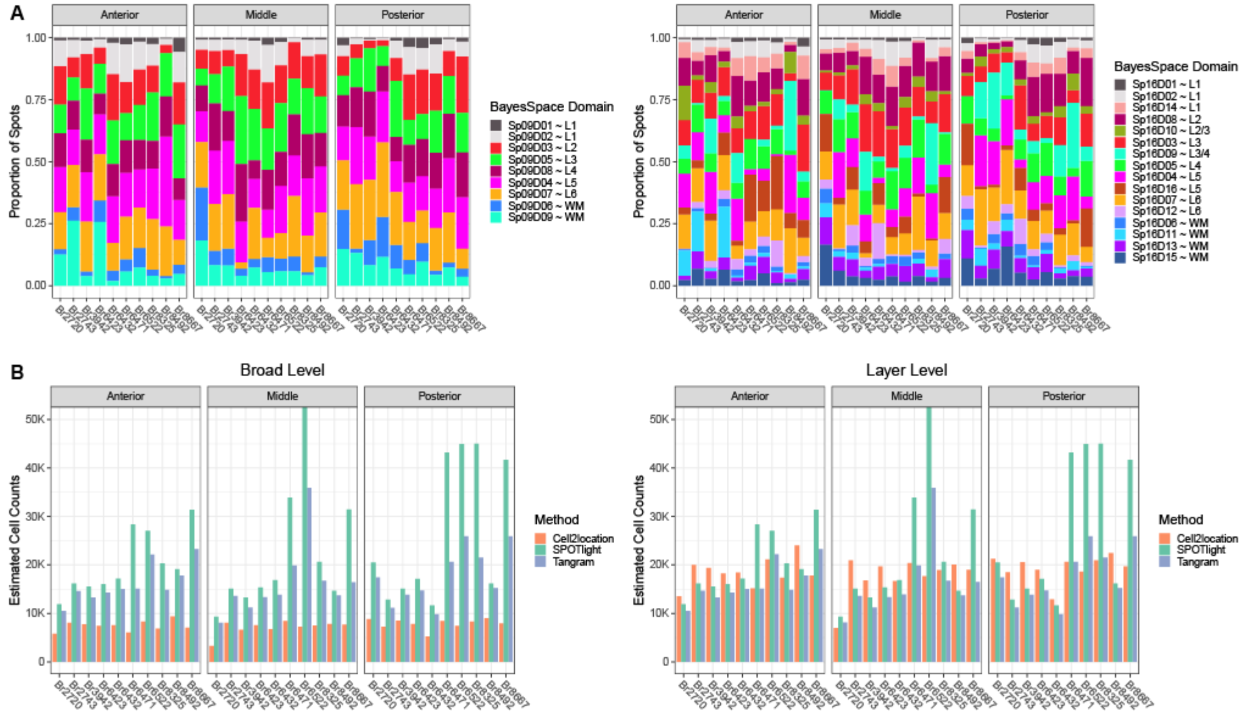


**Supplementary Figure 38**. **Composition summaries across spatial domains (SpDs) and spot deconvolution results.** (**A**) Proportion of spots assigned to Sp$_9$Ds and Sp$_{16}$Ds across all 30 samples separated by position (anterior, middle, posterior). (**B**) Total number of predicted cells resulting from deconvolution with *Tangram*, *Cell2location*, or *SPOTlight* softwares across all 30 samples separated by position. The left side shows the broad cell type level estimates and the right side shows the layer-level estimates. As noted in **Fig S31**, the number of cells outputted from *SPOTlight* matches the number of cells used as input (based on *Vistoseg* segmentations), whereas *Cell2location* and *Tangram* can have mismatches between input and output cell numbers. As highlighted in **Fig. 4E**, the number of cells predicted by *Tangram* is the most consistent across resolutions.

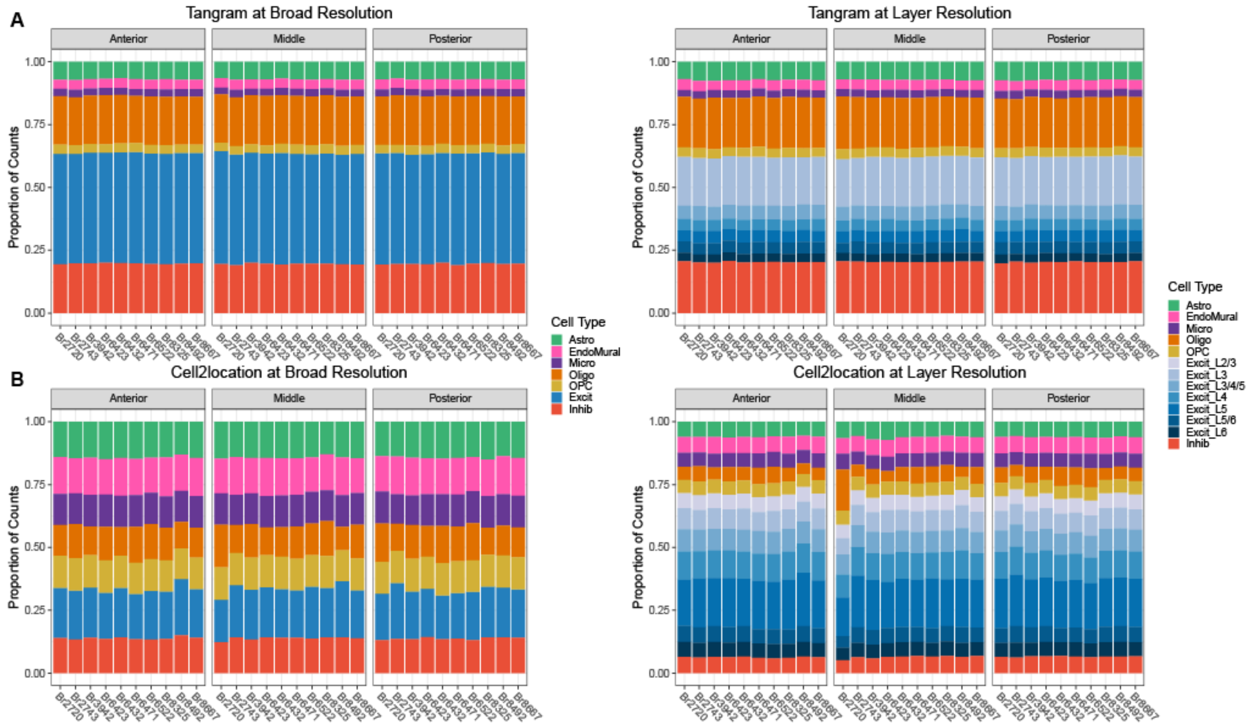


**Supplementary Figure 39**. **Spot deconvolution results for *Tangram* and *Cell2location* across all Visium samples.** (**A**) *Tangram* and (**B**) *Cell2location* predicted cell compositions across all Visium samples separated by position (anterior, middle, posterior). Consistent with **Fig. 4E**, *Tangram* shows similar cell type proportions at broad and layer-level resolutions. Both methods show consistent cell type proportions across samples and DLPFC position.

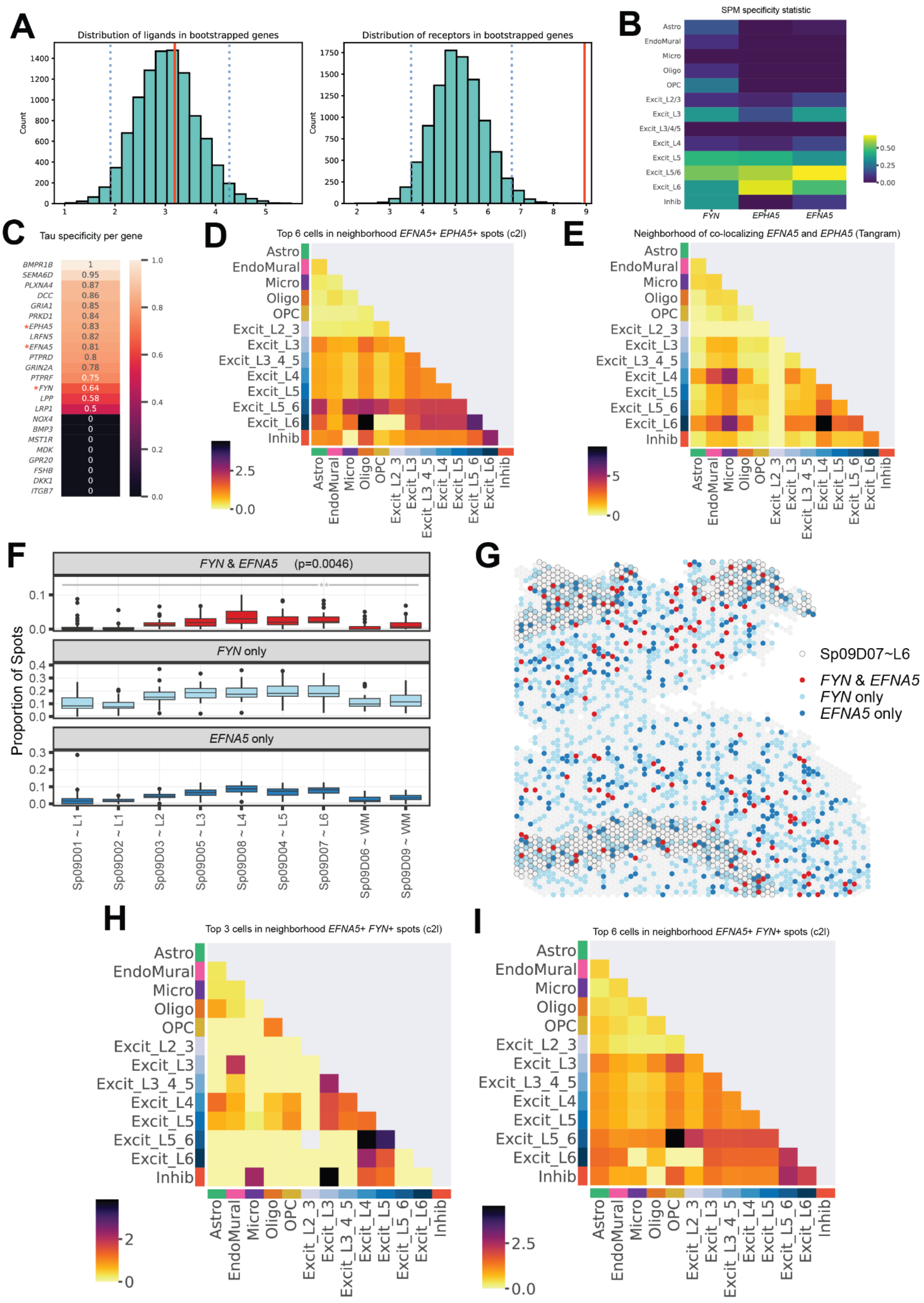
A
Distribution of ligands in bootstrapped genes
Distribution of receptors in bootstrapped genes
Count
B
SPM specificity statistic
Astro
EndoMural
Micro
Oligo
OPC
Excit_L2/3
Excit_L3
Excit_L3/4/5
Excit_L4
Excit_L5
Excit_L5/6
Excit_L6
Inhib
FYN
EPHA5
EFNA5
C
Tau specificity per gene
BMPR1B 1
SEMA6D 0.95
PLXNA4 0.87
DCC 0.86
GRIA1 0.85
PRKD1 0.84
*EPHA5 0.83
LRFN5 0.82
*EFNA5 0.81
PTPRD 0.8
GRIN2A 0.78
PTPRF 0.75
*FYN 0.64
LPP 0.58
LRP1 0.5
NOX4 0
BMP3 0
MST1R 0
MDK 0
GPR20 0
FSHB 0
DKK1 0
ITGB7 0
D
Top 6 cells in neighborhood EFNA5+ EPHA5+ spots (c2l)
E
Neighborhood of co-localizing EFNA5 and EPHA5 (Tangram)
Astro
EndoMural
Micro
Oligo
OPC
Excit_L2_3
Excit_L3
Excit_L3_4_5
Excit_L4
Excit_L5
Excit_L5_6
Excit_L6
Inhib
F
FYN & EFNA5 (p=0.0046)
FYN only
EFNA5 only
Proportion of Spots
Sp09D01 ~ L1
Sp09D02 ~ L1
Sp09D03 ~ L2
Sp09D05 ~ L3
Sp09D08 ~ L4
Sp09D04 ~ L5
Sp09D07 ~ L6
Sp09D06 ~ WM
Sp09D09 ~ WM
G
Sp09D07~L6
FYN & EFNA5
FYN only
EFNA5 only
H
Top 3 cells in neighborhood EFNA5+ FYN+ spots (c2l)
I
Top 6 cells in neighborhood EFNA5+ FYN+ spots (c2l)

**Supplementary Figure 40**. **Ligand-receptor analysis performed at spatial resolution in context of genetic risk for schizophrenia (SCZ)** (**A**) Ligand and receptor occurrence in SCZ risk gene list compared to bootstrapped dataset of same size. (**B**) SPM specificity statistics calculated using snRNA-seq data for genes of interest for each cell type. (**C**) Tau specificity statistic for SCZ risk genes of interest, where "*" indicates genes involved in the LR interaction also identified with data-driven cell-cell communication tool *LIANA (12)* (**D**) Cell type neighborhood colocalization analysis using top 6 *Cell2location (c2l)*-predicted cell types per Visium spot in *EFNA5* and *EPHA5* co-expressing spots (**E**) Colocalization analysis using *Tangram*-predicted cell types in *EFNA5* and *EPHA5* co-expressing spots (**F**) The proportion of spots co-expressing *EFNA5* an *FYN* across data-driven $SpD_9$s. Consistent with **Fig. 5E**, spots co-expressing FYN and *EFNA5*-are enriched in $Sp_9D_7$. (**G**) Spatial mapping of *FYN*-*EFNA5* co-expressing spots in relation to $Sp_9D_7$~L6. (**H**) Cell type neighborhood colocalization analysis for top 3 *c2l*-predicted cells in *EFNA5* and *FYN* co-expressingspots. (**I**) Cell type neighborhood colocalization analysis for top 6 *c2l*-predicted cells in *EFNA5* and *FYN* co-expressing spots.

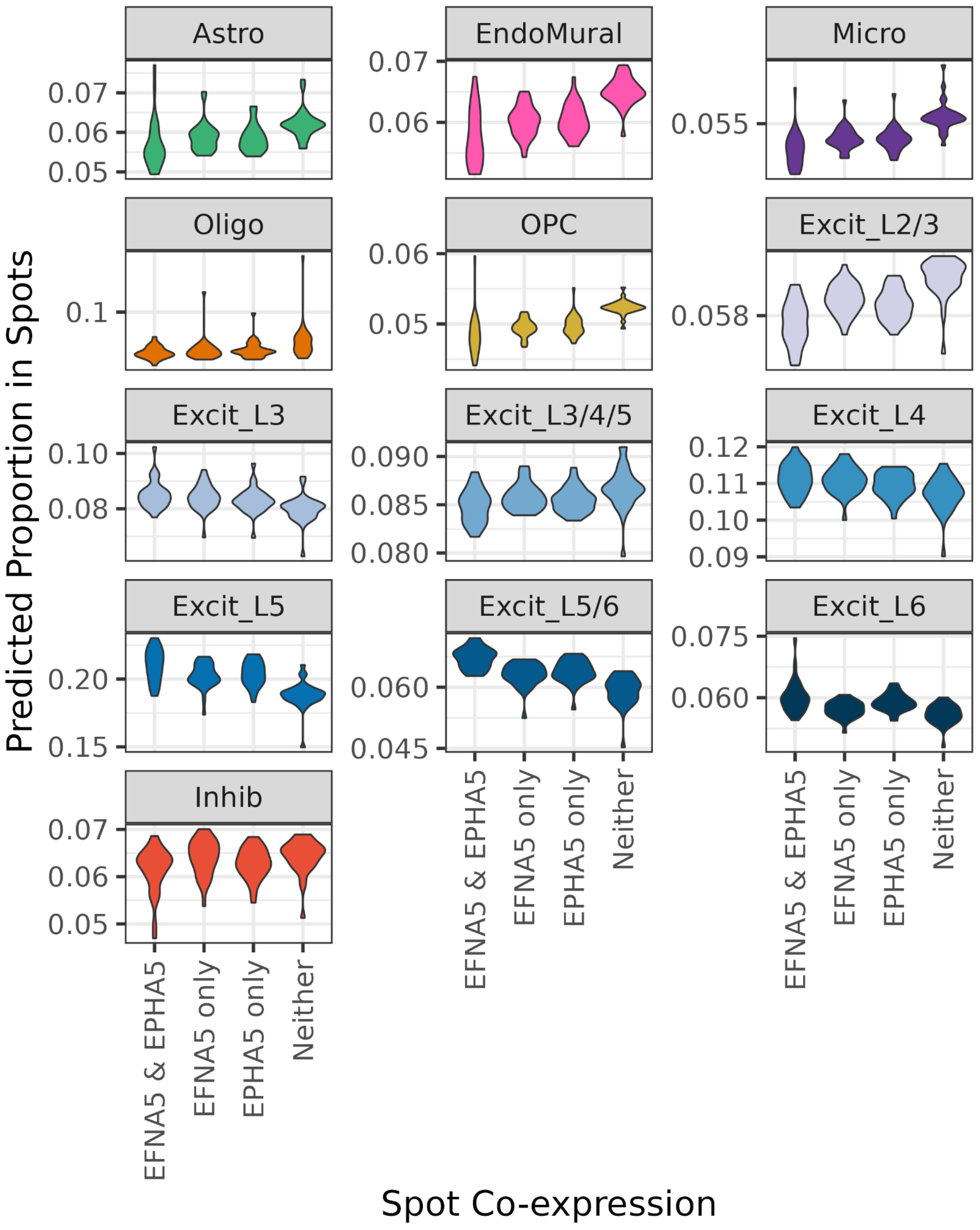


**Supplementary Figure 41. Distributions of predicted cell proportions in *EFNA5*, *EPHA5* and *EFNA5-EPHA5* co-expressing spots for all Visium samples.** Spots co-expressing *EFNA5* and *EPHA5* show a higher *Cell2location*-predicted proportion of Excit_L5/6 neurons (p=1.8e-12) and Excit_L6 neurons (p=3.0e-4) compared to spots where they are not co-expressed. This is also observed for Excit_L3 (p=5.9e-3), Excit_L4 (p=9.0e-3) and Excit_L5 (p=9.9e-6) neurons, but not for other excitatory neuron (Excit_L2/3 p=1.00), inhibitory neuron (p=0.99), or non-neuronal populations (all p>0.99).

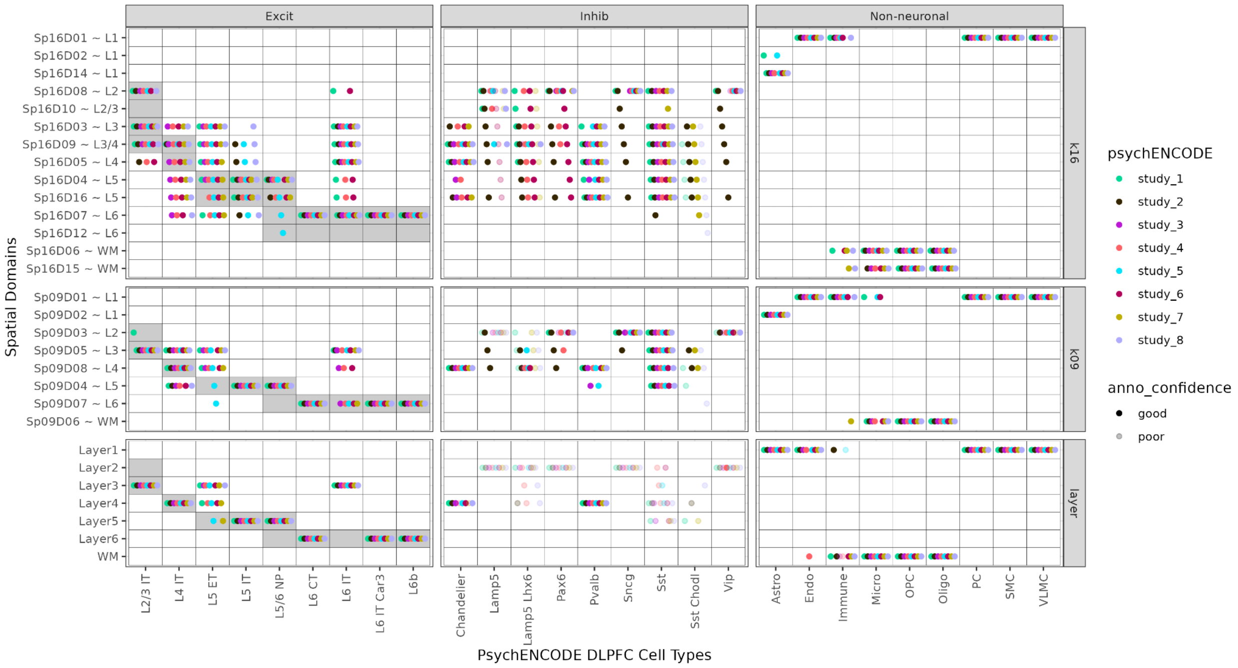


**Supplementary Figure 42. Spatial registration of full PsychENCODE snRNA-seq dataset (cases and controls).** Summary of spatial registration for PsychENCODE snRNA-seq DLPFC datasets, including case and control samples from a variety of neurodevelopmental and neuropsychiatric disorders. Registration was performed against manually annotated histological layers (6) as well as unsupervised *BayesSpace* clusters at $k$=9 and $k$=16 ($Sp_9Ds$ and $Sp_{16}Ds$, respectively). Each dot shows the histological layer(s) or $Sp_kD$(s) where a dataset's cell type was annotated during spatial registration. Solid dots show good confidence in the spatial annotation ($\rho > 0.25$), while translucent dots show poor confidence ($\rho \leq 0.25$) in the annotation.
